# Phylogenomics reveals the deep ocean as an accelerator for evolutionary diversification in anglerfishes

**DOI:** 10.1101/2023.10.26.564281

**Authors:** Elizabeth Christina Miller, Rose Faucher, Pamela B. Hart, Melissa Rincon-Sandoval, Aintzane Santaquiteria, William T. White, Carole C. Baldwin, Masaki Miya, Ricardo Betancur-R, Luke Tornabene, Kory Evans, Dahiana Arcila

## Abstract

Colonization of a novel habitat is often followed by radiation in the wake of ecological opportunity. Alternatively, some habitats should be inherently more constraining than others if the challenges of that environment have few evolutionary solutions. We examined the push-and-pull of these factors on evolution following habitat transitions, using anglerfishes (Lophiiformes) as a model. Deep-sea fishes are notoriously difficult to study, and poor sampling has limited progress thus far. Here we present a new phylogeny of anglerfishes with unprecedented taxonomic sampling (1,092 loci and 40% of species), combined with three-dimensional phenotypic data from museum specimens obtained with micro-CT scanning. We use these datasets to examine the tempo and mode of phenotypic and lineage diversification using phylogenetic comparative methods, comparing lineages in shallow and deep benthic versus bathypelagic habitats. Our results show that anglerfishes represent a surprising case where the bathypelagic lineage has greater taxonomic and phenotypic diversity than coastal benthic relatives. This defies expectations based on ecological principles since the bathypelagic zone is the most homogeneous habitat on Earth. Deep-sea anglerfishes experienced rapid lineage diversification concomitant with colonization of the bathypelagic zone from a continental slope ancestor. They display the highest body, skull and jaw shape disparity across lophiiforms. In contrast, reef-associated taxa show strong constraints on shape and low evolutionary rates, contradicting patterns suggested by other shallow marine fishes. We found that Lophiiformes as a whole evolved under an early burst model with subclades occupying distinct body shapes. We further discuss to what extent the bathypelagic clade is a secondary adaptive radiation, or if its diversity can be explained by non-adaptive processes.

## INTRODUCTION

How does evolution proceed after the colonization of novel but harsh environments? The bathypelagic zone of the deep sea (>1,000 m) is characterized by a lack of solar light, food limitation, high pressure, low temperatures, and large expanses of homogeneous space^1–4^. Fishes living at this depth converged on specializations including large jaws and teeth, reduced metabolic rate, reduced musculature and skeletal density, sensitive eyes, and bioluminescence^1,2,5–13^. The repeated evolution of these adaptations across distantly related lineages may be an indication that there are a limited number of potential solutions to overcome the challenges of this environment^14^. In contrast to the deep sea, coastal marine environments such as coral reefs and estuaries are diverse, productive and topologically complex^15,16^. Due to their sharper biotic and abiotic clines, and presumably greater number of niches, we should expect coastal habitats to promote ecological, morphological, and lineage diversification relative to open ocean or deep sea settings^17–24^. Yet, recent studies using phylogenetic comparative methods have shown that fishes from the latter habitats can have greater phenotypic diversification rates and disparity in body shape^25–29^. The reasons for this remain unclear, but nonetheless contradict expectations based on first principles^30^.

The order Lophiiformes is an iconic clade of marine fishes whose members are characterized by a lure on their head that is used for sit-and-wait hunting. Lophiiformes contains ∼350 species among five well-supported suborders: Lophioidei (monkfishes), Ogcocephaloidei (hand batfishes), Antennarioidei (frogfishes), Chaunacoidei (sea toads), and Ceratioidei (dreamers and sea devils)^31^. Four of the five suborders are benthic and occupy the continental shelf, slope and rise, while the ceratioids are bathypelagic. The ceratioids are known for their extreme sexual size dimorphism and varying degrees of sexual parasitism in which males fuse to a female, a phenomenon not found in any other vertebrate^32^. In addition to their habitat diversity, anglerfishes also exhibit diverse body shapes ranging from laterally compressed, dorsoventrally compressed, globose, and elongated. Specializations of benthic lophiiforms include extreme oral gape expansion^33^, a tetrapod-like walking gait^34^, and extremely slow breathing in low-oxygen settings^35,36^. It is believed that their shape diversity is related to the evolution of restricted gill openings, which frees constraints on cranial morphology^37^ and allows the body to fill with water to perform these specialized functions.

How have habitat transitions shaped the evolution of anglerfishes? First, we hypothesize that shallow and/or benthic species will have faster rates of phenotypic and lineage diversification than bathypelagic anglerfishes. Even deep benthic environments are more heterogeneous than the deep pelagic zone^3,38,39^, and substrate preferences are evident from videos of deep benthic chaunacids and lophiids^40–42^. In contrast, the homogeneity of the bathypelagic zone is unparalleled on Earth^2^. There are few barriers to dispersal which should limit speciation^43–45^ (but see ^46–49^). Further, the environmental challenges in the deep pelagic zone should impose constraints on evolution, limiting the number of viable phenotypes^14^ and thereby reducing rates of phenotypic evolution^50^. Phenotypic constraints associated with a particular habitat can be detected using a model-fitting approach, with an Ornstein-Uhlenbeck (OU) model being most consistent with this type of constraint (Table 1).

**Table 1.** Summary of hypotheses and predictions.

| Hypothesis | Description | Mechanisms | Predictions |  |  |
| --- | --- | --- | --- | --- | --- |
|  |  |  | Evolutionary mode | Phenotypic Disparity | Evolutionary rates |
| 1 | Benthic habitats promote evolution while the bathypelagic zone constrains evolution | More niches and opportunities for allopatry in benthic habitats; harsh conditions with few evolutionary solutions, and few barriers to dispersal, in the bathypelagic | Ceratioid evolution best described by bounded (OU) models | Shallow-water and/or benthic suborders with greater morphological disparity than ceratioids | Ceratioids with slower rates of evolution and lineage diversification than benthic suborders |
| 2a | Bathypelagic zone promotes evolution relative to benthic habitats; ceratioid diversification is ongoing | Ceratioids have not exhausted ecological opportunity in the bathypelagic zone, or phenotypic change is non-adaptive as well as adaptive | Ceratioid evolution best described by unbounded (BM) models | Ceratioids with greater morphological disparity than shallow-water and/or benthic suborders | Ceratioids with faster rates of evolution and lineage diversification than benthic suborders; rates do not slow through time |
| 2b | Bathypelagic zone promotes evolution relative to benthic habitats; ceratioid diversification has slowed down | Ceratioids have exhausted ecological opportunity in the bathypelagic zone | Ceratioid evolution best described by bounded (EB) models | Ceratioids with greater morphological disparity than shallow-water and/or benthic suborders | Ceratioids with faster rates of evolution and lineage diversification than benthic suborders; rates slow through time |

Alternatively, we hypothesize that the bathypelagic anglerfishes could have faster rates of diversification and be less evolutionarily constrained than shallow-water or deep-benthic relatives. Specifically, due to the lack of solar light, predator-prey interactions occur over short spatial scales in the deep sea, often facilitated by bioluminescence^2,9,51^. This presumably reduces selection for the fusiform body shapes common among shallow-water pelagic fishes^22,23,26,29,52–55^, allowing ceratioids to explore new areas of morphospace. If this ecological release is associated with an increase in phenotypic diversity, speciation, and the filling of novel ecological niches^56,57^, then ceratioids would fit the search image of an adaptive radiation incited by the colonization of a novel habitat^58–60^. If this hypothesis is supported, we would expect ceratioid morphological disparity to be higher than that of benthic relatives.

We can further divide this latter hypothesis into two sub-hypotheses, distinguishable by the mode of evolution (Table 1). First, phenotypes in ceratioids may be continuously diversifying over time. This could occur if the radiation is still in its early stages, if ecological opportunity has not been exhausted, or if phenotypic diversity accumulates via non-adaptive processes such as genetic drift in addition to adaptive evolution. In this case, we would expect phenotypes to be evolving under an unbounded Brownian motion (BM) model of evolution. Alternatively, we may expect to see a slowdown in phenotypic evolution in ceratioids following their initial radiation from the benthos. This could indicate that the radiation is in its late stages, that competition for similar resources prevents lineages from overlapping in morphology, or that there are few ecological niches in the bathypelagic zone to begin with. Under this sub-hypothesis, ceratioid phenotypes would be evolving under an “early burst” (EB) model, in which phenotypic and lineage diversification is fastest early in a clade’s history as subclades occupy new adaptive zones free from negative ecological interactions, but slows with time as diversification proceeds within these adaptive zones. Unlike BM, the EB model enforces a constraint on phenotypic evolution; unlike OU models, the constraint is time-dependent^61^. While some authors associate the EB model with diagnosing adaptive radiation sensu Simpson^61,62^ (i.e., process-based definition), we prefer a broader definition of adaptive radiation as a lineage that has evolved taxonomic and phenotypic diversity associated with different ecologies^20,59,63,64^ (i.e., outcome-based definition). The EB model might therefore be interpreted as an “ecological limits” model instead of an adaptive radiation model.

Sampling of deep-sea fishes for phylogenetic analysis is stymied by the difficulty of collecting^3,65,66^. Dense species sampling is needed to gain power for phylogenetic comparative methods^67^, ultimately limiting what we can learn about the evolution of deep-sea fishes. Here we present a novel phylogenomic hypothesis of anglerfishes (Lophiiformes) based on 1,092 single-copy exon markers. Due to contributions from many natural history collections and government agencies^68,69^, our taxonomic sampling greatly improves upon predecessors^70–72^, with nearly 40% of species and all deep-sea families sampled. This advance allowed us to apply phylogenetic comparative methods largely reserved for well-sampled terrestrial and shallow-water organisms to test hypotheses about evolution in the deep sea.

## RESULTS

### Phylogenomic inference and divergence times

We generated new genomic data for 152 lophiiform individuals from 120 species using exon capture approaches proven successful for fishes^55,73–75^ (Table S1). Sampling was augmented by mining exons from published UCEs^71,72^ and legacy markers from NCBI (Tables S2, S3). Final taxonomic sampling after quality control included 132 species of Lophiiformes (37.8% of species) and 20 of 21 families (all but Lophichthyidae). Sampling of ceratioids included all 11 families and 32.1% of species. Relationships were largely in agreement between concatenation- and coalescent-based phylogenomic analyses (Appendix A1). These relationships strongly suggest that obligate sexual parasitism (found in Ceratiidae, Neoceratiidae, and Linophrynidae) evolved more than once^32,70^. Detailed systematic results are given in Appendix A2.

We assembled a set of 21 node calibrations, including eight outgroup and ten ingroup fossils and three geologic calibrations (Appendix A3). Our calibration scheme is novel and includes six lophiiform fossils from the Eocene Monte Bolca communities^76^ (Fig. 1). To incorporate uncertainty in topology and divergence times for comparative analyses, we produced eight alternative time trees using either the IQ-TREE or ASTRAL tree, the calibration scheme with or without the controversial fossil †*Plectocretacicus*^75,77^, and using either MCMCtree^78,79^ or RelTime^80,81^ as the calibration method. The methodological choice with the largest impact on divergence times was MCMCTree versus RelTime (Fig. 1, Appendix A4). For this reason, some comparative analyses involving complex visualizations were repeated on two designated “master” trees: the IQ-TREE calibrated with the scheme including †Plectocretacoidea using either MCMCTree or RelTime (hereafter “master MCMCTree” or “master RelTime tree”).

**Figure 1:**
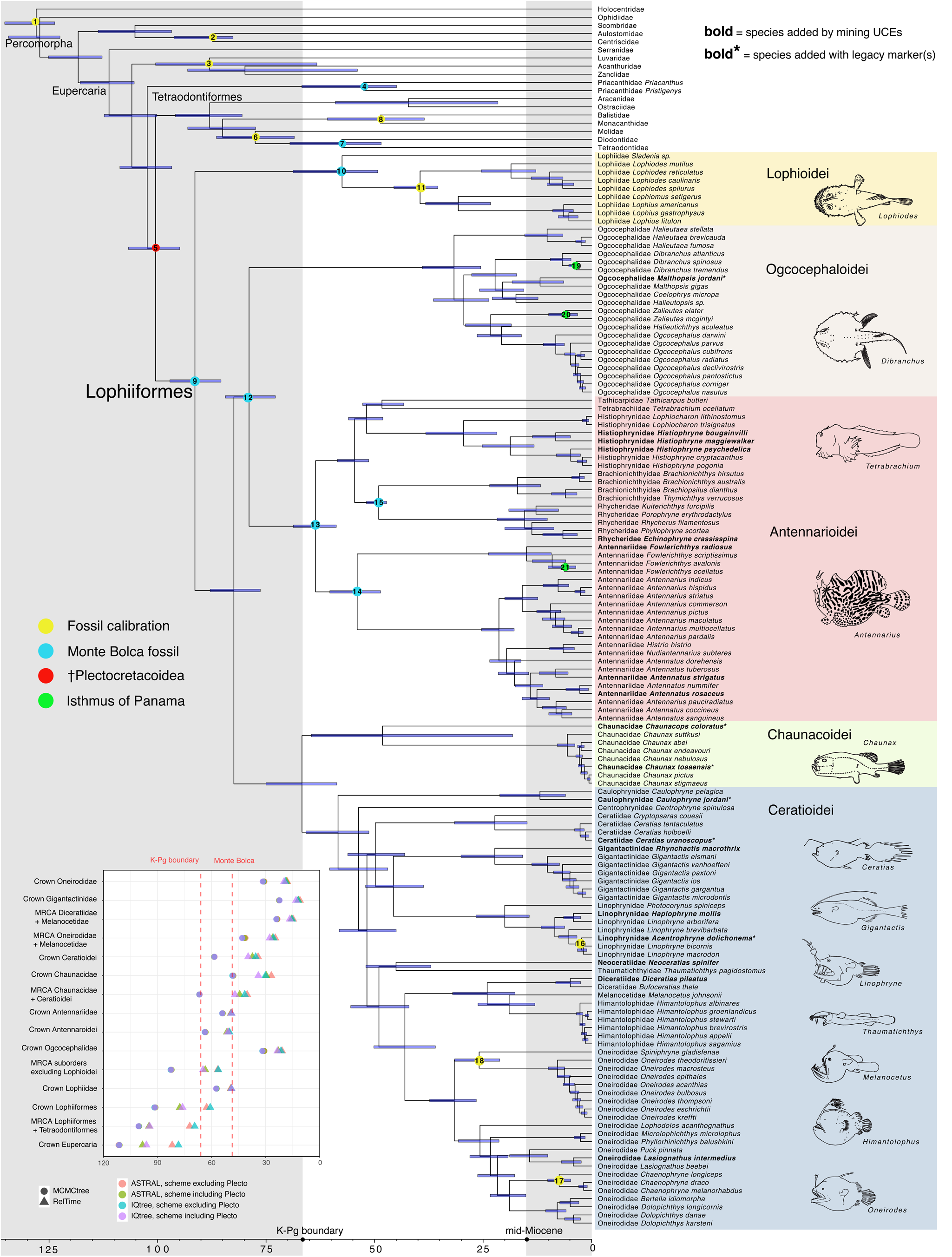
Time-calibrated phylogeny of Lophiiformes. Inset shows the range of dates for key nodes inferred across the eight alternative time trees. This tree was inferred using IQ-TREE and calibrated using MCMCTree with the scheme including †Plectocretacoidea (master MCMCTree); for the master RelTime tree see Appendix A1. Grey shading indicates the Cretaceous and the mid-Miocene (∼15 Ma) to present, the latter period identified as having elevated rates of speciation across deep-sea fishes^49^. Line art was digitized from FAO fisheries guides.

Six out of eight time trees inferred a Cretaceous origin of crown Lophiiformes (92–61 Ma across trees) (Fig. 1). In the MCMCTrees, Ceratioidei split from Chauancoidei near the K/Pg boundary (67 Ma), whereas in the RelTime trees this divergence occurred in the Eocene (47–40 Ma). Similarly, the two methods result in a >20 million-year difference in the age of crown Ceratioidei, either in the Paleocene (∼58 Ma using MCMCTree) or late Eocene (40–34 Ma using RelTime). Detailed discussion of divergence times is given in Appendix A4.

### Habitat transitions

Ancestral habitat reconstructions (Table S4) based on the best-fitting biogeographic model (BAYAREA+J; Table S5) indicated that the MRCA of all Lophiiformes had a widespread depth range spanning the continental shelf and slope^82^ (Fig. 2A). The bathypelagic ceratioids originated from a benthic continental slope ancestor. In other words, the most significant habitat transition associated with the ceratioids was benthic-to-pelagic, not shallow-to-deep. There were two independent transitions to a shallow-only habitat associated with frogfishes (Antennarioidei) and the hand batfish genus *Ogcocephalus*.

**Figure 2:**
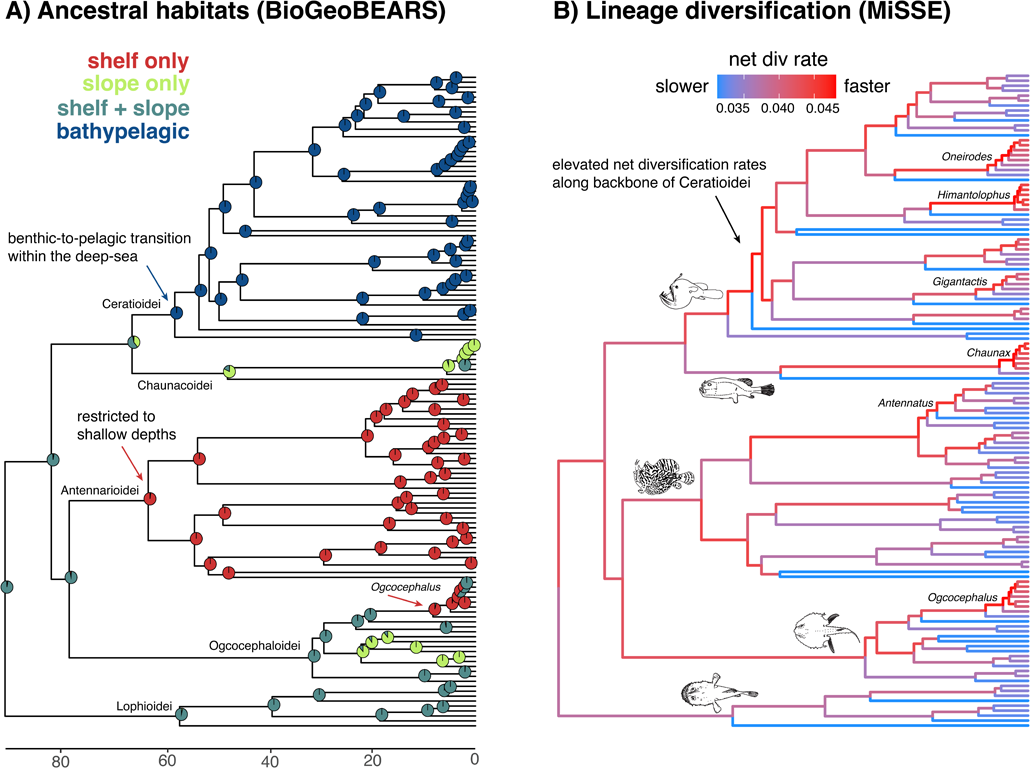
Timing of habitat transitions and lineage diversification rates. (A) Habitat reconstructions inferred using BioGeoBEARS. (B) Branch-specific net diversification rates inferred using MiSSE. For tip-associated rates across all trees see Fig. S1. Here the master MCMCTree is shown; for comparison with the master RelTime tree see Fig. S2.

### Lineage diversification rates

We estimated branch-specific net diversification rates using the MiSSE framework (missing state speciation and extinction)^83^. MiSSE models with 1–7 rate classes were supported with >5% of the relative Akaike weight across the alternative trees (Table S6). There was little consensus on the best-fit model for any tree, therefore we model-averaged rates^84^. The backbone of Ceratioidei had elevated net diversification rates following the benthic-to-pelagic transition at the base of the clade (Fig. 2B, Fig. S2). The distributions of recent (tip-associated) rates of net diversification overlapped among suborders and habitats (Fig. S1). Five genera had particularly high net diversification rates: the deep benthic *Chaunax*, the ceratioids *Gigantactis, Oneirodes,* and *Himantolophus,* and the shallow-water batfishes *Ogcocephalus* (Fig. 2B). Rates were higher overall in the RelTime trees compared to the MCMCTrees due to the generally shorter branch lengths of the former (Figs. S1, S2). Pruning for suspected taxonomic inflation in certain genera (Appendix A2) reduced rate variation overall, but the general patterns remained (Fig. 1).

### Phenotypic disparity

Phylomorphospace analyses^85^ showed that the five lophiiform suborders generally occupied distinct regions of morphospace associated with different body plans (Fig. 3, Fig. S3). The first principal component (PC1) explained 45.0% of the variation in body shape. Taxa with laterally compressed bodies and small eyes had negative values, while dorsoventrally compressed, large-eyed taxa had positive values (Fig. 3A). The second PC axis explained 21.3% of the variation and corresponded to body elongation, mouth width, and jaw length, with short bodies and small mouths having low values and elongate bodies and large mouths having high values. By habitat, the body shape of female ceratioids were generally restricted to low values of PC1 and high values of PC2. Benthic species found on the continental slope were restricted to high values of PC1 but were distributed throughout PC2. Both continental shelf clades (Antennarioidei and *Ogcocephalus*) were restricted to low values of PC2. Thus, the transition from deep benthic to deep pelagic habitats incurred a relative increase in jaw size and decrease in eye size. Shallow-water species generally exhibit more truncated bodies and mouths compared to deep-sea species.

**Figure 3:**
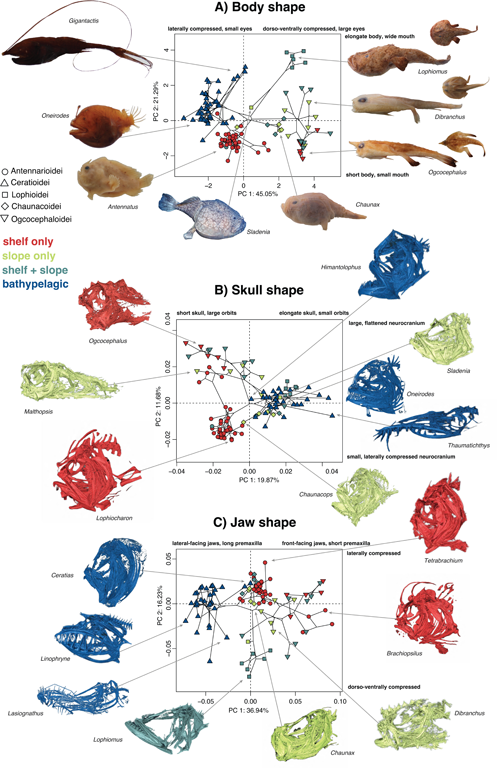
Phylomorphospace analyses of (A) body shape, (B) skull shape and (C) jaw shape. Body shape was inferred from ten linear measurements (Fig. S3). Skull and jaw shapes were inferred using geometric morphometrics from CT scans (Fig. S4). *Sladenia* image from NOAA.

Morphospace analyses based on micro-CT scans of skulls (Fig. S4, Table S7) showed greater overlap in shapes among suborders compared to analyses based on body shape (Fig. 3B). The first PC axis explained 19.9% of skull shape variation and was related to elongation of the skull and the relative size and position of the jaws and orbit (with *Ogcocephalus* having the lowest values and *Thaumatichthys* having the highest values). The second PC axis explained 11.7% of the variation and was generally related to size and compression of the neurocranium (with *Lophiocharon* having the smallest values and *Ogcocephalus* having the highest values). We found a strong split in skull shape morphospace by habitat, with all continental shelf taxa exhibiting negative values along PC1 while the bathypelagic taxa exhibit positive values along this axis. Continental shelf habitats are generally associated with shorter and narrower skulls with the orbit positioned high on the head. Deep benthic taxa were widely distributed in morphospace.

Convergence emerged as a theme in jaw shape morphospace (Fig. 3C). The first PC axis explained 37.0% of the variance, with positive values corresponding to foreshortened, front-facing jaws with truncate premaxillae relative to the dentaries (e.g. *Brachionichthys*) and negative values corresponding to more laterally-positioned jaws and elongate premaxillae relative to the dentaries (e.g. *Linophryne*). The second PC axis explained 16.2% of the variance and corresponded to lateral versus dorsoventral compression of the jaws (with *Lophiomus* having the most negative values and *Tetrabrachium* having the most positive values). Ceratioids were nearly all restricted to negative values of PC1 with exception of Ceratiidae, whose jaws more closely resembled chauancids and shallow-water antennarioids. Similarly, the antennarioid brachionichthyids (handfishes) converged with batfishes in jaw shape. By habitat, continental shelf taxa tended towards average or high values of PC1 and PC2, while deep benthic taxa were widely distributed across the morphospace.

We quantified shape disparity^86^ for suborder (Table S8) and habitat categories (Table S9). Across the three phenotypic datasets, the bathypelagic ceratioids had the greatest disparity accounting for 37–41% of the total disparity of Lophiiformes (Fig. S5). The remaining habitats each accounted for less disparity: shelf only (22–31%), shelf and slope (22–27%) and slope only (6–11%). Disparity among the remaining suborders was distributed as: Ogcocephaloidei (23– 30%), Antennarioidei (13–25%), Lophioidei (9–12%), and Chaunacoidei (4–6%). Note that while the four benthic suborders individually contain less disparity than ceratioids, when combined they account for 59–63% of the disparity of Lophiiformes, meaning the benthic state in general contains more disparity than the pelagic state.

### Tempo and mode of phenotypic evolution

We used an evolutionary model fitting approach to identify the mode of body, skull and jaw shape evolution for Lophiiformes as a whole and within each suborder individually. Multivariate model-fitting analyses performed using mvMORPH^87^ found that the EB model had the best fit for body shape evolution for Lophiiformes (Fig. 4A). There was some support for EB dynamics for jaw shape as well, as this model had a ΔGIC (generalized information criterion) within 0–2 for all trees. The best-fit model for skull evolution was uncertain, and all three models were typically within 2 ΔGIC units across trees. Multivariate model fitting for suborders revealed clade-specific evolutionary dynamics. Shallow-water antennarioids were unique among suborders in that the OU model had the best fit for body shape evolution, and the attractor parameter was inferred to be high indicating strong stabilizing selection on shape. There was support for the EB model on antennarioid jaw shape evolution across all trees (likely driven by divergence of small-mouthed handfishes from large-mouthed frogfishes). For bathypelagic ceratioids, the best-fit model of body shape evolution was BM across all trees, though other models were within 2 units of GIC. For lophioids, the EB model was strongly supported as the best fit model of body shape evolution, driven by the divergence of the globose *Sladenia* from the strongly dorsoventrally flattened lophiids (Fig. 3A). There was strong support for an OU model for skull shape in ogcocephalids (Fig. 4B), as the skull of batfishes is very different from all other lophiiforms (Fig. 3).

**Figure 4:**
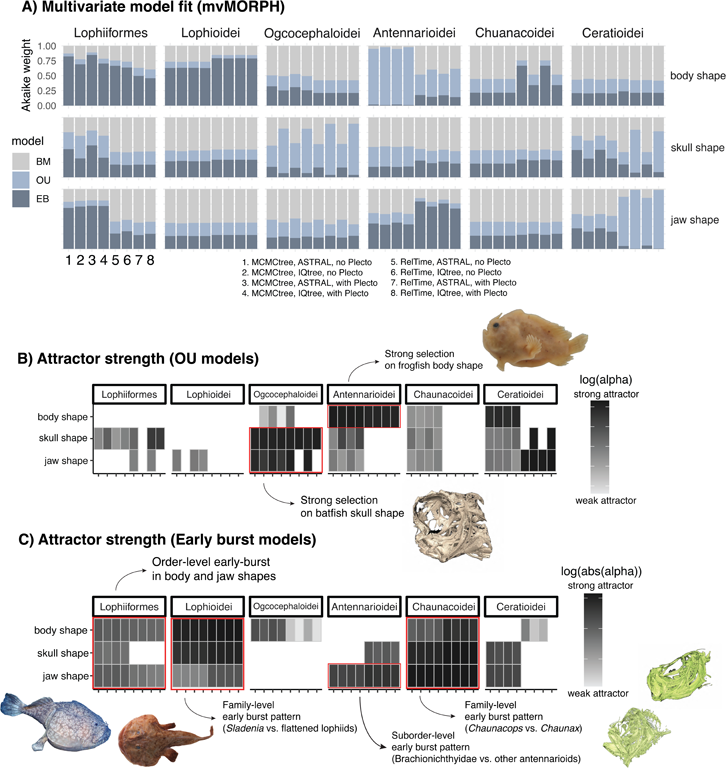
Results from multivariate model fitting using mvMORPH. (A) Akaike weight of three models of body, skull and jaw shape evolution across the eight trees. (B) Attractor strength (alpha) for OU models. (C) Attractor strength (alpha) for EB models. For panels B and C, poorly fitting models are not shown (i.e., only models within 2 ΔGIC units of the best-fitting model are shown).

We also performed univariate model fitting for the ten body shape linear measurements individually, revealing additional nuances (Fig. 5). As with multivariate analyses, the OU model had the best fit for all ten dimensions of antennarioid body shape indicating stabilizing selection. The OU model was also favored for most body shape dimensions in ceratioids, except standard length and interorbital length, for which BM was favored. As standard length becomes a reflection of body elongation when size-corrected with log shapes ratios^88^, this indicates that body elongation is less constrained than other shape dimensions in ceratioids. EB models did not have strong support in any of these analyses. This suggests that EB evolution detected with multivariate analyses (Fig. 4) was driven by the organization of trait combinations among clades.

**Figure 5:**
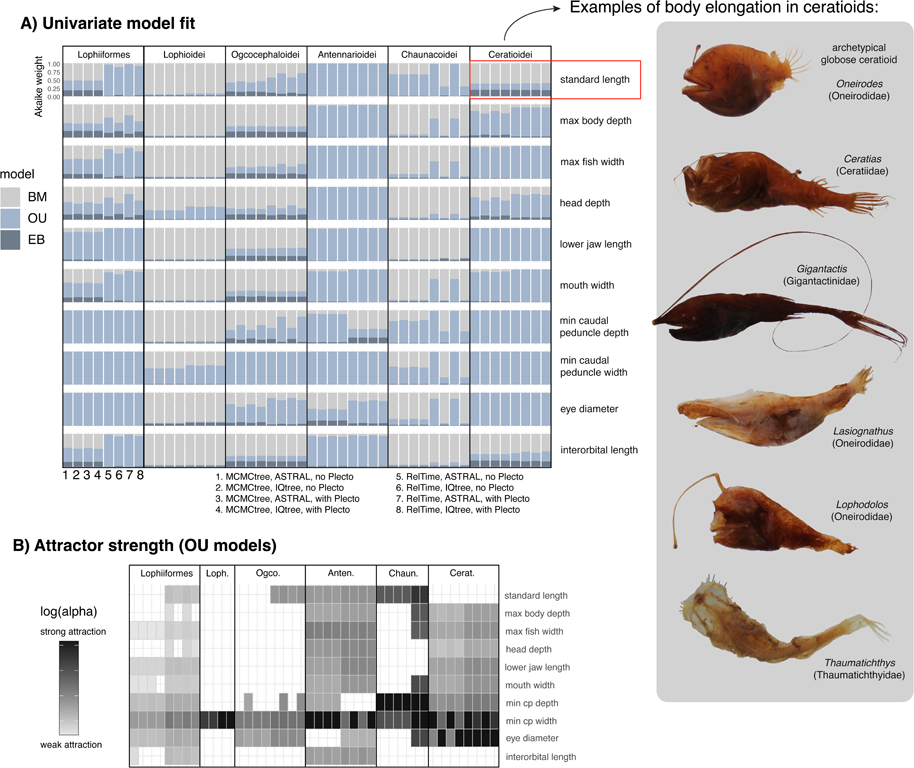
Univariate model fitting for individual body shape variables (Fig. S3). (A) Akaike weight support for three models across the eight time trees. (B) Attractor strength (alpha) for cases where the OU model had the best fit (greatest proportion of Akaike weight support).

Disparity-through-time analyses^89^ suggested that body shape disparity for Lophiiformes was relatively low within subclades early in the history of the clade but increased over time (Fig. S6), a signature of an early burst pattern of evolution for the order overall. Notably, ceratioids and antennarioids had high average subclade disparity in body, skull and jaw shapes throughout their entire history. This pattern indicates that subclades within these groups overlap greatly in morphology, a departure from the ordinal-level pattern.

PhyloEM models^90^ (Fig. S7) confirmed that adaptive peaks in body, skull and jaw shape reflected the same groups visible in morphospace (Fig. 3). Shifts in major body plans were generally associated with suborders, corroborating the early burst dynamics detected for body shape using other analyses (Fig. 4). An ancestral adaptive peak in overall skull shape was shared by the lophioids, ceratioids, and chaunacoids, with separate peaks for ogcocephalioids and anntenarioids. Lophioids and ceratioids each had unique adaptive peaks in jaw shape. Additional adaptive peaks were supported depending on which master tree was used, such as separate adaptive peaks in antennarioid and brachionichthyid jaw shapes when using MCMCTree (Fig. S7).

We inferred branch-specific evolutionary rates of body, skull, and jaw shape evolution across Lophiiformes using BayesTraits V4^91^ while fitting ten alternative models of trait evolution available within the software. Variable-rate models with a lambda transformation had the best fit in all cases. The slowest tip-associated evolutionary rates belonged to continental shelf taxa. Bathypelagic taxa had the highest rates of body shape evolution, and similar rates of skull and jaw evolution to deep benthic taxa (Fig. 6). Rate variation by branch revealed more complex patterns of trait evolution (Fig. 6). Evolutionary rates were generally low within the antennarioids across all three phenotypic datasets, with the exception of a few specialized species and along the stem branch leading to Brachionichthyidae. The ceratioids and ogcocephalids had several lineages with elevated rates corresponding to morphologically unique deep-sea genera. Therefore, we did not find that evolutionary rates slowed through time in deep-sea taxa, as predicted if ecological limits are driving the diversification process (Table 1). Rates of body shape evolution were high on the stem branches leading to Ceratioidei, Ogcocephalidae, and the dorsoventrally flattened lophiids, suggesting high rates are related to evolution of new body plans. Patterns were generally consistent between the two master trees (Fig. S8).

**Figure 6:**
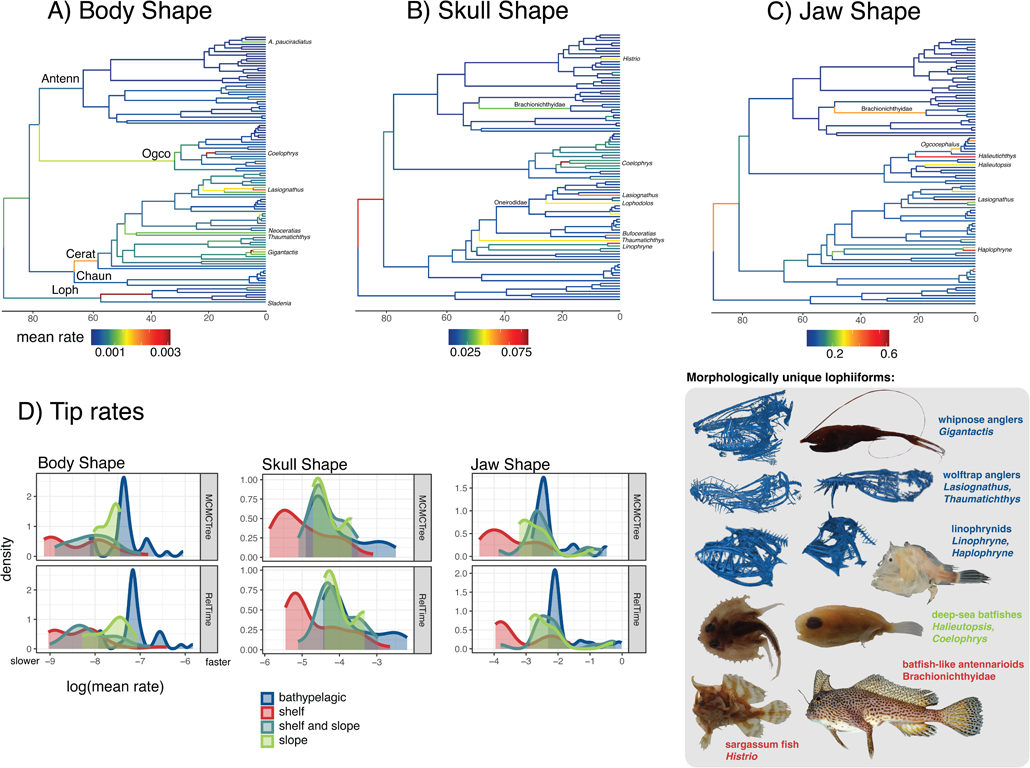
Rates of body, skull and jaw shape evolution inferred by BayesTraits. Panels A–C show branch-specific rates on the master MCMCTree. See Fig. S8 for a comparison between the master trees. Panel D shows tip-associated rates by habitat. See Fig. S8 for tip-associated rates by suborder. *Haplophryne* and *Brachionichthys* images from Fishes of Australia^121^.

## DISCUSSION

In this study we asked whether colonization of a novel but harsh environment should promote or constrain evolution. Colonization of new environments is generally believed to be a precursor to evolutionary radiation^58^. Yet, some environments should be inherently more constraining than others, potentially because there are few available niches or the challenges of that habitat only have a few viable solutions^14,50^. We examined the push-and-pull of these factors on evolution in the anglerfishes (Lophiiformes) with three guiding hypotheses (Table 1). We discuss the evidence for each of these hypotheses below.

### Early burst of lophiiform phenotypes

We found strong evidence that evolutionary dynamics for the order Lophiiformes as a whole evolved under early-burst dynamics. We found that an EB model had the best fit for body shape evolution (Fig. 4). The five suborders generally occupy distinct regions of the body shape morphospace, which was confirmed by phyloEM models (Fig. 3A, Fig. S7). Since four of five suborders are benthic, this supports the idea that benthic habitats in general contain more body shape diversity. This is potentially due to the greater topographic complexity of benthic versus pelagic habitats, which should promote niche evolution^22,55^. For example, the dorsoventrally compressed body plan only evolves in benthic fishes^22,92^, represented in Lophiiformes by the lophiids and ogcocephalids. These two clades diverged further in diet, with ogcocephalids eating small invertebrates^93^ and lophiids eating fishes^9^, explaining additional shape variation related to mouth size and position (Fig. 3). The early appearance of diverse body plans is also preserved in the fossil record: Monte Bolca fossils closely resemble living lophiids, antennarioids, and batfishes^94–99^.

Of all benthic environments, we should expect coastal shelf habitats, especially coral reefs, to promote phenotypic evolution^17,19,20,23,100^. Yet, the most reef-associated clade of lophiiforms, the antennarioids, was the most constrained in shape, fitting a pattern of “branch packing”^85^ (Fig. 3, Fig. S6). Unique among the five suborders, the OU model had the strongest support for multivariate body shape of antennarioids (Fig. 4) as well as for nearly all individual body shape variables (Fig. 5). Antennarioids also had the lowest rates of phenotypic evolution among Lophiiformes (Fig. 6, Fig. S8). The other lophiiform clade that specialized on continental shelf habitats, the genus *Ogcocephalus,* was also restricted in morphospace relative to ogcocephalids from deep-sea habitats (Fig. 3). Therefore, shelf habitats alone cannot explain the higher diversity of benthic lophiiforms, but rather the entire spectrum of benthic habitats including deep-sea environments must have played a role in generating this diversity.

### Evidence for adaptive radiation in the bathypelagic zone

Within ceratioids, most individual body shape variables evolved under an OU model (Fig. 5), and ceratioids were generally confined to a region of morphospace associated with small eyes and large jaws (Fig. 3), suggesting that these features are a response to bathypelagic conditions. For example, at these depths all light comes from bioluminescent point sources, which are bright enough for small eyes to detect^11^. Despite these constraints, we found that the bathypelagic ceratioids had the highest disparity when considering suborders individually, comprising 37– 41% of the total disparity of Lophiiformes (Tables S8, S9; Fig. S5). Ceratioids have been able to diversify as long as general constraints related to a bathypelagic existence are satisfied. This diversification includes instances of convergence on shallow-water shapes (Fig. 3C), as well as the evolution of entirely novel phenotypes related to predation (Fig. 6). Most strikingly, the “wolftrap” phenotype, in which the upper jaw and teeth are enlarged to ensnare prey, evolved twice independently (in *Lasiognathus* and *Thaumatichthys*) and is associated with high rates of evolution (Fig. 6). Ceratioids especially show a lot of diversity on the spectrum of body elongation, which was found to be evolving under BM (Fig. 5). Even though the “archetypical” ceratioid in popular imagination is globose, elongate forms have evolved repeatedly such as *Ceratias, Gigantactis, Lasiognathus*, and *Thaumatichthys* (Fig. 5).

Are ceratioids an adaptive radiation themselves (nested within the lophiiform radiation), or a different type of evolutionary radiation generated through non-adaptive processes^63,101,102^? This is not a pedantic exercise^63^, but is crucial for understanding fundamental questions about deep sea evolution. For example, given the paucity of resources, is adaptive radiation even possible in the deep sea? If so, does it conform to patterns described for terrestrial, freshwater and shallow marine adaptive radiations^60^? Ecological opportunity, the kindling that incites adaptive radiation, is thought to be highest upon colonizing a novel habitat that lacks competitors, especially when coupled with a key innovation that provides access to novel resources^56,58,60^. Ceratioids are by far the most diverse vertebrate clade in the bathypelagic zone today^6^. Their lure, large jaws, low metabolism, and extensible stomachs are shared with their benthic relatives^33,36^, which may have predisposed them for ecological success in the food-limited deep sea relative to non-lophiiform competitors^5,49^. They colonized this habitat from a deep-benthic ancestor and shortly after experienced a burst in lineage diversification rates (Fig. 2) and evolved novel phenotypes (Figs. 3, 6). Their sister group, the benthic chaunacids, have comparably low taxonomic and phenotypic diversity^102^ (Fig. S5). These pieces of evidence paint the picture of a potential adaptive radiation^103^.

While the EB model was developed to characterize adaptive radiation based on Simpson’s conceptualization^62,104^, in practice this model seems to be a poor representation of many adaptive radiations^61^ including the ceratioids. Despite rapid lineage diversification early on (Fig. 2B), there is little evidence for a similar early burst of phenotypic evolution (Figs. 4, 5). Phylomorphospace analyses (Fig. 3) and diversity-through-time plots (Fig. S6) showed phenotypic overlap in body, skull and jaw shapes throughout the entire history of ceratioids, distinct from the early burst pattern seen for Lophiiformes as a whole (Fig. S6). BayesTraits analyses showed that relatively young lineages have experienced rapid rates of evolution (Fig. 6). The wolftrap and whipnose anglers are examples of lineages that have evolved novel prey capture strategies relatively recently in the context of the ceratioid radiation. Although ceratioids are at least 30 million years old (Fig. 1), it seems unlikely that they are exhausting ecological opportunity such that they can no longer diversify^105,106^. We know very little about what ecological opportunity looks like in the deep sea. On one hand, the bathypelagic zone is the most food-limited and environmentally homogeneous habitat on Earth. On the other hand, population density of ceratioids is very low, and populations are spread across the globe^6,45^. Environments with patchy resources should promote coexistence by preventing any species from becoming dominant^107^. Therefore, resources are very limited, but competition should also be very low^108^.

A remaining mystery is the degree to which non-adaptive processes contributed to the diversity of ceratioids. Relaxed selection due to ecological release is believed to play an important role in the initial stages of adaptive radiation by broadening phenotype diversity, giving way to a later stage of disruptive selection among these phenotypes^56,57,60^. Yet, some authors hypothesize that selection on body shape is perennially relaxed in the bathypelagic zone^29^. Bathypelagic fishes have neither the demands of shallow-water pelagic predators for pursuing prey^52^, nor the challenges of navigating obstacles like benthic fishes^22,26^. Therefore, shape disparity may have accumulated over time in this habitat if new shapes are neutral with respect to selection. Ceratioid body elongation may fit this pattern of evolution (Fig. 5). While elongation is also a common theme for benthic-to-pelagic transitions in shallow-water fish clades^22,53,55^, the difference is that elongation in these groups is under selection for reducing drag for sustained swimming. Videos in-life suggest that globular^109^ and elongate^110^ ceratioids are both incapable of sustained swimming due to their reduced skeletal and muscular architecture. It is unclear why elongation would be under selection for some ceratioids but not others. Similarly, ceratioids have diverse jaw and tooth shapes which yield differences in function^111^, yet they seem to be opportunistic generalist carnivores based on largely anecdotal evidence^9,112^. We know from videos and trawl records that ceratioids show some differences in hunting behavior^111^, and a few genera inhabit the benthic boundary layer with demersal prey making up some portion of their diet^6,39,110^. Otherwise, evidence of phenotype-ecology matching is lacking for ceratioids, whereas this has been a crucial piece of evidence for the adaptive radiation process in terrestrial and shallow-water organisms that are easier to study^59,103,113^. Without this evidence, it is difficult to understand why so many body and jaw shapes have evolved in ceratioids and the strength of disruptive selection on these different shapes.

### Phenotypic constraint in shallow-water lophiiforms

Phenotypic stasis could arise from the lack of ecological opportunity (external constraints) or functional limitations (internal constraints)^24,114,115^. Slow and constrained evolution of shallow-water frogfishes is unexpected because it contradicts the trend seen in other fish clades. Wrasses (Labridae) show higher diversification on reefs which is partially driven by exploration of novel phenotypes to acquire new resources^19,116^. Grunts (Haemulidae) are not as trophically diverse as wrasses yet still have faster phenotypic diversification on reefs, probably due to finer partitioning of existing niches^100^. Unlike wrasses and grunts, frogfishes did not evolve novel diets nor partition dietary resources more finely than other lophiiforms. No lophiiform has evolved herbivory or planktivory, so frogfishes are not taking advantage of the full array of opportunities provided by coastal habitats^19–21,23^. They are indiscriminate carnivores with extensible stomachs^33^ capable of the largest volume of oral expansion known among reef fishes, allowing them to catch prey from long distances using suction feeding. Their prey capture success rate is therefore much higher than other reef fishes^33^. Evolutionary innovations may result in specialization instead of diversification if the innovation does not broaden the array of potential resources^117,118^. We might therefore conclude that the frogfish bauplan functions in a variety of coastal environments by increasing their success as a generalist carnivore, and there is little external incentive to modify it even with the genetic or developmental ability to do so. Note that while frogfishes are constrained in shape, they are highly variable in color allowing them to mimic sponges, corals and urchins^33^; they likely have very high rates of color evolution.

### The timeline of lophiiform evolution

A novel result from our study is that the crown age of Lophiiformes is well within the Cretaceous (Fig. 1). Even our trees with the youngest estimates have confidence intervals extending to ∼76 Ma (Appendix A4, Table A4). Yet, other studies found that Lophiiformes have a Cenozoic origin as part of a post-K/Pg diversification event affecting spiny-rayed fishes broadly^72,119^. The primary reason for the older age estimates in our study is our use of six fossil calibrations from Monte Bolca which included crown representatives of Lophoidei and Antennarioidei (Fig. 1). Older age estimates were not limited to analyses using †Plectocretacoidea, a controversial Cretaceous fossil^77^. We believe that at minimum, the age of lophiiform subclades were underestimated by prior studies (discussed in detail in Appendix A4). Past studies used at most three Monte Bolca calibrations for Lophiiformes (Appendix A4, Table A5). This was most likely due to lower taxonomic sampling compared to our study, providing fewer nodes to place calibrations.

The fossil record gives no direct evidence of lophiiforms prior to the Eocene. Yet, the presence of several lineages in Monte Bolca, including crown representatives of two suborders, strongly suggests that Lophiiformes were already diverse by then. A Cenozoic crown age of Lophiiformes would require that suborders diversified rapidly in the intervening 17.5 million years between the K/Pg boundary and Monte Bolca^94^. Yet, no such rapid radiation is visible in our phylograms (Appendix A1). Therefore, we suggest that a Cretaceous origin of Lophiiformes is the best explanation to reconcile molecular data with the fossil record.

Notably, Hughes et al.^74^ recently found a crown age of Labridae of ∼79 Ma using an expanded fossil calibration list compared to past studies, which found a Cenozoic crown age. Both Lophiiformes and Labridae are members of Eupercaria, one of nine series within Percomorpha^120^ and one of the groups implicated in the post-K/Pg radiation of acanthomorphs. It remains to be seen whether an older age of labrids and lophiiforms changes the finding of rapid post-K/Pg radiation of acanthomorphs found by recent studies^72,119^. Regardless, it is clear that improved taxonomic sampling made possible by collections^68^ combined with paleontological systematics^77,95,97^ stands to transform our understanding of the timescale of fish evolution.

### Conclusions

We combined a well-sampled phylogenomic hypothesis with three-dimensional morphometric data to examine the tempo and mode of evolution following habitat transitions in anglerfishes. The bathypelagic anglerfishes experienced a burst of lineage diversification and now contain the greatest phenotypic diversity of all lophiiform clades, whereas continental shelf lineages are relatively constrained in morphology. These findings contradict ecological expectations, since we expect complex coastal habitats to promote niche evolution relative to the homogeneous bathypelagic zone. Our findings prompt new questions about deep-sea ecology and evolution, such as to what extent radiation is possible in harsh environments, as well as the role of adaptive versus neutral processes for generating diversity in these settings.

## ACKNOWLEDGEMENTS

We are grateful to the following museums and personnel for providing voucher specimens and/or tissues: UWFC (Katherine Maslenikov), FLMNH (Robert Robins), SIO (Ben Frable and Phil Hastings), AM (Amanda Hay), CSIRO (Alastair Graham and John Pogonoski), MCZ (Andrew Williston and Meaghan Sorce), FSBC (Eric Post), YPM (Gregory Watkins-Colwell and Tom Near), KU (Andrew Bentley and Leo Smith), LSUMZ (Prosanta Chakrabarty and Seth Parker), LACM (Todd Clardy and Bill Ludt), and USNM (Chris Huddleston and Diane Pitassy). Tracey Sutton kindly gifted tissues to UWFC.

We thank Adam Summers, Karly Cohen, and Zach Heiple for help with CT scanning at Friday Harbor Laboratories. Scanning was supported by the oVert TCN (NSF DBI-1701665). Sarah Panciroli, Divinity Patterson, Leo MacLeod, Jonathan Huie, and Jenny Gardner assisted with collecting and processing CT scans and were funded by the William W. and Dorothy T. Gilbert Ichthyology Research Fund. Sarah Friedman and Julien Clavel gave advice on morphometric approaches.

We thank the MERLAB at University of Washington for assistance with wet lab work. Exon capture and sequencing was performed by Arbor Biosciences and funded by FishLife (NSF DEB-1541554). Computational resources were provided by the University of Oklahoma Supercomputing Center for Education and Research (OSCER). Liam Ward assisted with data curation and quality control.

ECM was supported by an NSF Postdoctoral Fellowship (DBI-1906574) and NSF DEB-2015404. PBH was supported by an NSF Postdoctoral Fellowship (DBI-2109469). KE was supported by NSF DEB 2237278. DA was supported by NSF DEB-2144325 and NSF DEB-2015404.

## METHODS

### Data acquisition

We generated new genomic data from tissue samples associated with museum specimens (Table S1). New data was collected from 152 individuals from 120 species of Lophiiformes. DNA was extracted using the DNeasy Blood and Tissue Kit (Qiagen, Valencia, CA). We shipped DNA extractions to Arbor Biosciences (Ann Arbor, MI) for library preparation, target enrichment, and sequencing. Sequencing of pair end 150 bp reads was completed on a HiSeq 4000 with a total of 192 samples multiplexed per lane. Target capture probes were based on a set of 1,105 single-copy nuclear exon markers designed for fish phylogenomics (Eupercaria bait set of Hughes et al. ^73^). An additional 19 nuclear legacy markers, as well as mitochondrial DNA, were also targeted using this probe set. Information for individuals with new genomic data can be found in Table S1. We mined exons from genomes available on NCBI for eight additional outgroup and two ingroup species. Our outgroup sampling (Table S1) included one holocentrid (representing the sister lineage to Percomorpha), one ophidiid (the earliest diverging member of Percomorpha), one pelagiarian, two syngnatharians, 18 tetraodontiforms and 15 additional eupercarians^75^.

Taxonomic sampling was improved using two approaches. First, we mined exons from published UCE alignments^71,72^. We assembled the raw reads from these studies into loci using the FishLife Exon Capture pipeline described below. Between 5–357 exons (mean 40.3 per individual) were successfully mined for 93 individuals representing 48 species. After quality control steps, 12 species were retained in the “final” alignment (see below) on the basis of these mined exons. Information for individuals with exons mined from UCEs can be found in Table S2. Second, we downloaded legacy markers for 10 species available from GenBank (Table S3).

These species had between 1–5 markers available. Due to the large amounts of missing data introduced in the alignment, we only pursued legacy markers for species that would be new to our dataset. After quality control steps, two genera and six species not available elsewhere were retained in the “final” alignment on the basis of these legacy markers (Table S3).

Our final taxonomic sampling when combining all data and remaining after all quality control steps (see below) was 132 ingroup species (37.8% of species and 78.1% of genera in Lophiiformes) and 20 of 21 families (all but the monotypic Lophichthyidae). Suborder-level sampling is as follows: 9 species of Lophioidei (32.1% of species and all four genera), 21 species of Ogcocephaloidei (28.7% of species and eight of ten genera), 40 species of Antennarioidei (62.5% of species and 77.3% of genera [17 of 22 genera]), eight species of Chaunacoidei (50% of species and both genera), and 54 species of Ceratioidei (32.1% of species and 74.3% of genera [26 of 35 ceratioid genera]).

### Assembly, alignment and quality control

Assembly, initial raw data quality control steps, and alignment were conducted using the pipeline^73^ available at https://github.com/lilychughes/FishLifeExonCapture. Low quality raw reads and adapter contamination were trimmed using Trimmomatic v.0.39^122^. Trimmed reads were mapped against the reference sequences used for probe design with BWA v.0.7.17^123^ and PCR duplicates were removed using SAMtools v.1.9^124^. An initial sequence for each marker was assembled with Velvet v.1.2.10^125^, and the longest contig was used as a reference sequence to extend contigs using aTRAM 2.2^126^ with the Trinity v.2.2 as the assembler^127^. Redundant contigs were excluded with CD-HIT-EST v.4.8.1^128,129^, and open reading frames for the remaining contigs were identified using Exonerate v.2.4.0^130^. Redundant contigs with reading frames exceeding 1% sequence divergence were discarded.

New data, mined exons from UCEs, and legacy markers were aligned using MACSE v.2.03^131^ with the -cleanNonHomologousSequences option. After alignment, we discarded 26 exons with low capture efficiency (those with <50 taxa). Next, some legacy markers can retain paralogues when obtained using our target capture probe set and deserve additional scrutiny^73^. For these markers, we checked their gene trees by eye for pseudogenes. Five exons had pseudogenes (rhodopsin, zic1, sh3px3, plag2, and ENC1) and were excluded from our dataset. After these steps, the dataset contained 1,092 markers. This number included 1,077 FishLife exons, 13 additional nuclear legacy markers, and two mitochondrial legacy markers (CO1 and ND1).

Further quality control steps follow those described by Arcila et al. ^132^. We performed branch length correlation (BLC) tests^133^ to detect within-gene contamination that may not be easily detectable once genes are concatenated. The logic of this test is that contaminated sequences will show very long branches once constrained to a reference topology. We generated a reference phylogeny using the program IQ-TREE MPI multicore v.2.0^134^ based on the concatenated alignment of all 1,092 genes and using mixture models^135^. We then generated gene trees for each marker with the topology constrained to match the reference phylogeny. We generated a branch-length ratio for every taxon in every gene tree, which was the length of the branch in the gene tree over the length of the corresponding branch in the reference tree (after pruning the reference tree to the same individuals contained by the gene tree). All branches with a ratio >5 were flagged, and all flagged branches were then checked by eye. Ultimately, 1,416 sequences (taxa in gene trees) were discarded from our dataset due to suspected contamination (very long branches in the gene trees). In addition, two taxa were later dropped entirely from the dataset because we observed them to have extremely long branches across many gene trees (Table S1).

Species identifications of sequences were confirmed with two complimentary approaches. First, for species with more than one individual sampled, we checked the phylogram produced containing all individuals (see below) by eye with the assumption that species should be monophyletic. Second, we referenced CO1 sequences against the BOLD (Barcode of Life Data System) database^136^ using scripts from the “fishlifeqc” package available at: https://github.com/Ulises-Rosas/fishlifeqc. For genera with short branch lengths (specifically *Ogcocephalus, Chaunax, Oneirodes, Gigantactis,* and *Himantolophus*), we could not obtain confident species identifications using BOLD, and species were often non-monophyletic. This is potentially due to incomplete lineage sorting after rapid speciation, low substitution rates, and/or misidentification. We checked the literature for evidence of “taxonomic inflation” in these genera (in which more species are described from morphology than exist based on molecular divergence), and believed this scenario to potentially apply to *Ogcocephalus* and *Himantolophus* (discussed in Appendix A2). For individuals outside of these five genera that failed our checks, we checked the voucher specimen whenever possible. This resulted in the re-identification of two museum specimens. We also flagged four previously published sequences from UCE studies as misidentified. If we could not confirm an individual’s identification because there was no CO1 sequence and no conspecific replicate, we referred to the literature to check if the position of the species in the phylogeny was as expected compared to prior hypotheses, or at least within the expected genus or family. We preferred to retain individuals for the “final” alignment (see below) of which we could be reasonably confident of their species identification. Quality control results for all individuals can be found in Table S1 (new genomic data) and Table S2 (individuals taken from UCE alignments).

### Phylogenomic inference

We produced trees from two sets of alignments made from the 1,092-marker-dataset. The first “all individuals” set contained all sequences that made it past the BLC step of quality control (*n*=258 ingroup individuals). The tree made from this alignment (Appendix A1, Figure A1) was checked by eye to confirm species identity of sequences (for those species with multiple individuals in the dataset) as the final step of quality control (see above). The second “final” alignment was produced by choosing one individual to represent each species (*n*=132 ingroup species). When multiple conspecific individuals were available, this representative was always the individual with the greatest number of genes assuming no quality control flags (Tables S1– S3). This “final” alignment was the one used to produce the phylograms used for time calibration and comparative methods. After pruning down to nearly half the number of individuals between the “all-individuals” and the final alignment, genes were un-aligned using the “unalign.md” script within the Goalign toolkit^137^, then re-aligned. The final alignment was 457,635 base pairs long, and alignments for individual markers varied in length from 105–2,682 bp (mean 420 bp).

All 1,092 markers were concatenated using utility scripts in the AMAS package^138^. Trees were constructed with maximum likelihood using the program IQ-TREE MPI multicore v.2.0^134^ implementing mixture models^135^ (option -m set to “MIX{JC,K2-,HKY,GTR}). Support was measured using 1000 Ultrafast bootstrap replicates ^139^ with the “-bnni” option to reduce the risk of overestimating support due to severe model violations.

To account for potential incomplete lineage sorting, we also performed a multi-species coalescent analysis using ASTRAL-II v.5.7.1^140^ based on gene trees estimated using IQ-TREE with the same settings as above. Prior to use with ASTRAL, nodes within gene trees with bootstrap values <33% were collapsed into polytomies to reduce noise^141^. Support was evaluated using local posterior probabilities^142^ (option “-t 3”).

### Divergence time estimation

We assembled a list of 21 node calibrations from the literature, including 8 outgroup and 10 ingroup fossil calibrations based on well-preserved articulated skeletal remains, as well as geologic calibrations based on the Isthmus of Panama to constrain the divergence time of three sister-species pairs. Calibration details and justifications are given in Appendix A3. Following the recommendations by Parham et al. ^143^, we established minimum age constraints (i.e., the youngest fossil ages) to determine lower bounds for each calibration.

We used two calibration schemes including or excluding the controversial fossil †*Plectocretacicus clarae*, which we placed on the MRCA of Tetraodontiformes and Lophiiformes ^75^. The extinct superfamily †Plectocretacoidea is purportedly a stem tetraodontiform, and phylogenetic analyses using morphological characters place it as the sister to all remaining Tetraodontiformes^77,144,145^. The earliest plectocretacicoid fossils are 94 million years old^144^. Therefore, due to the apical position of Tetraodontiformes within acanthomorphs, and the sister group relationship between Tetraodontiformes and Lophiiformes, this fossil has potential to greatly increase the age of early nodes in the phylogeny of Lophiiformes. However, some authors do not believe †Plectocretacoidea are related to Tetraodontiformes, or at least that the evidence for such a relationship is uncompelling^119,146–148^.

We produced eight alternative time trees using either the IQ-TREE (concatenated) or ASTRAL (coalescent) trees, the fossil calibration scheme with or without †*Plectocretacicus*, and using either MCMCtree or RelTime as the calibration method. Both MCMCTree and RelTime are feasible for use with genomic-scale datasets, but these approaches are otherwise quite different. MCMCTree uses a birth-death tree prior and an independent rates clock model in which rates follow a log-normal distribution in a Bayesian framework^78,79^. RelTime does not use priors on lineage rates, and instead computes relative time and lineage rates directly from branch lengths in the phylogram (the “relative rate framework”)^80,81^. Note that RelTime tends to underestimate divergence times for branches with very few molecular substitutions, unlike methods that include a tree prior^149,150^.

For MCMCTree, fossil calibrations used uniform distributions and geologic calibrations used Cauchy distributions (Appendix A3, Table A3). We used distribution densities based on the algorithm proposed by Hedman^151^. This approach uses a list of fossil outgroup age records based on the oldest minima to produce a probable distribution of the origin of a given clade (details in Appendix A3). From the distribution estimated for each calibration, we extracted the 95% confidence interval to set the soft upper bound (maximum age) for MCMCTree, and to calculate the mean and standard deviation for log-normal distributions in RelTime.

We implemented MCMCTree analyses using the PAML v.4.9h package^152^. We divided the alignment into two partitions: 1^st^ and 2^nd^ codon position, and 3^rd^ codon position. We used the HKY85 substitution model and the independent rate relaxed clock model. Additional prior parameters were set as follows: BDparas: 1, 1, 0.38; kappa_gamma: 6, 2; alpha_gamma = 1, 1; rgene_gamma = 2, 200, 1; sigma2_gamma = 2, 5, 1. To improve computation time, we first used the approximate method to calculate the likelihood^79^. MCMC chains were run twice independently for 20 or 30 million generations as needed to converge (number of samples= 200000, sample frequency= 100 or 150, and burnin= 2000). We used Tracer v1.7.1^153^ to check for convergence.

RelTime uses a maximum likelihood framework implemented in the software MEGAX^154^. For the IQ-TREEs, we applied the RelTime-Branch Lengths approach, employing a Max Relative Rate Ratio of 20, with the tree topology serving as the input. For the ASTRAL trees, we used RelTime-ML with the GTR+I model while maintaining the default settings to optimize branch lengths. The ASTRAL topology along with the concatenated alignment were used as inputs. This is necessary because the ASTRAL tree was made from gene trees and not estimated directly from the alignment.

Some analyses were repeated for all eight time-calibrated trees in order to incorporate variation in topology and divergence times. Analyses involving complex visualizations were repeated on two designated “master” trees: the IQ-TREE calibrated with the scheme including †Plectocretacoidea using either MCMCTree or RelTime (hereafter “master MCMCTree” or “master RelTime tree”). This was because of the three methodological choices for time calibration, the decision with the largest impact was MCMCTree versus RelTime (Fig. 1; Appendix A4).

### Ancestral habitat and lineage diversification rates

Following Miller et al. ^49^, we used BioGeoBEARS v.1.1.3^155^ to infer ancestral habitats. This approach allowed us to code species as occurring in more than one “region”. Our analysis included three regions: benthic continental shelf, benthic continental slope to abyssal plain, and the bathypelagic zone. Habitats were coded based on: FishBase^156^, Fishes of Australia^121^, Pietsch^6^, and Friedman et al. ^22^ (Table S4). The maximum number of regions allowed per species was set to two. We compared the fit of six alternative models using Akaike weights^157^. These were: DEC^158^, DIVA-LIKE^159^, BAYAREA-LIKE^160^, and their equivalents with the +J parameter (Table S5). We performed these analyses on the two master trees, with results being nearly identical; therefore, only results using the master MCMCtree are shown (Fig. 2).

We estimated lineage diversification rates using the MiSSE framework (missing state speciation and extinction)^83^ implemented in the hisse R package v2.1.1. MiSSE operates like HiSSE^161^ but does not consider the influence of any characters chosen by the researcher, instead modelling rate shifts agnostic of any *a priori* hypothesis. We performed analyses for all eight time trees individually. We were concerned that taxonomic inflation could inflate speciation rates in the genera *Himantolophus* and *Ogcocephalus* (Appendix A2). Therefore, we also performed analyses on a set of eight trees with these genera pruned to two species (to retain the crown age), for a total of sixteen sets of analyses (Table S6). We compared the fit of models with 1–10 rate classes, setting a global sampling fraction of 38%. Following recommended practices^84^, we model-averaged rates among the set of models with >5% of the relative Akaike weight, where the contribution of each model towards the mean was proportional to its Akaike weight. We plotted model-averaged rates onto the branches of the tree using the gghisse package v.0.1.1^162^. Note that SSE models avoid issues of identifiability raised by Louca and Pennell^163^ because they incorporate multiple information sources to infer rates^164^.

### Phenotypic datasets

Body shape was measured using linear measurements from museum specimens. We took eight measurements following Price et al.^88^ (standard length, maximum body depth, maximum fish width, head depth, lower jaw length, mouth width, minimum caudal peduncle depth, and minimum caudal peduncle width) plus two additional measurements (eye diameter and interorbital distance). Measurements are shown in Fig. S3. We took measurements using digital calipers with a minimum resolution of 0.1 mm. Measurements were size corrected using log-shapes ratios^88,165^: each variable was divided by the geometric mean of standard length, maximum body depth, and maximum fish width (a more realistic way to approximate size for globular fishes versus using a single measurement like standard length), and then log-transformed. For quality control, we flagged measurements that were outside the inter-quartile range for the genus, and specimens with flags were excluded. The final dataset after quality control contained measurements for 327 individuals from 112 species (representing 84.8% of tips in the phylogeny), in which 1–9 individuals per species were measured (mean 2.9 individuals per species). No male ceratioids were used. The dataset with voucher information is available in the Dryad package associated with this study. The species means for each trait were used for phylogenetic comparative methods.

Skull shape was measured using three-dimensional geometric morphometrics collected from micro-computed tomography (micro-CT) scans of museum specimens^166^. Scans were collected at the Karel F. Liem Bio-Imaging Center at the University of Washington Friday Harbor Laboratories and Rice University. Skulls were segmented from scales and the rest of the body using Amira v.2020.3^167^ and exported as mesh files. Mesh files were digitized with 111 three-dimensional landmarks (41 point and 70 semi-sliding; Fig. S4) in the software Stratovan Checkpoint^168^. Landmarks were treated as bilaterally symmetrical and thus only placed on the left side of the skull^169^. Our CT scan dataset contained 100 species of Lophiiformes (*n*=1 scan per species) representing 75.7% of the tips in our phylogeny (Table S7). Of these, 38 are new to this study, 33 were previously published^111^, and 29 were downloaded from the online repositories MorphoSource (https://www.morphosource.org/) or Virtual Natural History Museum (http://vnhm.de/VNHM/index.php).

The highly mobile and interconnected nature of the teleost fish skull can increase the likelihood of preservation artifacts^21,170,171^. To reduce these artifacts, we performed a local superimposition to standardize the position of individual skull elements^172^ before any downstream analyses using shape data.

### Phenotypic evolution

We performed all analyses of phenotypic evolution on three datasets: body shape, whole skulls, and the oral jaws, with the latter two based on CT scans. To measure jaw shape, we isolated the 41 (13 point and 28 semi-sliding; Fig. S4) landmarks placed on the premaxilla, angular, and dentary. The same set of bones were isolated by Heiple et al.^111^ in their analysis of jaw and tooth shape using linear measurements.

We visualized shape variation using a phylomorphospace analysis^85^ performed with the function “gm.prcomp” from the geomorph R package v.4.0.5^173^. For use with downstream analyses, we exported the PC scores for the number of axes summing to 95% (body shape) or 85% (skull and jaws) of the variance. For example, when using our two master trees this number was six axes for body shape, 28 axes for skulls, and 12 axes for jaws. We did this for all eight time trees, as well as the phylogeny for each suborder isolated from the eight trees, for a total of 48 sets of phylogenetically-corrected PC scores.

We calculated disparity by suborder and habitat category using a test of morphological partial disparities for the overall mean^86^ (Tables S8, S9). We plotted disparity-through-time using the “dtt” function in the geiger package v.2.0.11^174^. The observed disparity was compared to a Brownian motion null model that was simulated 1,000 times across the master MCMCTree ^89^.

We performed univariate model fitting analyses for the ten body shape variables individually using the “FitContinuous” function in geiger, inputting all 48 trees, for a set of 480 analyses. We compared the fit of three models using Akaike weights: Brownian motion (BM), single-peak Ornstein-Uhlenbeck (OU), and Early Burst (EB)^61^.

We performed multivariate model fitting using PC scores from the three phenotypic datasets, inputting all 48 trees, summing to 144 sets of analyses. Following Clavel et al. ^87^, we fit models using penalized likelihood with the “fit_t_pl” function in RPANDA v2.2^175^ using the rotation-invariant ridge quadratic null penalty (method=”RidgeAlt”) and accounting for measurement errors (option SE=TRUE). The fit of the same three models (BM, OU, EB) was assessed using the generalized information criterion with the “GIC” function in mvMORPH v.1.1.7^176^, as GIC is appropriate for penalized likelihood. The relative model support was then compared using Akaike weights. In addition, we fit multiple-peak OU models to detect Simpsonian adaptive regimes using the PhylogeneticEM package v.1.6.0^90^, performing these analyses on the two master trees. We compared the fit of models with 0–20 regime shifts using the selection criterion adapted by Bastide et al.^90^ (Fig. S7).

To infer branch-specific evolutionary rates we performed reversible-jump MCMC analyses within BayesTraits V4^91^. We investigated rates of evolution in body, skull and jaw shape, for our two master trees, for a set of six analyses. Following Coombs et al.^177^, we used Bayes Factors to evaluate the relative support of ten models: Brownian motion, kappa, delta, lambda, and OU tree transformations, each with single- and variable-rate alternatives. We accounted for correlated trait evolution with the setting “TestCorrel” which constrains the correlation between trait axes to zero. Chains were run for 200 million generations with a burnin of 30%. A stepping stone sampler was used to estimate the marginal likelihood with 100 stones to run for 1,400,000 generations after convergence. Analyses were run twice, and convergence of the runs was confirmed based on trace plots and Gelman diagnostics near 1, using the packages coda v.0.1.9-4^178^. BayesTraits output was processed using utility functions from the packages BTProcessR v.0.0.1^179^, BTRTools 0.0.0.9^180^ and scripts written by R. Felice^181^. The output of variable-rate analyses is a set of phylogenies where each branch was scaled by its Brownian motion rate of evolution. We plotted the mean rate for each branch based on the best-fit model, and extracted tip-associated rates to compare rates by habitat.

