## Appendix A1 for "Phylogenomics reveals the deep ocean as an accelerator for evolutionary diversification in anglerfishes"

### APPENDIX A1: Phylograms

**Figure A1:** Maximum likelihood phylogram including all individuals used for quality control (validating species identifications).

**Figure A2:** Maximum likelihood phylogram made using IQ-TREE using set of representative individuals for each species.

**Figure A3:** Coalescent phylogram made using ASTRAL using set of representative individuals for each species.

**Figure A4:** Comparing the topology from IQ-TREE and ASTRAL.

**Figure A5:** RelTime master tree with 95% confidence intervals.

**Figure A1:** Phylogram including all sequences for purposes of validating species identifications.
Tree was constructed using IQ-TREE; ultrafast bootstrap values are shown.
For sequence details and barcoding results, see Table S1 (new data), Table S2 (exons mined from UCEs) and Table S3 (legacy markers).
Names in boldface indicate ingroup sequences that were chosen to represent the species in the final trees.

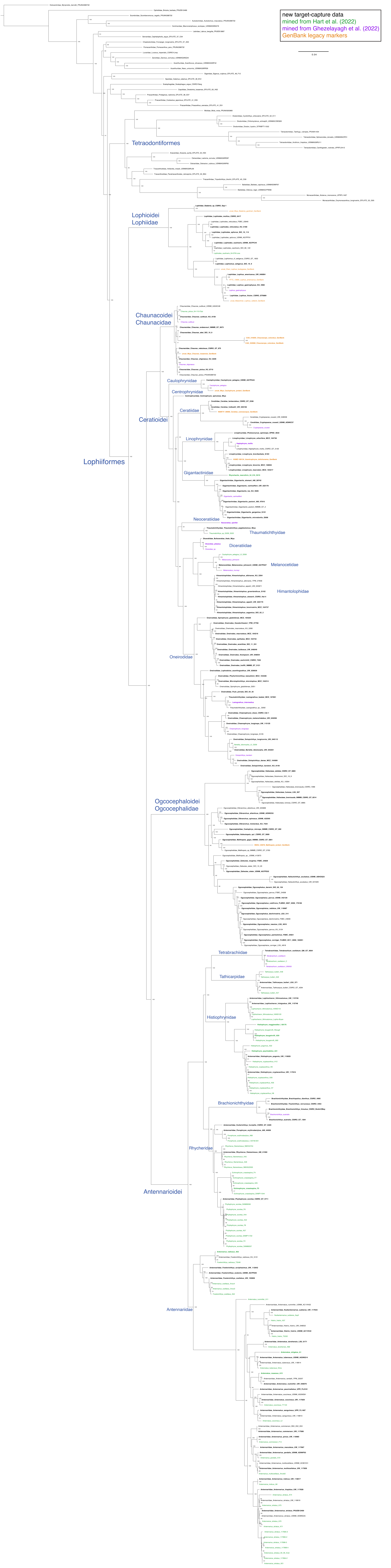

**Figure A2:** Phylogram made using IQ-TREE based on 1,092 concatenated markers. Sampling was pruned to one sequence per species for use with phylogenetic comparative methods. For tree showing all sequences, see Fig. A1. Ultrafast bootstrap support values are shown.

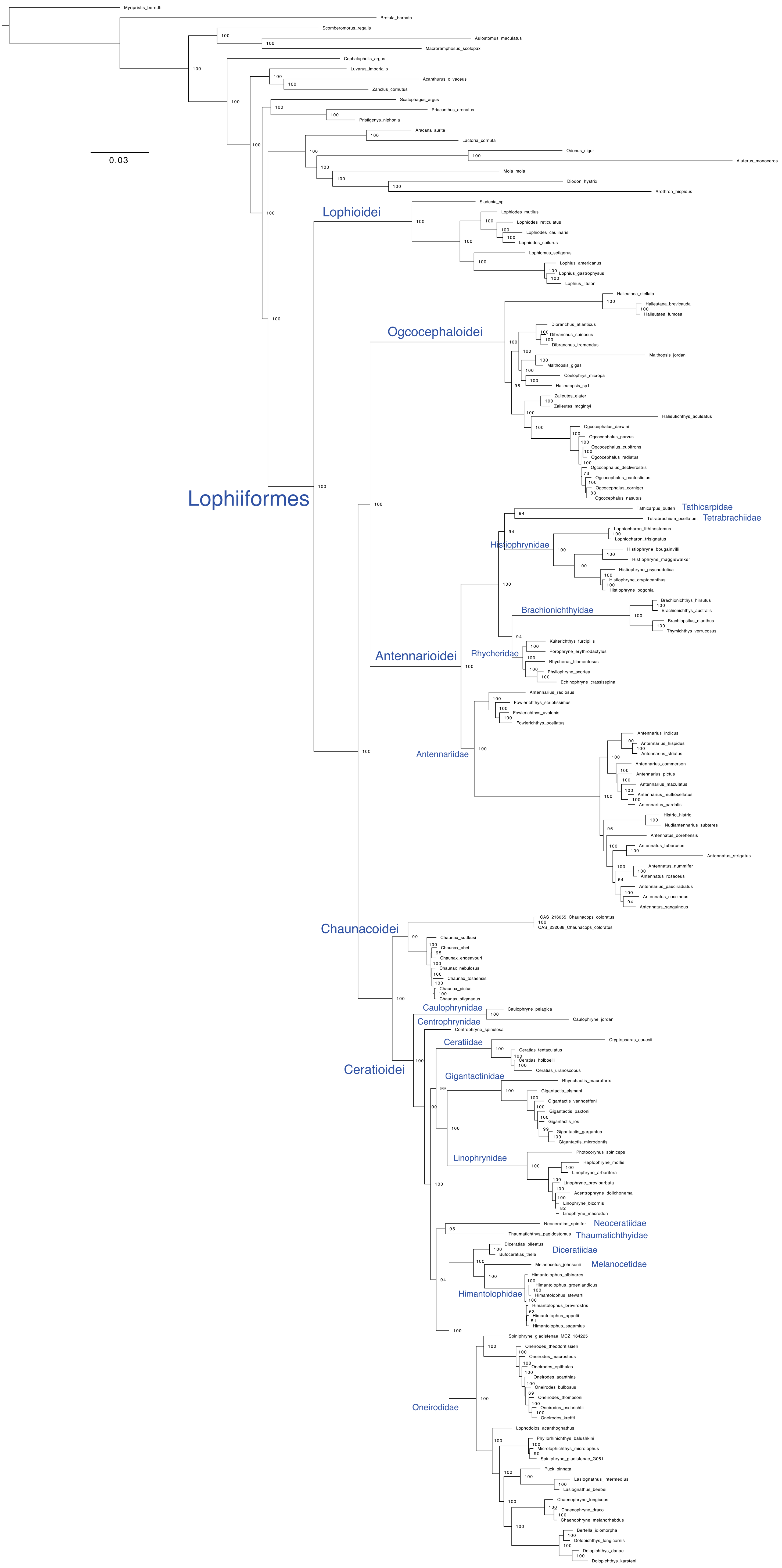

**Figure A3:** Phylogram made using ASTRAL-II based on 1,092 gene trees.
Sampling was pruned to one sequence per species for use with phylogenetic comparative methods.
For tree showing all sequences, see Fig. A1. Quartet support values are shown.

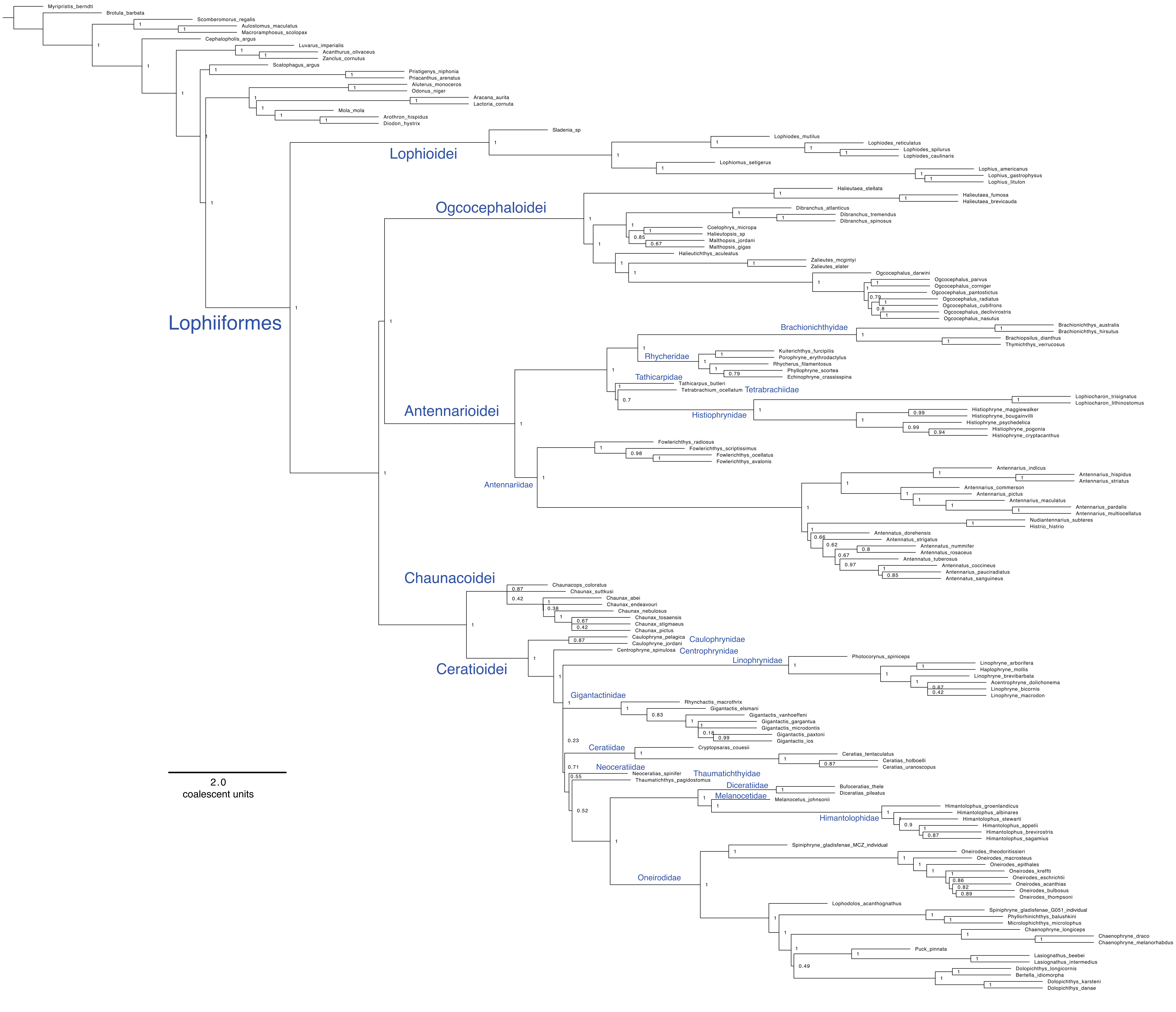

**Figure A4:** Comparing the topologies of concatenated (IQ-TREE) and coalescent (ASTRAL) phylogenies.
The IQ-TREE cladogram is shown.
Black circle = taxa form a clade in the ASTRAL tree
White circle = taxa do not form a clade in the ASTRAL tree

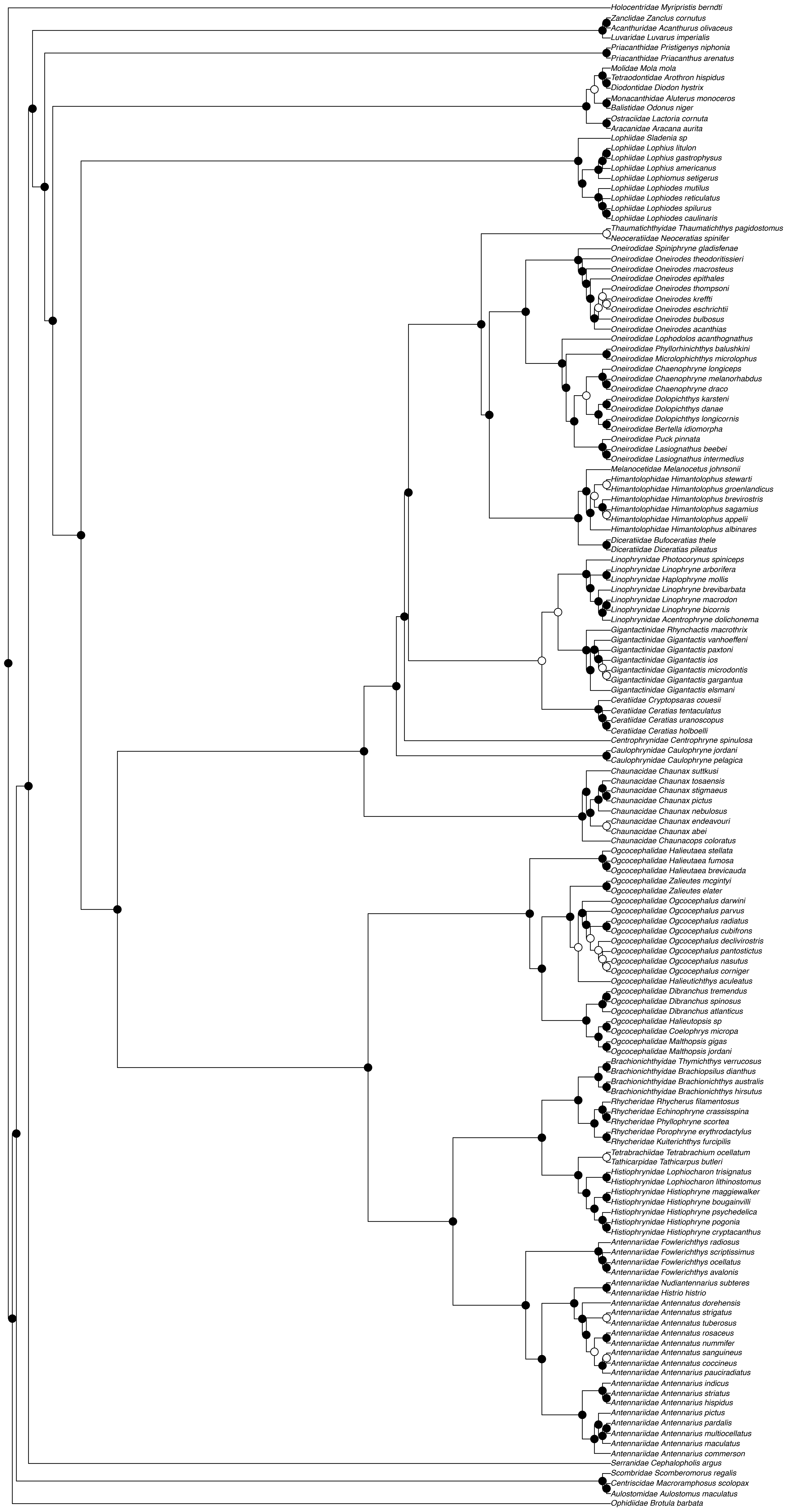

**Figure A5:** RelTime master tree with 95% CI (IQ-TREE calibrated with RelTime using †Plectocretacoidea) For the master MCMCTree see Fig. 1.

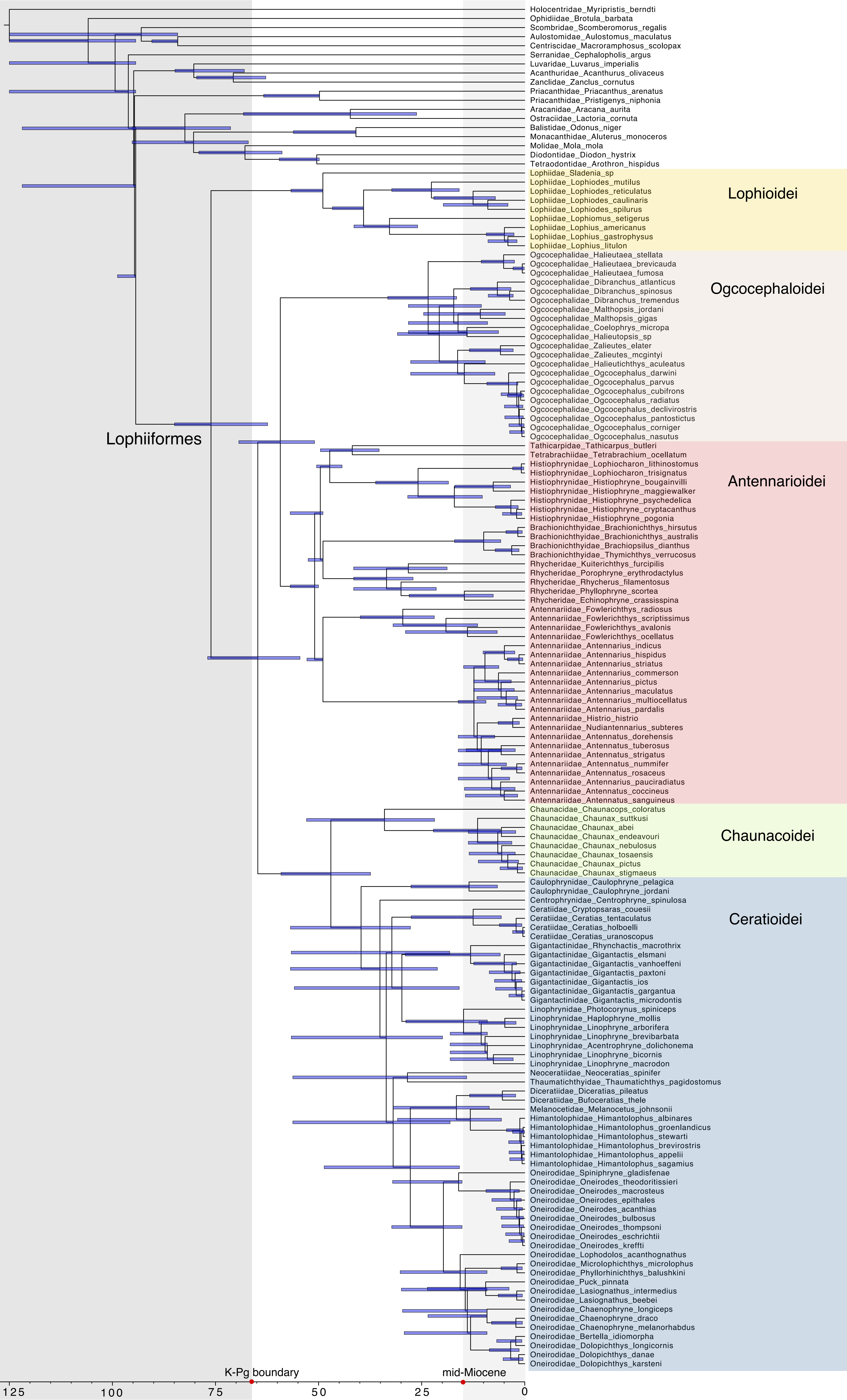
