## Supplementary Information for "Phylogenomics reveals the deep ocean as an accelerator for evolutionary diversification in anglerfishes"

**Appendix A1: Additional phylogeny figures (*separate document*)**

Figure A1: All-individuals phylogram
Figure A2: Final IQ-TREE phylogram
Figure A3: Final ASTRAL phylogram
Figure A4: IQ-TREE vs ASTRAL comparison
Figure A5: Master RelTime tree with 95% confidence intervals

***In this document:***

**Appendix A2: Systematic results and discussion**

**Appendix A3: Fossil calibration report**

Table A1: Outgroup sequence on the root
Table A2: Geologic calibration
Table A3: List of MCMCTree priors

**Appendix A4: divergence times results and discussion**

Table A4: Summary of major nodes with 95% intervals
Table A5: Comparison of fossil calibrations across other studies
Figure A6: Comparison of ages across studies and compatibility with fossil record

**Table S1:** Voucher information and quality control results for new exon data
**Table S2:** Information for UCE-mined individuals and quality control results
**Table S3:** Information for legacy marker individuals
**Table S4:** Habitat codes
**Table S5:** BioGeoBEARS model fits
**Table S6:** MiSSE model fits
**Table S7:** CT scan specimen information
**Table S8:** Disparity analyses by suborder
**Table S9:** Disparity analyses by habitat

**Figure S1:** MiSSE tip rates by tree and after pruning for taxonomic inflation
**Figure S2:** MiSSE results comparing master MCMCTree and RelTime
**Figure S3:** Body shape measurements
**Figure S4:** CT scan landmarks
**Figure S5:** Disparity by species richness
**Figure S6:** Disparity through time
**Figure S7:** PhyloEM results
**Figure S8:** BayesTraits comparing RelTime and MCMCTree, and tip rates by suborder

**Appendix A2: Systematic results and discussion**

Here we discuss our systematic results in detail with comparison to past molecular and morphological studies of Lophiiformes. We emphasize those relationships that are stable across our own analyses (concatenated and coalescent trees) and are also stable across prior molecular studies. All phylograms are in Appendix A1; Fig. 1 depicts the master MCMCTree.

Pietsch[1] is credited with the five-suborder rearrangement of Lophioidei, Antennarioidei, Ogcocephaloidei, Chaunacoidei, and Ceratioidei. Unlike classification within many percomorph clades[2], the suborders of Lophiiformes have continued to be recovered across numerous morphological and molecular studies, remaining stable for almost 40 years. A recent study proposed relegating Lophiiformes as a single suborder Lophioidei, and the Ceratioidei as the superfamily Ceratioidea[3]. We do not follow this classification, instead preferring the stable taxonomy of Pietsch[1].

**Lophioidei (Lophiidae):** Consistent with all prior studies (with exception of Shedlock et al. [4]), we find that Lophiidae is sister to all other lophiiforms. The early-branching position of Lophiidae has been long recognized from morphology[1,5].

Within Lophiidae, *Sladenia* is sister to all other lophiids. *Lophiomus* is sister to *Lophius,* and this clade is sister to *Lophiodes.* These results are identical to analyses based on morphological characters[6,7].

**Ogcocephaloidei (Ogcocephalidae):** The position of ogcocephalids within Lophiiformes has been controversial across molecular studies. We found that the family Ogcocephalidae is sister to the antennarioids[3,8,9]. Two of the cited studies are at the phylogenomic-scale, suggesting a large number of genes are needed to gain enough phylogenetic information along the base of Lophiiformes. Reticulate evolution may also be responsible for this uncertainty[9]. Alternative positions of Ogcocephalidae across past studies include the sister to the clade formed by Antennarioidei, Chaunacoidei, and Ceratioidei[10–12], and sister to the clade formed by Chaunacoidei and Ceratioidei[2].

Unlike molecular studies, morphological analyses suggest ogcocephalids are sister to the bathypelagic Ceratioidei instead of Chaunacidae, on the basis of three synapomorpies[1,13]. Nevertheless, Bertelsen[14] was open-minded about whether the sister to ceratioids were the ogcocephalids or the chaunacids, given that both occupy benthic deep-sea habitats.

Within ogcocephalids, we find that *Halieutaea* is sister to all other ogcocephalids[10]. A deep-sea clade is formed by *Dibranchus, Halieutopsis, Coelophrys,* and *Malthopsis.* It is sister to the clade formed by *Halieutichthys, Zalieutes,* and *Ogcocephalus,* with the latter two genera being predominately found on the continental shelf. Most recent molecular studies had poor sampling of ogcocephalids, with the exception of Derouen et al. [8] who sampled all genera. In their study of two mitochondrial and three nuclear genes, they also recovered the three major clades described here (*Halieutaea* as sister to all other ogcocephalids, with shallow and deep-sea sister clades). They also sampled the deep-sea *Solocisquama* and *Halicmetus* and found them to be within the deep-sea clade. The relationships among genera within these clades varies across our IQ-TREE and ASTRAL analyses and also in comparison to Derouen et al. [8], suggesting this is an area of uncertainty. Note that compared to molecules, morphological characters produce a very different set of relationships within Ogcocephalidae, with the exception of *Halieutaea*[15].

Hunt [16] showed using UCEs that the species *O. declivirostris, O. corniger, O. cubifrons, O. pantostictus,* and *O. radiatus* were not monophyletic and had extremely short branch lengths among them. She suggested that the species boundaries are in need of revision and diagnostic characters for valid species need to be re-evaluated. Similarly, we find very short branch lengths within the Atlantic *Ogcocephalus*, and non-monophyly of the species *O. parvus* and *O. declivirostris* (Appendix A1, Figure A1). Our *Ogcocephalus* samples were impossible to identify using DNA barcoding, with barcodes potentially matching several species (Table S1). For these reasons, we performed additional analyses with the genus *Ogcocephalus* pruned to avoid artificial inflation of speciation rates (see below).

**Antennarioidei:** The suborder Antennarioidei used to contain four nominal families: Antennariidae, Brachionichthyidae, Tetrabrachiidae, and Lophichthyidae. Unfortunately, the family Lophichthyidae is the only family of lophiiforms not sampled in this study or any other molecular phylogeny at time of writing. The non-monophyly of Antennariidae had been suspected since the first molecular phylogeny of Lophiiformes[4], in which the authors considered whether *Tathicarpus* should be recognized as its own family pending improved taxonomic sampling. This was finally accomplished 18 years later by Hart et al. [9], who solved this issue by splitting Antennariidae into four monophyletic families. Our results based on exons support their revisions, and so we use their revised family circumscriptions in this study.

Much prior molecular work has elucidated the relationships within frogfishes[9,11,17,18]. Our results are largely in agreement with these studies. We found two sister clades within the Antennarioidei: the family Antennariidae *sensu stricto* (corresponding to the former subfamily Antennariinae), and the clade formed by the families Rhycheridae, Brachionichthyidae,

Histiophrynidae, Tathicarpidae, and Tetrabrachiidae (including members of the former antennariid suborder Histiophryninae). The former clade is circumtropical and are broadcast spawners, while the latter clade is restricted to the Indo-West Pacific, Australia and Tasmania and undergo direct development[18]. Stem brachionichthyid fossils are known from Monte Bolca, well-nested within the Indo-Pacific frogfish clade[19]. This implies that the clade actually originated in the East Atlantic and migrated to the Indo-Pacific upon tectonic rearrangement[20].

Within the antennariids, *Fowlerichthys* is sister to the remaining genera. We found a long branch between *Fowlerichthys* and the other genera, which has been noted previously[17]. The genus *Antennarius* is sister to the clade formed by *Antennatus, Histrio,* and *Nudiantennarius.* Note that we found *Antennarius pauciradiatus* to be nested within *Antennatus,* causing its non-monophyly. Both *A. pauciradiatus* and *A. randalli* are neotenic species that have been difficult to place[17,18]. The tissue of *A. randalli* we acquired was later found to be a misidentified *Antennatus nummifer* (Table S1). The placements of the neotenic species deserves further scrutiny. It should also be noted that three specimens of *A. nummifer* that we sampled were found to be completely unrelated (Appendix A1, Figure A1), and the placement of this species remains unclear. Finally, one specimen of *A. commerson* sampled by Hart et al. [9] was re-examined and re-identified as *A. pictus* (Table S1, Appendix A1, Figure A1). The species *A. commerson* was found to be paraphyletic in their study, but a close relationship between *A. commerson, A. pictus, A. pardalis, A. multioccelatus,* and *A. maculatus* was predicted by Arnold and Pietsch[17], providing a solution to the non-monophyly found by Hart et al. [9].

Within the Indo-West Pacific frogfishes, we found that Rhycheridae and Brachionichthyidae are sister families, and together they are sister to the clade formed by Histiophrynidae, Tetrabrachiidae, and Tathicarpidae. We confirm the suspicion of Hart et al. [9]

that *Kuiterichthys* belongs in the family Rhycheridae. We found disagreement among our analyses in the relationship between Tetrabrachiidae and Tathicarpidae, where in the IQtree they are sister families (with ultrafast bootstrap support = 100), and in the ASTRAL Tetrabrachiidae is sister to Histiophrynidae (though this relationship has poor quartet support). The IQtree topology is in agreement with the UCE tree of Hart et al. [9]. Notably, the splits network analysis performed by Hart et al. [9] detected reticulation between these two families, which contributes to the uncertainty in their placement.

Unfortunately, the brachionichthyid *Sympterichthys* was the first (and only at time of writing) marine ray-finned fish to be declared extinct by the IUCN. This genus was only known from the formalin-fixed holotype[21]. Only four of the remaining 13 species of brachionichthyids are sampled in our phylogeny. Of the missing species, five are Critically Endangered or Endangered[21].

**Chaunacoidei (Chaunacidae):** We confidently find that the chaunacids are the sister group to the bathypelagic Ceratioidei, in agreement with all other molecular studies to date[2–4,8–12]. Interestingly, Bertelsen[14] believed that the pelagic larvae of chauancids gave insight into the transition from benthic to pelagic habitats by the ceratioids, hypothesizing that the pelagic stage persisted for increasingly longer durations into the adult lifespan.

**Ceratioidei:** The family-level relationships within Ceratioidei have historically been difficult to resolve. This is due to the potential confluence of rapid radiation along the backbone[4,9,10], introgression [9], poor taxonomic sampling due to the rarity of material[2,4,9,22], and the lack of reliable morphological synapomorphies due to homoplasy and the reduction of skeletal elements[10,14]. It should be noted that analyses based on morphological characters produce a very different set of relationships within the ceratioids[13] than those based on molecular data (see below), but the reliability of these characters has since been challenged[10].

By reviewing our results and other studies, we suggest that some relationships are now well-resolved, while others are still problematic. The following results are stable across our analyses and show precedence from past studies: Caulophrynidae is the sister to all other ceratioids[3,4,10], which had been suggested since Bertelsen[14,23] from morphology. While Hart et al. [9] found that Melanocetiidae was an early-branching family and Caulophrynidae was sister to Himantolophidae, we found strong evidence suggesting that a *Caulophryne* sample they used was a misidentified melanocetid based on CO1 barcoding and the position of their sample in our tree (Table S1; Appendix A1, Figure A1). A result unique to our study is that Centrophrynidae is the successive sister to the remaining ceratioids exclusive of Caulophrynidae. A relatively nested clade is well-resolved: Oneirodidae is sister to a clade formed by Diceratiidae, Melanocetidae, and Himantolophidae. This clade has been recognized since the first molecular studies[2–4,10–12,22]. The clade formed by Himantolophidae, Melanocetidae, and Diceratiidae is supported by one morphological character (ventromedial extensions of the frontal that make no contact with the parasphenoid)[13]. It is now accepted that *Lasiognathus* is in the family Oneirodidae instead of Thaumatichthyidae[3,10,11,24], meaning that the bizarre "wolftrap" phenotype has evolved twice[25]. The placement of *Lasiognathus* in Oneiroididae is hinted by several morphological characters not shared with *Thaumatichthys*[13,26]. Within Oneirodidae, the genus *Bertella* is nested within *Dolopichthys* and may be in need of taxonomic revision[27,28].

The positions of the remaining ceratioid families have been unstable across prior studies, and unfortunately remain uncertain in our analyses (Appendix A1, Figure A4). In the IQ-TREE,

Gigantactinidae is sister to Linophrynidae[22], and Neoceratiidae is sister to Thaumatichthyidae[11]. In the ASTRAL, these families are successive sister to the clade formed by Oneirodidae, Diceratiidae, Melanocetiidae, and Himantolophidae. The position of Ceratiidae is also unstable. Among the arrangements found by past studies were that Ceratiidae is sister to Gigantactinidae[11], Linophrynidae is an early-diverging ceratioid[4,11], Linophrynidae is sister to Thaumatichthyidae[10], and Linophrynidae is sister to Ceratiidae[3,9]. Nevertheless, it appears likely across all studies that obligate sexual parasitism (found in Ceratiidae, Neoceratiidae, and Linophrynidae) evolved more than once[4,10,13,22].

Interestingly, we found very short branch lengths in the genus *Himantolophus,* and we could not use DNA barcoding to confirm their species identification (Table S1). This has been noted before. Bañon et al. [27] evaluated the utility of DNA barcodes, and found that five individuals identified to four different morphological species actually belonged to the same species based on barcodes. Kai et al. [29] found "remarkably low intraspecific genetic variation" of CO1 across *Himantolophus* even with dense taxonomic sampling. It remains unclear whether the lack of a barcode gap is due to rapid speciation, low mutation rates, rampant misidentification, or taxonomic inflation. On the chance that species are overdescribed based on escal morphology[29], we chose to perform additional analyses with *Himantolophus* pruned to avoid artificially inflated speciation rates. This is the second lophiiform genus for which we made this decision (along with *Ogcocephalus*).

Surprisingly, there are several fossilized ceratioids[30–32] (Appendix A3). All belong to living genera, and the oldest are from the Miocene. Since no fossils have been found for stem genera or families, we still lack a direct view into the early evolution of ceratioids.

**Note on the potential for taxonomic inflation:** The over-description of species based on morphology (when morphology corresponds poorly to molecular divergence) may artificially inflate speciation rate estimates. As noted above, the literature shows evidence of taxonomic inflation for the Atlantic *Ogcocephalus* and *Himantolophus.* In our analyses, this manifested as extremely short branch lengths in the phylogram, non-monophyly of species, and difficulty using DNA barcoding to identify species (Appendix A1, Figure A1; Table S1). For these two genera, we pruned the genus down to two species (to retain the crown) for a set of alternative comparative analyses. We considered doing the same for the genera *Chauanx, Gigantactis,* and *Oneirodes* for which the same symptoms appeared. We ultimately chose to leave them intact because we found no precedent in the literature for concern over species boundaries or diagnostic characters. Bañon et al. [27] believed that *Gigantactis vanhoeffeni* had cryptic species, but this is the opposite of taxonomic inflation (fewer species are described than really exist). Nevertheless, we recommend a detailed molecular evaluation of all five genera using improved taxonomic sampling than the present study.

### Appendix A3: Fossil Calibration Report

Herein, Calibrations 1–8 pertain to outgroups, 9–18 are fossils pertaining to Lophiiformes, and 19–21 are geologic calibrations related to the Isthmus of Panama.

#### Outgroup calibrations:

1. Root (Total Group Holocentridae)

   MRCA: *Myripristis berndti* & *Antennatus coccineus*

   Fossil: †*Stichocentrus liratus* NHMUK PV P.47835
   Stem holocentrid [33]

   Notes: We follow Friedman et al. [34] and Hughes et al. [35] in using this fossil to calibrate the node representing the MRCA between Holocentridae and all remaining acanthomorphs.

   Min. age = 94.0 Ma, Hjoûla Lagerstätten, Lebanon [36]

   95% soft upper (Hedman method) = 125.37698

   Outgroup age sequence = 247.1, 193.81, 193.81, 181.7, 164.9, 151.2, 150.89, 103.13, 98.0, 94.0

The Outgroup sequence for Acanthomorpha is based on [34,37,38]:
247.1 Ma, Holostei, †*Watsonulus eugnathoides*;
236.0 Ma, †*Prohalecites porroi*;
221.0 Ma, †*Pholidophoridae*, †*Knerichthys bronni*;
193.81 Ma, †*Dorsetichthys bechei*;
181.7 Ma, †*Leptolepis coryphaenoides*;
166.1 Ma, †Ichthyodectiformes, †*Occithrissops willsoni*;
151.2 Ma, Elopomorpha, †*Anaethalion zapporum*;
150.94 Ma, Otocephala, †*Tischlingerichthys viholi*;
150.94 Ma, non-eurypterygian Euteleostei †*Leptolepides haerteisi*;
125.0 Ma, Aulopiformes, †*Atolvorator longipectoralis*;
98.0 Ma, Lampridiformes, †*Aipichthys minor*;

The Hedman method requires a maximum hard bound; for this bound we used †*Discoserra* (322.8 Ma) representing a probable stem neopterygian.

**Table A1:** Outgroup sequence used to place bounds on the root.

| Study | Mean age Root (Ma) | 95% HPD | Mean age Eupercaria (Ma) | 95% HPD |
|---|---|---|---|---|

| | | | | |
|---|---|---|---|---|
| Alfaro et al. [38] | 124 | (112-136) | 76 | (68-83) |
| Betancur-R et al. [39] | 141 | (125-158) | 105 | (95-115) |
| Betancur-R et al. [2] | 145 | - | 105 | - |
| Chen et al. [40] | 125 | (114-136) | 96 | (80-100) |
| Ghezelayagh et al. [3] | 133 | - | 94 | - |
| Hughes et al. [41] | 128 | (115-125) | 96 | (93-99) |
| Near et al. [42] | 126 | (118-136) | 90 | (80-100) |
| Near et al. [43] | 126 | (118-136) | 85 | (78-92) |
| Rabosky et al. [12] | 129 | - | 113 | - |

2. Crown Syngnatharia

MRCA: *Aulostomus maculatus* & *Macroramphosus scolopax*

Fossil: †*Gasterorhamphosus zuppichinii* MCSNV Na T 877.

Orr [44] argued this is a stem lineage of a clade containing Macrorhamphosidae and Centriscidae (see also [42]).

Min. age = 83.6 Ma, "Calcari di Melissano", Porto Selvaggio, Lecce Province, Italy.

Notes: Previous studies used a much more recent minimum age (~69 Ma) [34]. Santaquiteria et al. [45] reviewed new evidence that supports an older age of †*Gasterorhamphosus,* suggesting that past studies underestimated the age of Syngnatharia.

95% soft upper (Hedman method) = 110.65666

Outgroup age sequence = 247.1, 193.81, 193.81, 181.7, 164.9, 151.2, 150.89, 103.13, 98.0, 94.0

3. Total group Luvaridae (Crown Acanthuriformes *sensu* [2])

MRCA: *Luvarus imperialis* & *Acanthurus olivaceus*

Fossil: †*Kushlukia permira,* PIN 2179/64, fossil family Kushlukiidae.

Notes: Phylogenetic analyses by Bannikov and Tyler [46] placed Kushlukiidae as sister to Luvaridae.

Min. age = 55.3 Ma, Danatinsk Formation, Turkmenistan [47–49]

95% soft upper (Hedman method) = 105.90811

Outgroup age sequence = 247.1, 193.81, 193.81, 181.7, 164.9, 151.2, 150.89, 103.13, 98.0, 94.0, 55.3

4. Total group *Pristigenys* (Crown Priacanthidae)

MRCA: *Pristigenys niphonia* & *Priacanthus arenatus*

Fossil: †*Pristigenys substriata,* MNHN F.Bol529

Notes: Phylogenetic analyses by Carnevale et al. [50] placed *Pristigenys* as sister to *Pseudopriacanthus.* This clade was sister to a clade containing all other living priacanthid genera: (*Cookeolus, Heteropriacanthus, Priacanthus*).

Min. age = 48.5 Ma, Monte Bolca, Italy [51]

95% soft upper (Hedman method) = 105.06236

Outgroup age sequence = 247.1, 193.81, 193.81, 181.7, 164.9, 151.2, 150.89, 103.13, 98.0, 94.0, 48.5

5. *****Note: We used two alternative calibration schemes with and without the following fossil:**

Total group Tetraodontiformes

MRCA: *Mola mola* & *Antennatus coccineus*

Fossil: †*Plectocretacicus clarae,* MCSV S.L.1/2

Notes: According to phylogenetic analyses by Arcila and Tyler [52], the extinct †Plectocretacoidea is either sister to Lophiiformes+Tetraodontiformes or sister to Tetraodontiformes. Their position as stem Tetraodontiformes is supported by 14 synapomorphies. However, this position has been questioned by other authors, who instead believe it to be *incertae sedis* in Acanthomorpha (i.e. [38]). This fossil has potentially the greatest consequence on molecular dating of Lophiiformes, given the old age of the fossil and relationship to its sister group Tetraodontiformes [53].

Min. age = 94 Ma, Hakel, Lebanon [53–55]

95% soft upper (Hedman method) = 112.78038

Outgroup age sequence = 247.1, 193.81, 193.81, 181.7, 164.9, 151.2, 150.89, 103.13, 98.0, 94.0, 94.0

6. Total group Diodontidae+Tetraodontidae (Tetraodontoidei sensu [52])

MRCA: *Mola mola* & *Arothron hispidus*

Fossil: †*Balkaria histiopterygia*, PIN 5314/2. Fossil family Balkariidae

Notes: Phylogenetic analyses by Arcila and Tyler [52] placed this fossil as sister along the stem of Diodontidae. However, Bannikov et al. [56] suggested it is sister to (Diodontidae, Tetraodontidae). We follow Hughes et al. [35] by using the more conservative placement suggested by Bannikov et al. (2017).

Min. age = 55.8 Ma, Kheu River Formation, Russia [52,56]

Values in scheme including †Plectocretacoidea:
95% soft upper (Hedman method) = 98.29550
Outgroup age sequence = 247.1, 193.81, 193.81, 181.7, 164.9, 151.2, 150.89, 103.13, 98.0, 94.0, 94.0, 55.8

Values in scheme excluding †Plectocretacoidea:
95% soft upper (Hedman method) = 105.77898
Outgroup age sequence =247.1, 193.81, 193.81, 181.7, 164.9, 151.2, 150.89, 103.13, 98.0, 94.0, 55.8

7. Total group Tetraodontidae

MRCA: *Diodon hystrix* & *Arothron hispidus*

Fossil: †*Eotetraodon tavernei*, MCSNV IGVR 81994.

Notes: Phylogenetic analyses by Arcila and Tyler [52] placed this fossil along the stem of Tetraodontidae.

Min. age = 48.5 Ma, Monte Bolca Pesciara, Italy [51]

Values in scheme including †Plectocretacoidea:
95% soft upper (Hedman method) = 85.84214
Outgroup age sequence = 247.1, 193.81, 193.81, 181.7, 164.9, 151.2, 150.89, 103.13, 98.0, 94.0, 94.0, 55.8, 48.5

Values in scheme excluding †Plectocretacoidea:
95% soft upper (Hedman method) = 89.41161

Outgroup age sequence = 247.1, 193.81, 193.81, 181.7, 164.9, 151.2, 150.89, 103.13, 98.0, 94.0, 55.8, 48.5

8. Total group Balistidae

MRCA: *Odonus niger* & *Aluterus monoceros*

Fossil: †*Gornylistes prodigiosus*, PIN 4425/95

Notes: This is the earliest record of Balistidae [57]. Phylogenetic analyses by Arcila and Tyler [52] confirmed this is a stem balistid.

Min age = 38.4 Ma, Bartonian Stage, Gornyi Luch, Kuma Horizon, Caucasus, Russia [58]

Values in scheme including †Plectocretacoidea:
95% soft upper (Hedman method) = 96.76036
Outgroup age sequence = 247.1, 193.81, 193.81, 181.7, 164.9, 151.2, 150.89, 103.13, 98.0, 94.0, 94.0, 38.4)

Values in scheme excluding †Plectocretacoidea:
95% soft upper (Hedman method) = 103.87748
Outgroup age sequence = 247.1, 193.81, 193.81, 181.7, 164.9, 151.2, 150.89, 103.13, 98.0, 94.0, 38.4)

**Ingroup fossil calibrations:**

9. Total group Lophiidae (Crown Lophiiformes)

MRCA: *Sladenia sp.* and *Antennatus coccineus*

Fossil: †*Sharfia mirabilis* (articulated skeleton), MNHN Bol 38-39 [59]

Notes: Diagnostic characters unambiguously place †*Sharfia* in Lophiidae. It is distinct from all other lophiids by its triangular opercle and non-fimbriate subopercle (a state also found in some frogfishes, batfishes, and chaunacids). Phylogenetic analyses by Carnevale and Pietsch [7] using a matrix of 38 characters show that †*Sharfia* is sister to all other known lophiids based on one synapomorphy (opercle strongly bifurcate). †*Caruso* and †*Sharfia* are together the oldest known fossil representatives of Lophiidae. Since Lophiidae is sister to all other Lophiiformes, this fossil is effectively a calibration on Crown Lophiiformes.

Min. age = 48.5 Ma, late early Eocene, Ypresian; Monte Bolca, Pesciara site [51]

Values in scheme including †Plectocretacoidea:

95% soft upper (Hedman method) = 97.64885
Outgroup age sequence = 247.1, 193.81, 193.81, 181.7, 164.9, 151.2, 150.89, 103.13, 98.0, 94.0, 94.0, 48.5

Values in scheme excluding †Plectocretacoidea:
95% soft upper (Hedman method) = 105.06236
Outgroup age sequence = 247.1, 193.81, 193.81, 181.7, 164.9, 151.2, 150.89, 103.13, 98.0, 94.0, 48.5

10. Total group *Sladenia* (Crown Lophiidae)

MRCA: *Sladenia sp.* and *Lophiodes spilurus*

Fossil: †*Caruso brachysomus* (articulated skeleton), MNHN Bol 42/43 [7].

Notes: Phylogenetic analyses by Carnevale and Pietsch [7] using a matrix of 38 characters show that the fossil †*Caruso* is sister to extant *Sladenia* based on two synapomorphies. This places it within Crown Lophiidae. †*Caruso* and †*Sharfia* are together the oldest known fossil representatives of Lophiidae.

Min. age = 48.5 Ma, late early Eocene, Ypresian; Monte Bolca, Pesciara site [51]

Values in scheme including †Plectocretacoidea:
95% soft upper (Hedman method) = 84.74384
Outgroup age sequence = 247.1, 193.81, 193.81, 181.7, 164.9, 151.2, 150.89, 103.13, 98.0, 94.0, 94.0, 48.5, 48.5

Values in scheme excluding †Plectocretacoidea:
95% soft upper (Hedman method) = 88.58789
Outgroup age sequence = 247.1, 193.81, 193.81, 181.7, 164.9, 151.2, 150.89, 103.13, 98.0, 94.0, 48.5, 48.5

11. *Lophius* + *Lophiodes*

MRCA: *Lophius litulon* & *Lophiodes spilurus*

Fossil: †*Eosladenia caucasica* (complete, partially disarticulated skeleton), PIN 4425–72 [60]

Notes: Phylogenetic analyses by Carnevale and Pietsch [7] using a matrix of 38 characters show that the clade (†*Eosladenia*,(*Lophiomus*,*Lophius*)) is sister to *Lophiodes.* The sister group relationship between †*Eosladenia* and *Lophiomus* + *Lophius* is supported by one synapomorphy (caudal centrum depressed, bearing lateral transverse processes). This places the fossil well within crown Lophiidae.

Min. age = 38.4 Ma [58], Bartonian Stage, Gornyi Luch, Russia [60]

Values in scheme including †Plectocretacoidea:
95% soft upper (Hedman method) = 69.43063
Outgroup age sequence = 247.1, 193.81, 193.81, 181.7, 164.9, 151.2, 150.89, 103.13, 98.0, 94.0, 94.0, 48.5, 48.5, 38.4

Values in scheme excluding †Plectocretacoidea:
95% soft upper (Hedman method) = 71.70811
Outgroup age sequence = 247.1, 193.81, 193.81, 181.7, 164.9, 151.2, 150.89, 103.13, 98.0, 94.0, 48.5, 48.5, 38.4

12. Total group Ogcocephalidae

MRCA: *Zalieutes elater* & *Antennatus coccineus*

Fossil: †*Tarkus squirei*, MCSNV T158/T159 [61]. Taxon is based on five skeletons including one complete articulated skeleton.

Notes: The fossil is diagnosed as an ogcocephalid based on the shape of the illicium, tubercles on the head and body, depressed body shape, horizontal gape of the mouth, and several skeletal and meristic characters [61]. Its relationship to living ogcocephalid genera is unclear. This is hampered in part because the relationships among living genera are also unclear [15]. It is morphologically unique from all other ogcocephalids, though an affiliation with *Halieutaea* and *Halieutichthys* was suggested by Carnevale and Pietsch [61]. This fossil is the earliest known record, as well as the first articulated skeletal record of Ogcocephalidae. Placing this calibration at the node corresponding to stem Ogcocephalidae is conservative.

Min. age = 48.5 Ma, Monte Bolca Pesciara, Italy [51]

Values in scheme including †Plectocretacoidea:
95% soft upper (Hedman method) = 84.74384
Outgroup age sequence = 247.1, 193.81, 193.81, 181.7, 164.9, 151.2, 150.89, 103.13, 98.0, 94.0, 94.0, 48.5, 48.5

Values in scheme excluding †Plectocretacoidea:
95% soft upper (Hedman method) = 88.58789
Outgroup age sequence = 247.1, 193.81, 193.81, 181.7, 164.9, 151.2, 150.89, 103.13, 98.0, 94.0, 48.5, 48.5

13. Crown Antennarioidei, or Total group Antennariidae *sensu stricto* [9]

MRCA: *Antennatus coccineus* (Antennariidae) & *Kuiterichthys furcipilis* (Rhycheridae)

Fossil: †*Eophryne barbutii,* MCSNV B.6513, articulated skeleton [62]

Notes: Identification as an antennariid on the basis of: general physiognomy of the body, globose shape, mouth large and oblique, simple pectoral fin, sigmoid vertebral column, meristic value of dorsal and anal fins.

Hart et al. [9] broke the former Antennariidae into four families, where Antennariidae is now restricted to the former subfamily "Antennariinae". Carnevale and Pietsch [62] noted similarity of †*Eophryne* to the living genera *Antennarius* (which was not considered distinct from *Fowlerichthys* at the time of their study), *Histrio*, and *Nudiantennarius*. All three of these genera remain in the restricted circumscription of Antennariidae *sensu* Hart et al. [9]. †*Eophryne* has an epural present, which is a diagnostic character of "Antennariinae", though it lacks the other diagnostic character of an endopterygoid being present [63]. This provides justification that †*Eophryne* is at least a crown antennarioid, and (conservatively) a stem antennariid. Past molecular studies instead used †*Eophryne* to calibrate Total group Antennarioidei, whether as a deliberately conservative choice [3] or erroneously [12].

Min. age = 48.5 Ma, Ypresian of Monte Bolca, Pesciara site [51]

Values in scheme including †Plectocretacoidea:
95% soft upper (Hedman method) = 71.56426
Outgroup age sequence = 247.1, 193.81, 193.81, 181.7, 164.9, 151.2, 150.89, 103.13, 98.0, 94.0, 94.0, 48.5, 48.5, 48.5

Values in scheme excluding †Plectocretacoidea:
95% soft upper (Hedman method) = 73.76086
Outgroup age sequence = 247.1, 193.81, 193.81, 181.7, 164.9, 151.2, 150.89, 103.13, 98.0, 94.0, 48.5, 48.5, 48.5

14. Total group *Fowlerichthys,* or Crown Antennariidae *sensu* Hart et al. [9] (formerly called Antennariinae)

MRCA: *Fowlerichthys avalonis* (Antennariidae) & *Antennatus coccineus* (Antennariidae)

Fossil: †*Neilpeartia ceratoi,* CMC 3, a nearly complete and well-preserved articulated skeleton [63]

Notes: While Histiophryninae had not yet been separated from Antennariidae at the time this fossil was published [9], Carnevale et al. [63] pointed out that †*Neilpeartia* has diagnostic features of "Antennariinae" (now Antennariidae *sensu stricto*): both endopterygoid and epural present. Note that the epural is also present in †*Eophryne* (our calibration #13). A sister group relationship with the extant genus *Fowlerichthys* is supported by the five distally branched pelvic fin rays. This would place †*Neilpeartia* in crown "Antennariinae". *Fowlerichthys* is the sister to all other "antennariines", and is separated from them by a long internal branch [17] (Fig. 1). Therefore, this calibration is justifiable for crown "Antennariinae" or Antennariidae *sensu stricto* [9]. This represents the "oldest known unquestionable evidence of crown antennariids in the fossil record" [63]. An alternative, more conservative placement of this calibration would be at Total group "Antennariinae", which would make it redundant with †*Eophryne* (our calibration #13).

Min. age = 48.5 Ma, Ypresian of Monte Bolca, Pesciara site [51]

Values in scheme including †Plectocretacoidea:
95% soft upper (Hedman method) = 61.95415
Outgroup age sequence = 247.1, 193.81, 193.81, 181.7, 164.9, 151.2, 150.89, 103.13, 98.0, 94.0, 94.0, 48.5, 48.5, 48.5, 48.5

Values in scheme excluding †Plectocretacoidea:
95% soft upper (Hedman method) = 63.32703
Outgroup age sequence = 247.1, 193.81, 193.81, 181.7, 164.9, 151.2, 150.89, 103.13, 98.0, 94.0, 48.5, 48.5, 48.5, 48.5

15. Total group Brachionichthyidae

MRCA: *Brachionichthys australis* (Brachionichthyidae) & *Kuiterichthys furcipilis* (Rhycheridae)

Fossil: †*Histionotophrous bassani,* MGPD 68487, and †*Orrichthys longimanus,* MCSNV T.160/161. †*Histionotophrous* is known from at least a dozen specimens including complete skeletons. †*Orrichthys* is based on two nearly complete skeletons.

Notes: Pietsch [64] suggested the fossil †*Histionotophrous* was a brachionichthyid instead of an antennariid, and even supposed it could be synonymous with the living genus *Brachionichthys.* Phylogenetic analysis by Carnevale and Pietsch [19] using a matrix of 36 characters found †*Histionotophrous* and †*Orrichthys* to be sister genera. This clade is sister to the living brachionichthyid genera *Brachionichthys* and *Sympterichthys.* This analysis confirms that both fossil genera can be classified as stem brachionichthyids.

Brachionichthyidae is one of the four families of suborder Antennarioidei named prior to Hart et al. [9]. The presence of several antennarioid lineages at Monte Bolca (our calibrations #14 and #15) confirms that crown Antennarioidei itself must be at least as old as the Eocene. Past molecular studies placed hard minimum bounds on Total group Antennarioidei as Eocene [3,10,12]. No prior studies to date have applied Eocene calibrations on nodes more nested than this (Appendix A4).

Min. age = 48.5 Ma, Monte Bolca Pesciara, Italy [51]

Values in scheme including †Plectocretacoidea:
95% soft upper (Hedman method) = 61.95415

Outgroup age sequence = 247.1, 193.81, 193.81, 181.7, 164.9, 151.2, 150.89, 103.13, 98.0, 94.0, 94.0, 48.5, 48.5, 48.5, 48.5

Values in scheme excluding †Plectocretacoidea:
95% soft upper (Hedman method) = 63.32703
Outgroup age sequence = 247.1, 193.81, 193.81, 181.7, 164.9, 151.2, 150.89, 103.13, 98.0, 94.0, 48.5, 48.5, 48.5, 48.5

16. Total group *Acentrophryne* (Linophrynidae)

MRCA: *Acentrophryne dolichonema* & *Linophryne macrodon*

Fossil: *Acentrophryne sp.* LACM 117685 [31]. Fossil is a nearly-complete, moderately well-preserved skeleton of a metamorphosed female.

Notes: The fossil shares the most characters with the extant genus *Acentrophryne:* no spine on the preopercle, reduced number of teeth compared to other linophrynids, and an elongate illicium. It is unclear whether the fossil belongs to an extant species, or represents a new, extinct species. The fossil has an illicium length intermediate between the two living *Acentrophryne* species, suggesting it could be distinct from those species [31].

Min. age = 7.6 Ma, Chalk Hill, Yorba Member of the Puente Formation, Los Angeles, CA (see discussion in [30,31,65]).

Values in scheme including †Plectocretacoidea:
95% soft upper (Hedman method) = 79.222022
Outgroup age sequence = 247.1, 193.81, 193.81, 181.7, 164.9, 151.2, 150.89, 103.13, 98.0, 94.0, 94.0, 48.5, 7.6

Values in scheme excluding †Plectocretacoidea:
95% soft upper (Hedman method) = 82.692693
Outgroup age sequence = 247.1, 193.81, 193.81, 181.7, 164.9, 151.2, 150.89, 103.13, 98.0, 94.0, 48.5, 7.6

17. Crown *Chaenophryne* (Oneirodidae)

MRCA: *Chaenophryne melanorhabdus* & *Chaenophryne longiceps*

Fossil: *Chaenophryne aff. melanorhabdus* (extant), LACM 136809, 136810, 137674, 137703, articulated skeletons [30]

Notes: Specimens share features diagnostic to *Chaenophryne:* highly cancellous texture of bones, strongly convex forsolateral profile of the frontal, sphenotic spine absent, syphysial spine of the dentary absent, dentary teeth large, articular spine blunt, angular spine reduced and stout, opercle triangular with a slightly concave posterior margin, etc. The genus has five living species separated by escal morphology and meristics, which are not retained in the fossil specimen. The specimens were hypothesized to be of *C. melanorhabdus* based on body shape, opercle and subopercle shape, and anal ray count.

Min. age = 7.6 Ma, Yorba Member of the Puente Formation, Los Angeles, CA (see discussion in [30,31,65]).

Values in scheme including †Plectocretacoidea:
95% soft upper (Hedman method) = 79.222022
Outgroup age sequence = 247.1, 193.81, 193.81, 181.7, 164.9, 151.2, 150.89, 103.13, 98.0, 94.0, 94.0, 48.5, 7.6

Values in scheme excluding †Plectocretacoidea:
95% soft upper (Hedman method) = 82.692693
Outgroup age sequence = 247.1, 193.81, 193.81, 181.7, 164.9, 151.2, 150.89, 103.13, 98.0, 94.0, 48.5, 7.6

18. Total group *Oneirodes*

MRCA: *Oneirodes kreffti* & *Spiniphryne gladisfenae* (MCZ specimen)

Fossil: †*Oneirodes sp.,* ZIN 461p, a nearly complete, partly disarticulated skeleton [32]

Notes: The species identity is unclear, but fossil is assignable to *Oneirodes* on the basis of its anteriorly bifurcated frontal bones, the absence of pelvic fins and scales, presence of sphenotic spines, and meristics. This is the oldest record of *Oneirodes* in the fossil record, supplanting the 7.6 million-year-old fossil from the Puente Formation [30] used as a calibration in past molecular studies [10,12].

Min. age = 13.1 Ma, Kurasi Formation, Sakhalin, Russia (see discussion and references in [66,67])

Values in scheme including †Plectocretacoidea:
95% soft upper (Hedman method) = 79.75215
Outgroup age sequence = 247.1, 193.81, 193.81, 181.7, 164.9, 151.2, 150.89, 103.13, 98.0, 94.0, 94.0, 48.5, 13.1

Values in scheme excluding †Plectocretacoidea:
95% soft upper (Hedman method) = 83.47227
Outgroup age sequence = 247.1, 193.81, 193.81, 181.7, 164.9, 151.2, 150.89, 103.13, 98.0, 94.0, 48.5, 13.1

**Geologic calibrations:**

The final three calibrations are geologic calibrations on the age of robust sister-species pairs separated by the Isthmus of Panama. Following Rincon-Sandoval et al. [68], we use 2.8 Ma as the minimum age for these calibrations reflecting the undisputed minimum geologic age for the closure of the Isthmus of Panama.

**Table A2:** List of geologic calibrations.

| Cal. No. | Family | East Pacific species | West Atlantic species |
|---|---|---|---|
| 19 | Ogcocephalidae | *Dibranchus spinosus* | *Dibranchus tremendus* |
| 20 | Ogcocephalidae | *Zalieutes elater* | *Zalieutes mcgintyi* |
| 21 | Antennariidae | *Fowlerichthys avalonis* | *Fowlerichthys ocellatus* |

**Table A3:** Summary of all priors used in MCMCTree (units in hundreds of millions of years).

| Calib. Number | Absolute age (mya) | Distribution | Calibration type | Parameters, scheme including Plecto. | Parameters, scheme excluding Plecto. |
|---|---|---|---|---|---|
| 1 (Root) | 94 | Uniform | Soft upper and hard lower bound | B(0.94, 1.2537698) | Same |
| 2 | 83.6 | Uniform | Soft upper and hard lower bound | B(0.836, 1.1065666) | Same |
| 3 | 55.3 | Uniform | Soft upper and hard lower bound | B(0.553, 1.0590811) | Same |
| 4 | 48.5 | Uniform | Soft upper and hard lower bound | B(0.485, 1.0506236) | Same |
| 5 (Plecto) | 94 | Uniform | Soft upper and hard lower bound | B(0.94, 1.1278038) | Excluded |
| 6 | 55.8 | Uniform | Soft upper and hard lower bound | B(0.558, 0.9829550) | B(0.558, 1.0577898) |
| 7 | 48.5 | Uniform | Soft upper and hard lower bound | B(0.485, 0.8584214) | B(0.485, 0.8941161) |
| 8 | 38.4 | Uniform | Soft upper and hard lower bound | B(0.384, 0.9676036) | B(0.384, 1.0387748) |
| 9 | 48.5 | Uniform | Soft upper and hard lower bound | B(0.485, 0.9764885) | B(0.485, 1.0506236) |
| 10 | 48.5 | Uniform | Soft upper and hard lower bound | B(0.485, 0.8474384) | B(0.485, 0.8858789) |
| 11 | 38.4 | Uniform | Soft upper and hard lower bound | B(0.384, 0.6943063) | B(0.384, 0.7170811) |
| 12 | 48.5 | Uniform | Soft upper and hard lower bound | B(0.485, 0.8474384) | B(0.485, 0.8858789) |
| 13 | 48.5 | Uniform | Soft upper and hard lower bound | B(0.485, 0.7156426) | B(0.485, 0.7376086) |
| 14 | 48.5 | Uniform | Soft upper and hard lower bound | B(0.485, 0.6195415) | B(0.485, 0.6332703) |
| 15 | 48.5 | Uniform | Soft upper and hard lower bound | B(0.485, 0.6195415) | B(0.485, 0.6332703) |
| 16 | 7.6 | Uniform | Soft upper and hard lower bound | B(0.076, 0.79222022) | B(0.076, 0.82692693) |
| 17 | 7.6 | Uniform | Soft upper and hard lower bound | B(0.076, 0.79222022) | B(0.076, 0.82692693) |
| 18 | 13.1 | Uniform | Soft upper and hard lower bound | B(0.131, 0.7975215) | B(0.131, 0.8347227) |
| 19 | 2.8 | Cauchy | Hard lower bound | L(0.028) | Same |
| 20 | 2.8 | Cauchy | Hard lower bound | L(0.028) | Same |
| 21 | 2.8 | Cauchy | Hard lower bound | L(0.028) | Same |

### Appendix A4: Divergence time results and discussion

***Divergence time results:***

We produced eight alternative time trees using either the IQ-TREE (concatenated) or ASTRAL (coalescent) phylograms, the fossil calibration scheme with or without the controversial fossil Plectocretacicoidea, and using either MCMCtree or RelTime as the calibration method. The methodological decision with the largest impact on divergence times was MCMCTree versus RelTime. There was a consistent gap between MCMCtree and RelTime age estimates for important nodes in the tree, with MCMCTree often leaning older by 10–30 million years. For example, the crown age of Lophiiformes had a 31-million-year difference in estimates between the IQ-TREEs calibrated using MCMCTree versus RelTime while †Plectocretacoidea was excluded, which was reduced to a 15-million-year difference for the same node while including †Plectocretacoidea. As an example of a younger node, the age of crown Antennarioidei had a 14- and 13-million-year difference for the same two comparisons respectively. While we expected the inclusion of †Plectocretacoidea to have a major influence on divergence times, the difference was negligible when using MCMCtree. When using RelTime, only the early nodes in the tree showed a large difference when using †Plectocretacoidea. For example, the crown age of Lophiiformes differed by 15 million years when comparing IQ-TREEs calibrated using RelTime and either including or excluding †Plectocretacoidea; the corresponding difference in the crown age of Antennarioidei was only 1 million years. The difference in divergence times resulting from using the IQ-TREE versus the ASTRAL phylogram with all else equal was usually negligible.

Divergence time estimates of important nodes in the tree are summarized in Fig. 1 and Table A4. Six of the eight time trees inferred a Cretaceous origin of crown Lophiiformes (all but the RelTime trees excluding †Plectocretacoidea). The MRCA of the four suborders excluding Lophioidei was also Cretaceous in the MCMCTrees, but shortly after the K-Pg boundary in the RelTime trees. In the MCMCTrees, Ceratioidei diverged from Chauancoidei near the K-Pg boundary, whereas in the RelTime trees this split occurred well into the Eocene. Similarly, the two methods result in a >20 million-year difference in the origin of Crown Ceratioidei, either in the Paleocene (MCMCTree) or late Eocene (RelTime). A Paleocene age of Ceratioidei would be consistent with other open-ocean fishes that diversified in the wake of the K-Pg mass extinction[34,69,70].

Two caveats should be noted for the RelTime dates. First, there are few molecular substitutions (i.e. short internodes) visible in the phylogram along the backbone of Ceratioidei, and this is a condition when RelTime can underestimate divergence times[71,72]. Second, the 95% confidence intervals are much wider overall for the RelTime dates compared to the MCMCTree dates, but especially so for the backbone of Ceratioidei (Fig. 1, Appendix A1, Figure A5). Sadly, there are no known stem ceratioid fossils to use as hard calibrations in this region of the tree.

**Table A4:** Summary of divergence time estimates for important nodes (in millions of years). Point estimates are ranges from the eight time trees. The 95% CI intervals in brackets refer to those when using the IQ-TREE phylogram (for the MCMCTree column, the intervals are only when including †Plectocretacoidea). For a graphical depiction see Figure 1. For 95% confidence intervals on all nodes see Fig. 1 (MCMCTree using IQ-TREE and including †Plectocretacoidea) or Appendix A1, Figure A5 (RelTime using IQ-TREE and including †Plectocretacoidea).

| Node | MCMCTree | RelTime, including †Plectocretacoidea | RelTime, excluding †Plectocretacoidea |
|---|---|---|---|
| Stem Lophiiformes (MRCA of Lophiiformes and Tetraodontiformes) | 100 [94.9–106.7] | 94–95 [94.3–98.7] | 69–72 [57.5–103.5] |
| Crown Lophiiformes | 91–92 [85.4–97.2] | 76–78 [62.3–84.9] | 61–63 [53.6–76.0] |
| MRCA of the four suborders excluding Lophioidei | 82–83 [76.3–87.9] | 63–65 [54.5–76.9] | 56–57 [50.9–66.5] |
| Crown Lophioidei | 57 [49.3–68.8] | 49 [48.9–56.7] | 49 [48.9–57.6] |
| Crown Ogcocephaloidei | 31–32 [25.5–39.0] | 23 [16.5–33.2] | 21 [14.7–31.1] |
| Crown Antennarioidei | 63–64 [58.8–68.7] | 50 [50.0–56.8] | 51 [50.0–56.5] |
| MRCA of Chaunacoidei and Ceratioidei | 67 [58.7–75.0] | 44–47 [37.4–59.1] | 40–42 [33.6–51.4] |
| Crown Chaunacoidei | 48 [18.2–64.6] | 30–34 [21.8–52.9] | 27–30 [19.6–45.9] |
| Crown Ceratioidei | 58–59 [51.3–65.8] | 37–40 [27.7–56.8] | 34–35 [33.6–51.4] |
| Crown Oneirodidae | 31–32 [26.6–37.3] | 18–20 [15.2–32.2] | 18–19 [15.2–31.2] |

***An older Lophiiformes:***

A major result from our study is that the crown age of Lophiiformes is well within the Cretaceous. Even our trees with the youngest estimates (RelTime excluding †Plectocretacoidea) have confidence intervals extending to ~76 million years ago. Yet, recent molecular studies instead found that Lophiiformes have a Cenozoic origin, as part of a post-K-Pg diversification event affecting acanthomorphs more broadly[3,38]. Here we discuss this new result and suggest that the age of Lophiiformes was underestimated by past studies. To aid in this discussion, we compared the use of fossil calibrations and resulting divergence time estimates by seven recent molecular studies[2,3,8,10,12,38,41] to our study.

The primary reason for the generally older age estimates in our study is our use of six fossil calibrations from Monte Bolca (48.5 mya[51]). While we included a Cretaceous fossil calibration in some analyses (†Plectocretacoidea), the difference in divergence times using calibration schemes with and without this fossil was negligible for MCMCTree and limited to early nodes for RelTime. Our six Monte Bolca calibrations were placed on a range of stem and crown nodes: crown Lophiiformes, crown Lophiidae, stem Ogcocephaloidei, crown Antennarioidei, stem Brachionichthyidae, and crown Antennariidae (Fig. 1). The number of fossil calibrations of any kind differs greatly between our study and prior studies (Table A5). Two of the seven studies we examined used zero fossil calibrations for Lophiiformes[2,41]. Three studies used a single calibration within Lophiiformes[3,8,38]. The remaining studies used four fossil calibrations (three from Monte Bolca) [10] and six calibrations (three from Monte Bolca)[12] respectively.

Why did past studies use fewer calibrations? The most likely reason is that taxonomic sampling was limited in past studies, providing fewer nodes for which to place fossil calibrations. For example, Ghezelayagh et al. [3] sampled three species of antennarioids representing three of seven families; Betancur et al. [2] only sampled "antennariine" members of Antennarioidei (one family); Alfaro et al. sampled three lophiiforms in general (*Antennarius, Cryptopsaras,* and *Ogcocephalus*). Another obvious reason is that some fossils were only recently described; two of our ten ingroup fossil calibrations were described in 2020[32,63] (Table A5).

In addition to the number of calibrations, the usage of calibrations also differs between our study and past studies. Two studies placed a hard minimum bound on Total group Antennarioidei as Eocene based on the Monte Bolca fossil †*Eophryne*[3,12]. However, †*Eophryne* is a crown antennarioid (Appendix A3). Ghezelayagh et al. [3] justified their placement as the conservative choice, perhaps tacitly due to the non-monophyly of Antennariidae at the time (which has since been settled by Hart et al. [9]). Nonetheless, stem brachionichthyids are also well known from Monte Bolca[19], providing further evidence that crown Antennarioidei originated prior to Monte Bolca. Moving the calibration from stem to crown Antennarioidei will inevitably push the age of Antennarioidei older, likely affecting ancestral nodes as well.

As a consequence of these factors, the molecular divergence times for some lophiiform clades inferred by past studies are demonstrably too young. In light of Monte Bolca fossil representatives of crown Lophiidae, crown Antennarioidei, and crown Antennariidae, estimates of these nodes younger than 48.5 Ma are inconsistent with the fossil record (Fig. A6). These inconsistencies are present in some divergence times estimated by Derouen et al. [8], Betancur-R et al. [2], Rabosky et al. [12], and Ghezelayagh et al. [3]. Note that divergence time estimates by Miya et al. [10] are likely too old (e.g. Jurassic divergence for stem Lophiiformes), and are possibly driven by high substitution rates of mitochondrial DNA.

Is a Cretaceous origin of Lophiiformes reasonable, all things considered? There are no well-preserved fossils of Lophiiformes older than Monte Bolca. This means that we have no direct evidence of the origins of lophiiform clades prior to the Eocene. Yet, the presence of several lineages in Monte Bolca, including crown representatives of two suborders and stem fossils of a third suborder, strongly suggests that Lophiiformes was already well-diversified prior to Monte Bolca. Carnevale and Pietsch [62] wrote: “the origin of the various lophiiform clades must be searched for in the Paleocene or at least in the Late Cretaceous”. If Lophiiformes originated immediately after the K-Pg, this would provide only 17.5 million years for the origination of Lophioidei, Ogcocephaloidei, Antennarioidei, and the lineage that would eventually diverge into Chaunacoidei and Ceratiodei. Yet, no rapid radiation of the sort is visible in our phylograms (Appendix A1). The branch lengths leading to the suborders are not particularly short, especially compared to branches within the Ceratioidei corresponding to higher taxa within that clade. Therefore, we suggest that a Cretaceous origin of Lophiiformes is the best explanation to reconcile our molecular dataset with the fossil calibrations.

**Table A5:** Comparison of fossil calibrations across time-calibrated molecular phylogenies of Lophiiformes. Blank cells indicate the fossil was not used in that study.

| **Fossil** | **Publication date** | **Fossil age (mya)** | **Fossil taxonomy** | **Node assigned for calibration** | | | | | |
|---|---|---|---|---|---|---|---|---|---|
| | | | | **Ghezelayagh et al.** [3] | **Alfaro et al.** [38] | **Rabosky et al.** [12] ** | **Derouen et al.** [8] | **Miya et al.** [10] | **This study** |
| †*Sharfia* | 2011 | 48.5 | Stem lophiid | | | | | | Crown Lophiiformes |
| †*Caruso* | 2012 | 48.5 | Crown lophiid | | Used for outgroup sequence | Crown Lophiiformes | | Crown Lophiiformes ***** | Crown Lophiidae |
| †*Eosladenia* | 2004 | 38.4 | Crown lophiid | | | *Lophioides + Lophius* | | | *Lophiodes + Lophius* |
| †*Tarkus* | 2011 | 48.5 | Stem ogcocephalid | | Total group Ogcocephalidae | Total group Ogococephalidae | Crown Ogcocephalidae | Total group Ogococephalidae | Total group Ogcocephalidae |
| †*Eophryne* | 2009 | 48.5 | Crown antennarioid, stem antennariid | Total group Antennarioidei * | Used for outgroup sequence | Total group Antennarioidei *** | | | Crown Antennarioidei |
| †*Neilpeartia* | 2020 | 48.5 | Crown antennariid *sensu* Hart et al. | | | | | | Total group *Fowlerichthys* |

| | | | | | | | | | |
|---|---|---|---|---|---|---|---|---|---|
| | | | (formerly called Antennariinae) | | | | | | |
| †*Histionotophrous* | Revised in 2010 | 48.5 | Stem brachionichthyid | | | | | Total group Antennarioidei | Total group Brachionichthyidae |
| *Acentrophryne* | 2009 | 7.6 | Living genus | | | | | | Total group *Acentrophryne* |
| *Chaenophryne* | 2008 | 7.6 | Living genus | | | | | | Total group *Chaenophryne* |
| *Oneirodes* | 2020 | 13.1 | Living genus | | | Total group *Oneirodes* **** | | Crown Ceratioidei **** | Total group *Oneirodes* |

* Ghezelayagh et al. [3] acknowledged that this placement was conservative, despite the fossil's similarity to Antennariidae

** Rabosky et al. used six ingroup fossils. They used the fossil *Antennarius monodi*[73] to date the MRCA of *Antennarius indicus* and *A. striatus* (5.33 mya). We did not use this fossil because it is not clear if the "Antennarius ocellatus group" is a real clade. *Antennarius ocellatus* is now considered to be *Fowlerichthys ocellatus,* and *Fowlerichthys* has a very different phylogenetic placement from remaining antennariids.

*** This position of †*Eophryne* in this study was erroneous. Rabosky et al. [12] labelled the calibration as Total group Antennariidae, yet actually placed the calibration on the older node corresponding to Total group Antennarioidei.

**** A fossil discovered earlier[30] dated at 7.42 was used. This was the oldest ceratioid fossil known at the time of these studies, and has since been supplanted by a 13.1 mya fossil[32].

***** Likely refers to material misidentified as †*Lophius brachysomus* that has since been re-evaluated[7,59]

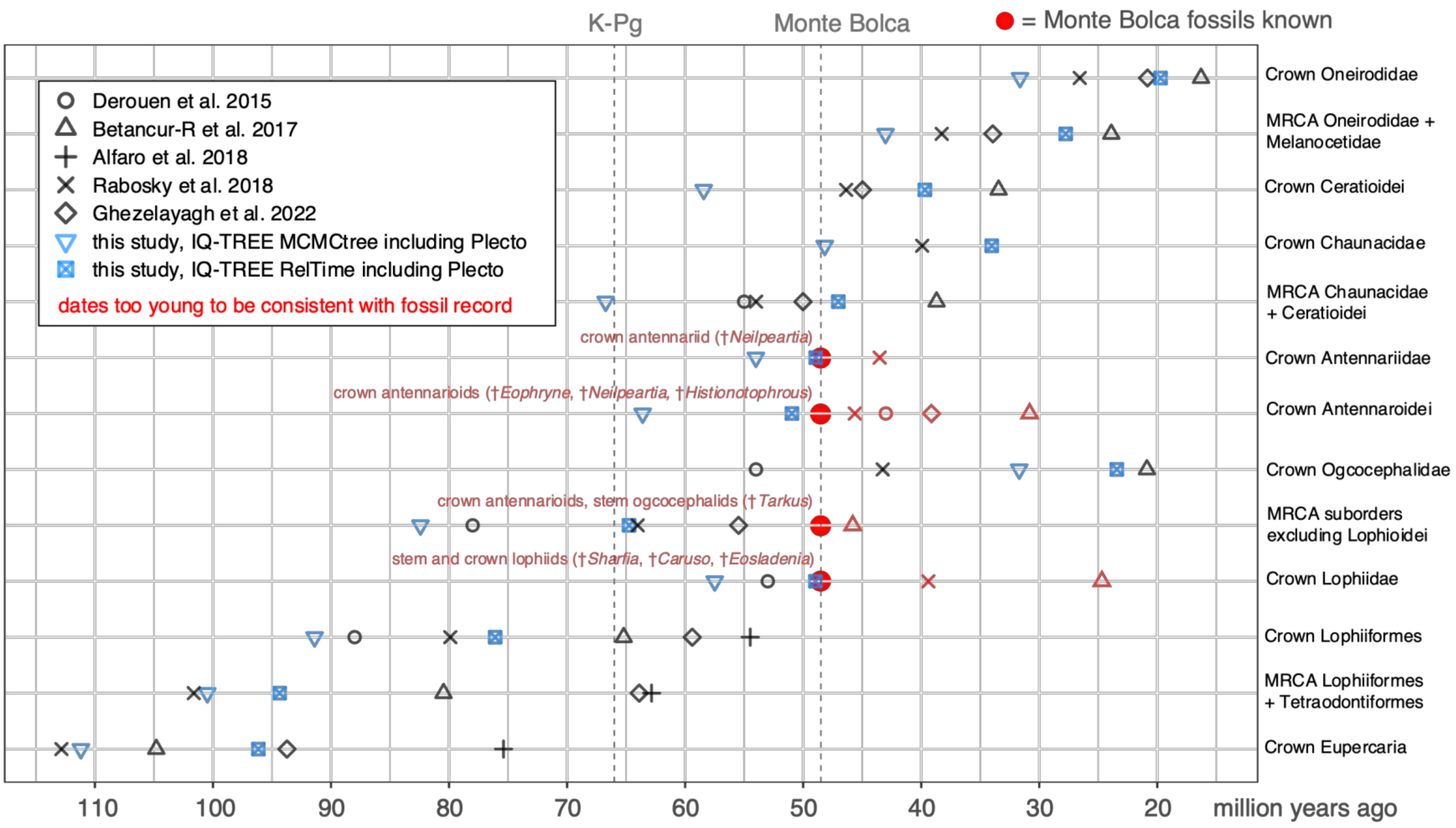

**Figure A6:** Comparing the age of important nodes from past studies to our results. Symbols colored in red represent nodes from past studies that are incompatible with the age of Monte Bolca fossils (i.e., the node must have a minimum age of 48.5 Ma to be consistent with the lineage's presence at Monte Bolca).

**Table S1.** List of tissue vouchers used to generate new genomic data in this study and results from quality control steps. Specimens likely to be misidentified are in bold.

| Species | Specimen voucher number | Tissue or biorepository number, if different | Number of exons assembled and remaining after quality control | BOLD top hit | Used in PCM trees? | Notes |
|---|---|---|---|---|---|---|
| *Antennarius commerson* | | BGI-052-053 | 842 | *Antennarius commerson* | No | |
| *Antennarius commerson* | UW 117686 | | 909 | *Antennarius commerson* | Yes | |
| *Antennarius hispidus* | UW 117828 | | 927 | *Antennarius hispidus* | Yes | |
| *Antennarius indicus* | UW 118817 | | 913 | *Antennarius indicus* | Yes | |
| *Antennarius maculatus* | UW 117687 | | 946 | *Antennarius maculatus* | Yes | |
| *Antennarius multiocellatus* | USNM 414303 | USNM AC8CC31 | 840 | *Antennarius multiocellatus* | No | |
| *Antennarius multiocellatus* | UW 117826 | | 888 | *Antennarius multiocellatus* | Yes | |
| *Antennarius pardalis* | USNM 405173 | USNM AD9NF63 | 892 | ambiguous | Yes | Specimen ID corroborated by tree topology |
| *Antennarius pauciradiatus* | MZUPRRP-I-00353 | UPR FL0161 | 927 | ambiguous | Yes | Unique species (no conspecific sequence to compare with) |
| *Antennarius pictus* | UW 118985 | | 703 | *Antennarius pictus* | Yes | |
| ***Antennarius randalli*** | **YPM 25207** | | **777** | ***Antennarius nummifer*** | **No** | **Likely to be mis-identified as *A. randalli* based on BOLD results and tree topology** |
| *Antennarius striatus* | ZSCM 32591 | in silico (BioProject number PRJEB12469) | 916 | no CO1 sequence | Yes | Specimen ID corroborated by tree topology |
| *Antennarius striatus* | USNM 405214 | USNM AD9NG45 | 825 | *Antennarius striatus* | No | |
| *Antennarius striatus* | UW 118819 | | 911 | *Antennarius striatus* | No | |
| *Antennatus coccineus* | USNM 439570 | USNM AG5NS91 | 825 | *Abantennarius coccineus, Abantennarius nummifer* | No | Specimen ID corroborated by tree topology |

| | | | | | | |
|---|---|---|---|---|---|---|
| *Antennatus coccineus* | UW 117829 | | 839 | *Abantennarius coccineus, Abantennarius nummifer* | Yes | Specimen ID corroborated by tree topology |
| *Antennatus dorehensis* | LSU 17134 | LSU 6171 | 888 | ambiguous | Yes | Specimen ID corroborated by tree topology |
| *Antennatus nummifer* | USNM 436321 | USNM AC1VH22 | 822 | *Abantennarius nummifer* | No | None of the *A. nummifer* sequences form a clade. The source of error is unclear. |
| *Antennatus nummifer* | UW 048070 | | 853 | *Antennarius nummifer* | Yes | None of the *A. nummifer* sequences form a clade. The source of error is unclear. Kept this sequence because of the three, its position is most consistent with past studies[18] |
| *Antennatus sanguineus* | CPUM-12938 | UPR FL1487 | 869 | *Antennatus sanguineus* | Yes | |
| *Antennatus sanguineus* | UW 118813 | | 833 | *Antennatus sanguineus* | No | |
| *Antennatus tuberosus* | USNM 392318 | USNM AG5NQ14 | 826 | *Antennatus tuberosus* | Yes | |
| *Antennatus tuberosus* | UW 118814 | | 759 | *Antennatus tuberosus* | No | |
| *Fowlerichthys avalonis* | USNM 422343 | USNM AG7PD25 | 919 | *Fowlerichthys avalonis* | Yes | |
| *Fowlerichthys ocellatus* | UW 150909 | | 868 | ambiguous | Yes | Specimen ID corroborated by tree topology |
| *Fowlerichthys radiosus* | KU:I 30134 | KU:IT 5131 | 791 | *Fowlerichthys radiosus* | No | This sequence was flagged for contamination during the BLC step |
| *Fowlerichthys scriptissimus* | UW 112642 | | 965 | *Fowlerichthys scriptissimus* | Yes | |
| *Histrio histrio* | USNM 413519 | USNM AC1VX42 | 849 | *Histrio histrio* | Yes | |
| *Histrio histrio* | UW 048052 | | 845 | *Histrio histrio* | No | |
| *Nudiantennarius subteres* | UW 117643 | | 846 | *Nudiantennarius subteres* | Yes | |
| *Brachionichthys australis* | | CSIRO GT 1304 | 806 | *Brachionichthys australis* | Yes | |

| | | | | | | |
|---|---|---|---|---|---|---|
| *Brachionichthys hirsutus* | | CSIRO BrahirOBay | 785 | *Brachionichthys hirsutus* | Yes | |
| *Brachiopsilus dianthus* | CSIRO 4995-01 | CSIRO GT 1707 | 71 | no CO1 sequence | Yes | No conspecific sequence to compare; placement in tree is consistent with expectations |
| *Thymichthys verrucosus* | CSIRO 4453-04 | CSIRO GT 1705 | 843 | *Thymichthys verrucosus* | Yes | |
| *Histiophryne cryptacanthus* | UW 117816 | | 946 | *Histiophryne cryptacanthus* | Yes | |
| *Histiophryne pogonia* | UW 119920 | | 693 | ambiguous | Yes | Sequences of *H. pogonius* are not monophyletic, a result also reported by Hart et al. [9] Taxonomic revision may be needed |
| *Lophiocharon lithinostomus* | UW 115749 | | 917 | *Lophiocharon trisignatus, Histiophryne psychedelica, Lophiocharon lithinostomus* | Yes | The three UCE-mined sequences of *Lophiocharon* are forming a clade separate from newly generated sequences. Likely an issue with missing data, not mis-identification |
| *Lophiocharon trisignatus* | UW 115748 | | 531 | *Lophiocharon trisignatus, Lophiocharon lithinostomus, Histiophryne psychedelica* | Yes | |
| *Kuiterichthys furcipilis* | CSIRO H 7233-01 | CSIRO GT 6329 | 409 | ambiguous | Yes | Rare species; no conspecific sequence to compare it to. Placement is generally consistent with expectations based on morphology[9] |
| *Phyllophryne scortea* | CSIRO H 7131-01 | CSIRO GT 5711 | 887 | *Phyllophryne scortea* | Yes | |

| | | | | | | |
|---|---|---|---|---|---|---|
| *Porophryne erythrodactylus* | AM I.44699 | | 898 | *Porophryne erythrodactylus* | Yes | |
| *Rhycherus filamentosus* | AM I.41560 | | 965 | *Rhycherus filamentosus* | Yes | |
| *Tathicarpus butleri* | | CSIRO GT 4294 | 336 | *Tathicarpus butleri* | No | |
| *Tathicarpus butleri* | LSU 13650 | LSU 371 | 895 | *Tathicarpus butleri* | Yes | |
| *Tetrabrachium ocellatum* | | QM GT 9504 | 358 | ambiguous | Yes | Specimen ID corroborated by tree topology |
| *Caulophryne pelagica* | USNM 422598 | USNM AG7PG44 | 973 | *Caulophryne pelagica* | Yes | |
| *Centrophryne spinulosa* | | gift from M. Miya | 957 | *Centrophryne spinulosa* | Yes | |
| *Ceratias holboelli* | UW 046158 | | 912 | *Ceratias uranoscopus, Ceratias tentaculatus, Ceratias holboelli* | Yes | No conspecific sequence to compare it to. Placement in tree is consistent with expectations |
| *Ceratias tentaculatus* | CSIRO H 6805-01 | CSIRO GT 2338 | 901 | *Ceratias sp., Ceratias tentaculatus, Ceratias holboelli* | Yes | No conspecific sequence to compare it to. Placement in tree is consistent with expectations |
| *Cryptopsaras couesii* | USNM 405020 | USNM AD9NC57 | 836 | *Cryptopsaras couesii* | Yes | |
| *Cryptopsaras couesii* | UW 048058 | | 772 | *Cryptopsaras couesii* | No | |
| *Bufoceratias thele* | | gift from M. Miya | 799 | *Bufoceratias thele* | Yes | |
| *Gigantactis elsmani* | AM I.28742 | | 835 | ambiguous | Yes | Branch lengths are short within *Gigantactis,* and specimens are rare. Difficult to identify species based on mitochondrial DNA |
| *Gigantactis gargantua* | not catalogued | G101 (gift from Tracey Sutton) | 408 | *Gigantactis vanhoeffeni* | Yes | Branch lengths are short within *Gigantactis,* and specimens are rare. Difficult to identify species based on mitochondrial DNA |
| *Gigantactis ios* | MCZ 163303 | KU:IT 5926 | 898 | *Gigantactis vanhoeffeni* | Yes | Branch lengths are short within |

| | | | | | | |
|---|---|---|---|---|---|---|
| | | | | | | *Gigantactis,* and specimens are rare. Difficult to identify species based on mitochondrial DNA |
| *Gigantactis microdontis* | not yet catalogued | G008 (gift from Tracey Sutton) | 123 | no CO1 sequence | Yes | Branch lengths are short within *Gigantactis,* and specimens are rare. Difficult to identify species based on mitochondrial DNA |
| *Gigantactis paxtoni* | CSIRO H 6177-01 | NMMB Taiwan GT 2 | 491 | *Gigantactis paxtoni, Gigantactis vanhoeffeni* | No | Branch lengths are short within *Gigantactis,* and specimens are rare. Difficult to identify species based on mitochondrial DNA |
| *Gigantactis paxtoni* | AM I.47816 | | 912 | *Gigantactis paxtoni, Gigantactis vanhoeffeni* | Yes | Branch lengths are short within *Gigantactis,* and specimens are rare. Difficult to identify species based on mitochondrial DNA |
| *Gigantactis vanhoeffeni* | UW 022178 | | 698 | *Gigantactis sp., Gigantactis vanhoeffeni* | Yes | Branch lengths are short within *Gigantactis,* and specimens are rare. Difficult to identify species based on mitochondrial DNA |
| *Himantolophus albinares* | MCZ 161521 | KU:IT 5264 | 947 | *Himantolophus albinares, Himantolophus groenlandicus, Himantolophus sagamius, Himantolophus appelii, Himantolophus sp., Himantolophus litoceras, Himantolophus stewarti* | Yes | Branch lengths are short within *Himantolophus*, difficult to distinguish species based on |

| | | | | | | |
|---|---|---|---|---|---|---|
| | | | | | | mitochrondrial DNA |
| *Himantolophus albinares* | YPM 27858 | | 941 | *Himantolophus albinares, Himantolophus groenlandicus, Himantolophus sagamius, Himantolophus appelii, Himantolophus sp., Himantolophus litoceras, Himantolophus stewarti* | No | Branch lengths are short within *Himantolophus*, difficult to distinguish species based on mitochrondrial DNA |
| *Himantolophus appelii* | UW 022179 | | 863 | *Himantolophus albinares, Himantolophus groenlandicus, Himantolophus sagamius, Himantolophus appelii, Himantolophus sp., Himantolophus litoceras, Himantolophus stewarti* | Yes | Branch lengths are short within *Himantolophus*, difficult to distinguish species based on mitochrondrial DNA |
| *Himantolophus appelii* | UW 025871 | | 765 | *Himantolophus albinares, Himantolophus groenlandicus, Himantolophus sagamius, Himantolophus appelii, Himantolophus sp., Himantolophus litoceras, Himantolophus stewarti* | No | Branch lengths are short within *Himantolophus*, difficult to distinguish species based on mitochrondrial DNA |
| *Himantolophus brevirostris* | MCZ 164737 | | 903 | *Himantolophus albinares, Himantolophus groenlandicus, Himantolophus sagamius, Himantolophus appelii, Himantolophus sp., Himantolophus litoceras, Himantolophus stewarti* | Yes | Branch lengths are short within *Himantolophus*, difficult to distinguish species based on mitochrondrial DNA |
| *Himantolophus groenlandicus* | not yet catalogued | G162 (gift from Tracey Sutton) | 561 | *Himantolophus albinares, Himantolophus groenlandicus, Himantolophus sagamius, Himantolophus appelii, Himantolophus sp.,* | Yes | Branch lengths are short within *Himantolophus*, difficult to distinguish species based on |

| | | | | | | |
|---|---|---|---|---|---|---|
| | | | | *Himantolophus litoceras, Himantolophus stewarti* | | mitochrondrial DNA |
| *Himantolophus sagamius* | SIO 02-2 | | 884 | *Himantolophus albinares, Himantolophus groenlandicus, Himantolophus sagamius, Himantolophus appelii, Himantolophus sp., Himantolophus litoceras, Himantolophus stewarti* | Yes | Branch lengths are short within *Himantolophus*, difficult to distinguish species based on mitochrondrial DNA |
| *Himantolophus stewarti* | CSIRO H 3278-01 | CSIRO Hst-4 | 671 | *Himantolophus albinares, Himantolophus groenlandicus, Himantolophus sagamius, Himantolophus appelii, Himantolophus sp., Himantolophus litoceras, Himantolophus stewarti* | Yes | Branch lengths are short within *Himantolophus*, difficult to distinguish species based on mitochrondrial DNA |
| *Haplophryne mollis* | | CSIRO GT 5120 | 18 | *Haplophryne mollis* | No | |
| *Linophryne arborifera* | MCZ 164736 | | 891 | *Haplophryne mollis* | Yes | Although specimen appears misidentified based on BOLD and tree topology, the voucher ID was confirmed by an expert (T. Pietsch). *Linophryne* and *Haplophryne* do not look alike, and *Haplophryne* is visually distinct from all other ceratioids. There may be a biological reason for the placement of this sequence |
| *Linophryne bicornis* | MCZ 138063 | | 882 | *Linophryne bicornis, Acentrophryne dolichonema* | Yes | Specimens are rare. It may be difficult to |

| | | | | | | |
|---|---|---|---|---|---|---|
| | | | | | | confirm species IDs based on molecular databases |
| *Linophryne brevibarbata* | not yet catalogued | G184 (gift from Tracey Sutton) | 362 | ambiguous | Yes | Specimens are rare. It may be difficult to confirm species IDs based on molecular databases |
| *Linophryne macrodon* | MCZ 164217 | | 887 | *Linophryne bicornis, Acentrophryne dolichonema* | Yes | Specimens are rare. It may be difficult to confirm species IDs based on molecular databases |
| *Photocorynus spiniceps* | not yet catalogued | DPND 3632 (gift from Tracey Sutton) | 884 | ambiguous | Yes | Specimens are rare. It may be difficult to confirm species IDs based on molecular databases |
| *Melanocetus johnsonii* | USNM 421406 | USNM AG7PG57 | 281 | *Melanocetus johnsonii* | Yes | |
| *Bertella idiomorpha* | UW 042301 | | 580 | *Bertella idiomorpha* | Yes | *Bertella* is nested within *Dolopichthys;* this has been found by other studies, and doesn't seem related to mis-identification |
| *Chaenophryne draco* | CSIRO H 3279-01 | CSIRO Cdr-1 | 662 | *Chaenophryne draco, Chaenophryne melanorhabdus* | Yes | |
| *Chaenophryne longiceps* | not yet catalogued | G135 (gift from Tracey Sutton) | 326 | *Chaenophryne longiceps* | No | |
| *Chaenophryne longiceps* | UW 115125 | | 682 | no CO1 sequence | Yes | Specimen ID corroborated by tree topology |
| *Chaenophryne melanorhabdus* | UW 049299 | | 850 | *Chaenophryne draco, Chaenophryne melanorhabdus* | Yes | |
| *Dolopichthys danae* | MCZ 164089 | | 680 | *Dolopichthys sp., Dolopichthys karsteni* | Yes | |
| *Dolopichthys karsteni* | MCZ 165969 | KU:IT 8149 | 694 | *Dolopichthys sp., Dolopichthys karsteni* | Yes | |

| | | | | | | |
|---|---|---|---|---|---|---|
| *Dolopichthys longicornis* | UW 046115 | | 717 | *Dolopichthys pullatus* | Yes | Specimens are rare. It may be difficult to confirm species IDs based on molecular databases |
| *Lasiognathus beebei* | MCZ 167891 | | 733 | ambiguous | Yes | Specimen ID corroborated by tree topology |
| *Lasiognathus sp* | not yet catalogued | G092 (gift from Tracey Sutton) | 271 | no CO1 sequence | No | Specimen ID corroborated by tree topology |
| *Lophodolos acanthognathus* | UW 029639 | | 836 | *Lophodolos acanthognathus* | Yes | |
| *Microlophichthys microlophus* | MCZ 164214 | | 853 | *Microlophichthys microlophus* | Yes | |
| *Oneirodes acanthias* | SIO 11-231 | | 884 | *Oneirodes sp., Oneirodes eschrichtii, Oneirodes notius, Oneirodes sabex, Oneirodes kreffti, Dolopichthys sp.* | Yes | Branch lengths are short in *Oneirodes,* difficult to identify species using mitochondrial DNA |
| *Oneirodes bulbosus* | UW 048049 | | 826 | *Oneirodes sp., Oneirodes eschrichtii, Oneirodes notius, Oneirodes sabex, Oneirodes kreffti, Dolopichthys sp.* | Yes | Branch lengths are short in *Oneirodes,* difficult to identify species using mitochondrial DNA |
| *Oneirodes epithales* | MCZ 164733 | | 919 | *Dolopichthys sp., Oneirodes sp., Oneirodes sabex* | Yes | Branch lengths are short in *Oneirodes,* difficult to identify species using mitochondrial DNA |
| *Oneirodes eschrichtii* | CSIRO 7320-03 | CSIRO GT 6725 | 826 | ambiguous | Yes | Branch lengths are short in *Oneirodes,* difficult to identify species using mitochondrial DNA |
| *Oneirodes kreffti* | CSIRO H 6876-06 | NMMB_Taiwan GT 5121 | 739 | *Oneirodes sp., Oneirodes eschrichtii, Oneirodes notius, Oneirodes kreffti, Oneirodes sabex, Dolopichthys sp.* | Yes | Branch lengths are short in *Oneirodes,* difficult to identify species using mitochondrial DNA |
| *Oneirodes macrosteus* | MCZ 161495 | KU:IT 5260 | 850 | *Oneirodes sp., Oneirodes macrosteus* | No | |

| | | | | | | |
|---|---|---|---|---|---|---|
| *Oneirodes macrosteus* | MCZ 164216 | | 913 | *Oneirodes sp., Oneirodes macrosteus* | Yes | |
| *Oneirodes theodoritissieri* | YPM 27790 | | 913 | ambiguous | Yes | Branch lengths are short in *Oneirodes,* difficult to identify species using mitochondrial DNA |
| *Oneirodes thompsoni* | UW 048054 | | 849 | *Oneirodes sp., Oneirodes eschrichtii, Oneirodes notius, Oneirodes sabex, Oneirodes kreffti, Dolopichthys sp.* | Yes | Branch lengths are short in *Oneirodes,* difficult to identify species using mitochondrial DNA |
| *Phyllorhinichthys balushkini* | MCZ 164228 | | 837 | *Microlophichthys microlophus* | Yes | There are no sequences for *Phyllorhinichthys* in BOLD database. Specimens are very rare. |
| *Puck pinnata* | SIO 04-35 | | 904 | *Puck pinnata* | Yes | |
| *Spiniphryne gladisfenae* | not yet catalogued | G051 (gift from Tracey Sutton) | 356 | no CO1 sequence | No | Sequences of *S. gladisfenae* are in different places in tree. We could not check this voucher specimen, so we chose the other sequence for which we had confirmation using BOLD |
| *Spiniphryne gladisfenae* | MCZ 164225 | | 892 | *Spiniphryne gladisfenae* | Yes | Sequences of *S. gladisfenae* are in different places in tree. We kept this sequence for which we had confirmation using BOLD |
| *Thaumatichthys pagidostomus* | | gift from M. Miya | 811 | *Thaumatichthys pagidostomus* | Yes | |
| *Chaunax abei* | SIO 19-9 | | 799 | *Chaunax abei, Chaunax breviradius, Chaunax cf. abei* | Yes | Branch lengths are short in *Chaunax,* difficult to identify |

| | | | | | | |
|---|---|---|---|---|---|---|
| | | | | | | species using mitochondrial DNA |
| *Chaunax endeavouri* | CSIRO H 7143-04 | NMMB Taiwan GT 5673 | 840 | *Chaunax endeavouri* | Yes | |
| *Chaunax nebulosus* | CSIRO H 6460-01 | NMMB_Taiwan/ CSIRO GT 875 | 904 | *Chaunax nebulosus* | Yes | |
| *Chaunax pictus* | | in silico (BioProject number PRJNA398732) | 711 | no CO1 sequence | No | Specimen ID corroborated by tree topology |
| *Chaunax pictus* | MCZ 166074 | KU:IT 8710 | 939 | *Chaunax fimbriatus, Chaunax penicillatus, Chaunax pictus* | Yes | Branch lengths are short in *Chaunax,* difficult to identify species using mitochondrial DNA |
| *Chaunax stigmaeus* | MCZ 166061 | KU:IT 8226 | 948 | *Chaunax fimbriatus, Chaunax penicillatus, Chaunax pictus* | Yes | Branch lengths are short in *Chaunax,* difficult to identify species using mitochondrial DNA |
| *Chaunax suttkusi* | USNM 400752 | USNM AA0AC48 | 952 | *Chaunax pictus* | No | Specimen ID is confirmed by tree topology. Branch lengths are short in *Chaunax,* difficult to identify species using mitochondrial DNA |
| *Chaunax suttkusi* | MCZ 166069 | KU:IT 8158 | 955 | *Chaunax pictus* | Yes | Specimen ID is confirmed by tree topology. Branch lengths are short in *Chaunax,* difficult to identify species using mitochondrial DNA |
| *Lophiodes caulinaris* | USNM 421345 | USNM AG7PC44 | 897 | *Lophiodes caulinaris* | Yes | |
| *Lophiodes caulinaris* | SIO 80-122 | | 643 | no CO1 sequence | No | Specimen ID corroborated by tree topology |

| | | | | | | |
|---|---|---|---|---|---|---|
| *Lophiodes mutilus* | CSIRO 6417-13 | CSIRO GT 943 | 925 | *Lophiodes cf. mutilus, Lophiodes sp.* B, *Lophiodes triradiatus* | Yes | Unique species, no conspecific sequence to compare with |
| *Lophiodes reticulatus* | FSBC 23940 | FWRI 0441 | 525 | *Lophiodes reticulatus* | No | |
| *Lophiodes reticulatus* | KU:I 30122 | KU:IT 5180 | 912 | *Lophiodes reticulatus* | Yes | |
| *Lophiodes spilurus* | USNM 421229 | USNM AG7PF91 | 840 | *Lophiodes spilurus* | No | |
| *Lophiodes spilurus* | SIO 10-114 | | 949 | no CO1 sequence | Yes | |
| *Lophiomus setigerus* | CSIRO H 6570-04 | CSIRO GT 1833 | 685 | ambiguous | No | Specimen ID corroborated by tree topology |
| *Lophiomus setigerus* | SIO 19-9 | | 923 | *Lophiomus setigerus, Lophius litulon* | Yes | Specimen ID corroborated by tree topology |
| *Lophius americanus* | UW 048064 | | 844 | *Lophius americanus* | Yes | |
| *Lophius gastrophysus* | MCZ 159682 | KU:IT 4960 | 929 | ambiguous | Yes | Specimen ID corroborated by tree topology |
| *Lophius litulon* | CSIRO H 7394066 | CSIRO GT 6889 | 839 | *Lophius litulon* | Yes | |
| *Sladenia sp* | | CSIRO Ssp-1 | 879 | no CO1 sequence | Yes | Specimen ID corroborated by tree topology |
| *Coelophrys micropa* | CSIRO H 6430-04 | NMMB_Taiwan/ CSIRO GT 892 | 787 | *Coelophrys micropa* | Yes | |
| *Dibranchus atlanticus* | USNM 405023 | USNM AD9NC63 | 894 | *Dibranchus atlanticus* | Yes | Sequences of *D. atlanticus* are not monophyletic despite species ID being confirmed by BOLD. For PCM tree, chose the sequence with the most data |
| *Dibranchus atlanticus* | UW 025869 | | 892 | *Dibranchus atlanticus* | No | |
| *Dibranchus spinosus* | USNM 422595 | USNM AG7PF46 | 924 | *Dibranchus sp.* E, *Dibranchus erinaceus, Dibranchus hystrix, Dibranchus spinosus* | Yes | |

| | | | | | | |
|---|---|---|---|---|---|---|
| *Dibranchus tremendus* | MCZ 164326 | KU:IT 7424 | 903 | *Dibranchus erinaceus, Dibranchus sp.* E, *Dibranchus hystrix* | Yes | Specimens are rare, it may be difficult to determine species ID using molecular databases. |
| *Halieutaea brevicauda* | CSIRO H 7283-02 | NMMB_Taiwan/ CSIRO GT 6514 | 729 | *Halieutaea fumosa, Halieutaea brevicauda* | Yes | |
| *Halieutaea brevicauda* | CSIRO 7266-08 | CSIRO GT 6523 | 696 | ambiguous | No | This sequence was flagged for contamination during the BLC step |
| ***Halieutaea fitzsimonsi*** | **SIO 19-9** | | **831** | ***Halieutaea sp.* VAD-2015, *Halieutaea stellata, Halieutaea stellata €*** | **No** | **Voucher specimen ID updated to *H. stellata* due to barcode results, tree topology, and further examination of morphology** |
| *Halieutaea fumosa* | CSIRO H 7394-27 | CSIRO GT 6865 | 821 | *Halieutaea fumosa, Halieutaea brevicauda* | No | |
| *Halieutaea fumosa* | LSU 13551 | LSU 267 | 826 | *Halieutaea fumosa, Halieutaea brevicauda* | Yes | |
| *Halieutaea stellata* | | NMMB_Taiwan/ CSIRO GT 5800 | 899 | *Halieutaea fitzsimonsi, Halieutaea coccinea, Halieutaea sp. 3, Halieutaea sp., Dibranchus sp., Halieutaea stellata* | Yes | |
| *Halieutaea stellata* | NSMT-P 75867 | KU:IT 10304 | 799 | *Halieutaea sp*. VAD-2015, *Halieutaea stellata, Halieutaea stellata (E)* | No | |
| *Halieutichthys aculeatus* | | USNM AB4OQ23 | 740 | *Halieutichthys aculeatus* | Yes | |
| *Halieutichthys aculeatus* | UW 021629 | | 677 | *Halieutichthys aculeatus* | No | |
| *Halieutopsis sp* | CSIRO H 7143-06 | CSIRO GT 5809 | 546 | no CO1 sequence | Yes | Unique species; no conspecific sequence to compare with |
| *Malthopsis gigas* | | NMMB_Taiwan/ CSIRO GT 5801 | 921 | *Malthopsis lutea, Malthopsis tiarella, Malthopsis gigas* | Yes | |

| | | | | | | |
|---|---|---|---|---|---|---|
| *Malthopsis sp* | | NMMB_Taiwan/ CSIRO GT 5793 | 262 | no CO1 sequence | No | Genus ID corroborated by tree topology |
| ***Malthopsis sp*** **(*Malthopsis gnoma*)** | **USNM 413973** | | **969** | **no CO1 sequence** | **No** | **Unrelated to other *Malthopsis* sequences. The voucher was confirmed to be *Malthopsis* (M. Girard pers. comm). The reason for non-monophyly is unclear** |
| *Ogcocephalus corniger* | FLMNH 182051 | FLMNH 2011-0090 | 866 | *Ogcocephalus declivirostris, Ogcocephalus vespertilio* | Yes | Specimen ID corroborated by tree topology |
| *Ogcocephalus corniger* | LSU 15551 | LSU 4818 | 811 | *Ogcocephalus declivirostris* | No | Specimen ID corroborated by tree topology |
| *Ogcocephalus cubifrons* | FLMNH 176166 | FLMNH 2007-0208 | 839 | *Ogcocephalus declivirostris, Ogcocephalus nasutus, Ogcocephalus vespertilio* | Yes | Branch lengths very short in *Ogcocephalus,* may be difficult to distinguish using mtDNA |
| *Ogcocephalus darwini* | SIO 00-154 | | 917 | ambiguous | Yes | Placement in tree is consistent with past studies[16] |
| *Ogcocephalus declivirostris* | FSBC 24830 | FWRI 1016 | 817 | *Ogcocephalus declivirostris, Ogcocephalus vespertilio* | No | *O. declivirostris* is not monophyletic, species boundaries in *Ogcocephalus* may require revision |
| *Ogcocephalus declivirostris* | LSU 13593 | LSU 314 | 885 | *Ogcocephalus declivirostris, Ogcocephalus vespertilio* | Yes | *O. declivirostris* is not monophyletic, species boundaries in *Ogcocephalus* may require revision |
| *Ogcocephalus nasutus* | LSU 15548 | LSU 4816 | 858 | *Ogcocephalus declivirostris, Ogcocephalus vespertilio* | Yes | Branch lengths very short in |

| | | | | | | |
|---|---|---|---|---|---|---|
| | | | | | | *Ogcocephalus,* may be difficult to distinguish using mtDNA |
| *Ogcocephalus pantostictus* | FSBC 24831 | FWRI 1021 | 748 | *Ogcocephalus declivirostris, Ogcocephalus nasutus, Ogcocephalus vespertilio* | Yes | Branch lengths very short in *Ogcocephalus,* may be difficult to distinguish using mtDNA |
| *Ogcocephalus parvus* | KU:I 30208 | KU:IT 5124 | 863 | *Ogcocephalus declivirostris, Ogcocephalus vespertilio* | No | *O. parvus* is not monophyletic, species boundaries in *Ogcocephalus* may require revision |
| *Ogcocephalus parvus* | USNM 442129 | USNM AE8WT40 | 869 | ambiguous | Yes | *O. parvus* is not monophyletic, species boundaries in *Ogcocephalus* may require revision. For purpose of comparative methods, we chose the sequence with the most genes |
| *Ogcocephalus parvus* | FSBC 24308 | FWRI 0732 | 843 | ambiguous | No | *O. parvus* is not monophyletic, species boundaries in *Ogcocephalus* may require revision |
| *Ogcocephalus radiatus* | UW 118987 | | 762 | ambiguous | Yes | Branch lengths very short in *Ogcocephalus,* may be difficult to distinguish using mtDNA |
| *Zalieutes elater* | USNM 421219 | USNM AG7PD32 | 868 | *Zalieutes elater* | Yes | |
| *Zalieutes elater* | SIO 10-20 | | 805 | *Zalieutes elater* | No | |

| | | | | | | |
|---|---|---|---|---|---|---|
| *Zalieutes mcgintyi* | FSBC 24829 | FWRI 1013 | 855 | ambiguous | Yes | Specimen ID is corroborated by tree topology |
| *Acanthurus olivaceus* | USNM 391238 | USNM AG5NP42 | 914 | *Acanthurus olivaceus, Acanthurus reversus* | Yes | Outgroup |
| *Naso unicornis* | USNM 399505 | USNM AG9RR39 | 941 | *Naso unicornis* | No | Outgroup |
| *Aulostomus maculatus* | | in silico (BioProject number PRJNA398732) | 666 | no CO1 sequence | Yes | Outgroup |
| *Forcipiger longirostris* | USNM 424145 | USNM AD9NL01 | 985 | *Forcipiger longirostris* | No | Outgroup |
| *Macroramphosus scolopax* | USNM 405231 | USNM AD9NG79 | 1000 | *Macroramphosus scolopax, Macroramphosus gracilis* | Yes | Outgroup |
| *Myripristis berndti* | | in silico (BioProject number PRJNA398732) | 815 | no CO1 sequence | Yes | Outgroup |
| *Labrus bergylta* | | in silico (BioProject number PRJEB13687) | 1040 | No CO1 sequence | No | Outgroup |
| *Luvarus imperialis* | | CSIRO "#1" | 993 | *Luvarus imperialis* | Yes | Outgroup |
| *Brotula barbata* | | in silico (BioProject number PRJEB12469) | 1035 | no CO1 sequence | Yes | Outgroup |
| *Pomacanthus paru* | | in silico (BioProject number PRJNA398732) | 728 | No CO1 sequence | No | Outgroup |
| *Cookeolus japonicus* | USNM 433134 | USNM AB4OQ12 | 909 | *Cookeolus japonicus, Heteropriacanthus cruentatus* | No | Outgroup |
| *Priacanthus arenatus* | USNM 405144 | USNM AD9NF05 | 916 | *Priacanthus arenatus* | Yes | Outgroup |
| *Pristigenys niphonia* | USNM 424846 | USNM AH0SX25 | 920 | *Pristigenys niphonia* | Yes | Outgroup |
| *Scatophagus argus* | CSIRO H 4877-02 | CSIRO "#1" | 954 | *Scatophagus argus* | Yes | Outgroup |
| *Owstonia tosaensis* | | CSIRO GT 1041 | 890 | *Owstonia tosaensis* | No | Outgroup |
| *Scomberomorus regalis* | | in silico (BioProject number PRJNA398732) | 709 | no CO1 sequence | Yes | Outgroup |
| *Siganus vulpinus* | | CSIRO UG0556 | 895 | *Siganus magnificus, Siganus vulpinus, Siganus unimaculatus* | No | Outgroup |

| | | | | | | |
|---|---|---|---|---|---|---|
| *Calamus calamus* | USNM 404444 | USNM AG8QS41 | 931 | *Calamus calamus, Calamus nodosus* | No | Outgroup |
| *Cephalopholis argus* | USNM 423470 | USNM AD9NP73 | 933 | *Cephalopholis argus* | Yes | Outgroup |
| *Zanclus cornutus* | USNM 392415 | USNM AG5NQ44 | 1025 | *Zanclus cornutus* | Yes | Outgroup |
| *Aracana aurita* | CSIRO H 6812-01 | CSIRO GT 2350 | 956 | *Aracana aurita, Aracana ornata* | Yes | Outgroup |
| *Odonus niger* | USNM 409200 | USNM AG7PW08 | 901 | *Melichthys niger, Odonus niger* | Yes | Outgroup |
| *Balistes capriscus* | USNM 405175 | USNM AD9NF67 | 887 | *Balistes capriscus* | No | Outgroup |
| *Diodon hystrix* | STRI-X-325 | STRI BFT11552 | 907 | *Diodon eydouxii, Diodon hystrix* | Yes | Outgroup |
| *Chilomycterus schoepfii* | USNM 415605 | USNM AC2WQ63 | 860 | *Chilomycterus schoepfii* | No | Outgroup |
| *Cyclichthys orbicularis* | | CSIRO GT 6432 | 779 | *Cyclichthys orbicularis* | No | Outgroup |
| *Mola mola* | | in silico (BioProject number PRJNA305960) | 1033 | no CO1 sequence | Yes | Outgroup |
| *Aluterus monoceros* | USNM 405891 | USNM AG9RG78 | 768 | *Aluterus monoceros* | Yes | Outgroup |
| *Oxymonacanthus longirostris* | AMS I.44739-002 | CSIRO UG0755 | 69 | *Oxymonacanthus longirostris* | No | Outgroup |
| *Lactoria cornuta* | USNM 403207 | USNM AG9RD87 | 941 | *Lactoria cornuta* | Yes | Outgroup |
| *Ostracion cubicus* | USNM 391997 | USNM AG5NP65 | 936 | *Ostracion cubicus, Ostracion immaculatus* | No | Outgroup |
| *Arothron hispidus* | USNM 400510 | USNM AG9RU11 | 868 | *Arothron hispidus, Arothron sp.* BDU-DMSRRK01 | Yes | Outgroup |
| *Takifugu rubripes* | | in silico (BioProject number PRJNA1434) | 1032 | No CO1 sequence | No | Outgroup |
| *Sphoeroides dorsalis* | USNM 433079 | USNM AB4OP01 | 825 | *Sphoeroides dorsalis, Sphoeroides parvus* | No | Outgroup |
| *Canthigaster rostrata* | MZUPRRP-I-00570 | UPR FL0412 | 810 | *Canthigaster rostrata, Canthigaster figueiredoi* | No | Outgroup |
| *Hollardia meadi* | USNM 431710 | USNM AG9RL06 | 1000 | ambiguous | No | Outgroup |
| *Paratriacanthodes retrospinis* | NMV A 29672-007 | CSIRO GT 1472 | 735 | *Bathyphylax bombifrons, Tydemania navigatoris, Paratriacanthodes retrospinis* | No | Outgroup |
| *Tripodichthys blochii* | USNM 424823 | USNM AH0SW64 | 812 | ambiguous | No | Outgroup |

**Table S2.** List of individuals with published UCE data (from which exons were mined for this study), and results from quality control steps. Studies: (1) Hart et al. [9]; (2) Ghezelayagh et al. [3]. "Tree topology" refers to all-individuals tree (Appendix A1, Fig. A1), "PCM" trees refer to the final trees used for comparative methods (Fig. 1; Appendix A1, Fig. A2, A3). Sequences that are likely to be misidentified are in bold.

| Original sequence name | Study | Voucher number | Tissue number | Number of exons mined | BOLD top hit | Used in PCM trees? | Notes |
|---|---|---|---|---|---|---|---|
| **Antennarius_commerson_F14** | **1** | **UW 118986** | | **14** | ***Antennarius pictus*** | **No** | **Checked voucher specimen and confirmed the correct ID is *A. pictus*** |
| Antennarius_indicus_A8 | 1 | UW 118818 | | 27 | *Antennarius indicus* | No | |
| Antennarius_multiocellatus_Amulti2 | 1 | UW 117827 | | 7 | no CO1 sequence | No | Specimen ID corroborated by tree topology |
| Antennarius_pardalis_A72 | 1 | CAS 235484 | TI 2010-109 | 61 | ambiguous | No | Specimen ID corroborated by tree topology |
| Antennarius_striatus_A70 | 1 | CAS 234886 | TI2010-132 | 28 | *Antennarius striatus* | No | |
| Antennarius_striatus_A71 | 1 | CAS 234890 | TI 2012-131 | 45 | *Antennarius striatus* | No | |
| Antennarius_striatus_A75 | 1 | NMNZ P.057359 | TS2 | 31 | *Antennarius striatus* | No | |
| Antennarius_striatus_A74 | 1 | NMNZ P.044669 | TS3 | 20 | ambiguous | No | Specimen ID corroborated by tree topology |
| Antennarius_striatus_00_09_Anst | 1 | | CBM-ZF-10514 | 6 | no CO1 sequence | No | Specimen ID corroborated by tree topology |
| Antennarius_striatus_117694-2 | 1 | UW 117694 | UW 117694 #2 | 16 | *Antennarius striatus* | No | |
| Antennarius_striatus_A21 | 1 | UW 118815 | | 19 | *Antennarius striatus* | No | |
| Antennarius_striatus_117695-3 | 1 | UW 117695 | UW 117695 #3 | 27 | *Antennarius striatus* | No | |
| Antennarius_striatus_117696-3 | 1 | UW 117696 | UW 117696 #3 | 10 | *Antennarius striatus* | No | |
| Antennarius_striatus_117696-4 | 1 | UW 117696 | UW 117696 #4 | 34 | *Antennarius striatus* | No | |
| Antennarius_striatus_117695-2 | 1 | UW 117695 | UW 117695 #2 | 34 | *Antennarius striatus* | No | |

| | | | | | | | |
|---|---|---|---|---|---|---|---|
| Antennatus_coccineus_T7143 | 1 | | KU:IT 7143 | 30 | *Abantennarius coccineus, Abantennarius nummifer* | No | Specimen ID corroborated by tree topology |
| Antennatus_coccineus_L2 | 1 | | NSMT-P 68051 | 9 | no CO1 sequence | No | Could not validate specimen ID. Sequence is related to *A. sanguineus,* not other *A. coccineus* |
| Antennatus_dorehensis_A69 | 1 | UW 157021 | | 8 | no CO1 sequence | No | Specimen ID corroborated by tree topology |
| Antennatus_nummifer_A11 | 1 | RUSI 65251 | KU:IT 5049 | 12 | ambiguous | No | None of the *A. nummifer* sequences form a clade, the source of error is unclear |
| Antennatus_rosaceus_A15 | 1 | | QS I. 38177 | 14 | no CO1 sequence | Yes | No conspecific sequence to compare with, but position in tree is consistent with other studies |
| Antennatus_strigatus_A1 | 1 | | LH05-205 | 13 | ambiguous | Yes | No conspecific sequence to compare with, but position in tree is consistent with other studies |
| Antennatus_tuberosus_Elmo | 1 | UW 115750 | | 31 | *Antennatus tuberosus* | No | |
| Antennarius_ocellatus_Anoc3 | 1 | UW 150911 | | 33 | ambiguous | No | Specimen ID corroborated by tree topology |
| Fowlerichthys_ocellatus_A52 | 1 | UW 150910 | | 28 | ambiguous | No | Specimen ID corroborated by tree topology |
| Antennarius_ocellatus_Anoc4 | 1 | UW 150912 | | 34 | no CO1 sequence | No | Specimen ID corroborated by tree topology |
| Fowlerichthys_radiosus_T3548 | 1 | | KU:IT 3548 | 5 | *Fowlerichthys radiosus* | No | |

| | | | | | | | |
|---|---|---|---|---|---|---|---|
| Antennarius_radiosus_A63 | 1 | | MCZ 144916 | 27 | *Fowlerichthys radiosus* | Yes | |
| Histrio_histrio_T5232 | 1 | | KU:IT 5232 | 33 | *Histrio histrio* | No | |
| Histrio_histrio_A37 | 1 | KU:I 29308 | KU:IT 3016 | 32 | ambiguous | No | Specimen ID corroborated by tree topology |
| Nudiantennarius_subteres_Asp2 | 1 | UW 119524 | | 12 | no CO1 sequence | No | Specimen ID corroborated by tree topology |
| Brachionichthys_australis | 2 | | CSIRO H 4465-01 | 322 | *Brachionichthys australis* | No | |
| Histiophryne_bougainvilli_A59 | 1 | UW 118990 | UW 118990 #4 | 32 | *Histiophryne bougainvilli* | Yes | |
| Histiophryne_bougainvilli_A60 | 1 | UW 118990 | UW 118990 #5 | 13 | *Histiophryne bougainvilli* | No | |
| Histiophryne_bougainvilli_Hboug2 | 1 | UW 118990 | UW 118990 #2 | 26 | *Histiophryne bougainvilli* | No | |
| Histiophryne_cryptacanthus_A28 | 1 | UW 117821 | | 30 | *Histiophryne cryptacanthus* | No | |
| Histiophryne_cryptacanthus_H6 | 1 | not cataloged, aquarium trade | | 13 | *Histiophryne cryptacanthus* | No | |
| Histiophryne_cryptacanthus_A56 | 1 | UW 117820 | | 14 | *Histiophryne cryptacanthus* | No | |
| Histiophryne_cryptacanthus_H7 | 1 | UW 117819 | | 17 | ambiguous | No | Specimen ID corroborated by tree topology |
| Histiophryne_cryptacanthus_H8 | 1 | not cataloged, aquarium trade | | 23 | *Histiophryne cryptacanthus* | No | |
| Histiophryne_cryptacanthus_H12 | 1 | UW 118816 | | 13 | ambiguous | No | This sequence is causing non-monophyly of *H. cryptacanthus* in the tree. Could not validate specimen ID |
| Histiophryne_maggiewalker_I-38176 | 1 | | QS I. 38176 | 18 | *Histiophryne sp.* QM I.38176, *Histiophryne maggiewalker* | Yes | |
| Histiophryne_pogonius_A32 | 1 | UW 118820 | | 29 | ambiguous | No | Sequences of *H. pogonius* are not monophyletic, a |

| | | | | | | | |
|---|---|---|---|---|---|---|---|
| | | | | | | | result also reported by Hart et al. 2022. Taxonomic revision may be needed |
| Histiophryne_psychedelica_A31 | 1 | | NCIP 6377 | 26 | ambiguous | Yes | No conspecific sequence to compare it to. Position in tree is generally consistent with expectations |
| Lophiocharon_lithinostomus_HANS119 | 1 | not reported | gift from Hsuan-Ching Ho | 28 | *Lophiocharon trisignatus, Lophiocharon lithinostomus, Histiophryne psychedelica* | No | The three UCE-mined sequences of *Lophiocharon* are forming a clade separate from newly generated sequences. Likely an issue with missing data, not mis-identification |
| Lophiocharon_lithinostomus_HANS120 | 1 | not reported | gift from Hsuan-Ching Ho | 40 | *Lophiocharon trisignatus, Lophiocharon lithinostomus, Histiophryne psychedelica* | No | |
| Lophiocharon_lithinostomus_Lophio-Bryan | 1 | not cataloged, aquarium trade | | 36 | *Lophiocharon trisignatus, Lophiocharon lithinostomus, Histiophryne psychedelica* | No | |
| Echinophryne_crassisspina_A25 | 1 | not reported | Yorke Peninsula, South Australia | 19 | ambiguous | No | Specimen ID corroborated by tree topology |
| Echinophryne_crassisspina_F5 | 1 | not reported | Yorke Peninsula, South Australia | 22 | ambiguous | Yes | Specimen ID corroborated by tree topology |
| Echinophryne_crassisspina_SAMP11544 | 1 | | SAM P11544 | 14 | *Echinophryne crassispina* | No | |

| | | | | | | | |
|---|---|---|---|---|---|---|---|
| Echinophryne_crassisspina_F7 | 1 | not reported | Yorke Peninsula, South Australia | 15 | ambiguous | No | Specimen ID corroborated by tree topology |
| Echinophryne_crassisspina_F4 | 1 | not reported | Yorke Peninsula, South Australia | 12 | no CO1 sequence | No | Specimen ID corroborated by tree topology |
| Phyllophryne_scortea_A42 | 1 | | NMV A29226.005 | 10 | *Phyllophryne scortea* | No | All UCE-mined sequences of *P. scortea* form a clade in tree, separate from newly generated conspecific sequences. Likely an issue with missing data, not mis-identification |
| Phyllophryne_scortea_F8 | 1 | not reported | Yorke Peninsula, South Australia | 11 | no CO1 sequence | No | |
| Phyllophryne_scortea_A44 | 1 | | SAM F17721 | 20 | *Phyllophryne scortea* | No | |
| Phyllophryne_scortea_F6 | 1 | not reported | Yorke Peninsula, South Australia | 25 | no CO1 sequence | No | |
| Phyllophryne_scortea_A57 | 1 | | SAM 11720 | 15 | ambiguous | No | |
| Phyllophryne_scortea_SAMP11722 | 1 | | SAM P11722 | 12 | no CO1 sequence | No | |
| Phyllophryne_scortea_F2 | 1 | not reported | Yorke Peninsula, South Australia | 18 | ambiguous | No | |
| Phyllophryne_scortea_SAM86337 | 1 | | SAM 86337 | 17 | *Phyllophryne scortea* | No | |
| Phyllophryne_scortea_SAM69556 | 1 | | SAM 69556 | 22 | *Phyllophryne scortea* | No | |
| Porophryne_erythrodactylus_A68 | 1 | UW 118988 | | 19 | *Porophryne erythrodactylus* | No | |
| Porophryne_erythrodactylus_I-43749-001 | 1 | | AMS I.43749.001 | 13 | *Porophryne erythrodactylus* | No | |
| Rhycherus_filamentosus_A45 | 1 | | NMV A29238.11 | 16 | *Rhycherus filamentosus* | No | |
| Rhycherus_filamentosus_NMVA22333 | 1 | | NMV A22333 | 19 | *Rhycherus filamentosus* | No | |

| | | | | | | | |
|---|---|---|---|---|---|---|---|
| Rhycherus_filamentosus_NMV24754 | 1 | | NMV 24754 | 21 | no CO1 sequence | No | Specimen ID corroborated by tree topology |
| Rhycherus_filamentosus_A46 | 1 | not reported | South Australia | 8 | no CO1 sequence | No | Specimen ID corroborated by tree topology |
| Tathicarpus_butleri_A47 | 1 | | WAM 32903.001 | 51 | *Tathicarpus butleri* | No | |
| Tathicarpus_butleri_A48 | 1 | | QS I. 38191 | 17 | *Tathicarpus butleri* | No | |
| Tathicarpus_butleri_A49 | 1 | | QS I. 38227 | 12 | no CO1 sequence | No | Specimen ID corroborated by tree topology |
| Tetrabrachium_ocellatum | 2 | UW 049710 | UW A67/ UW 049710B | 129 | *Tetrabrachium ocellatum* | No | |
| Tetrabrachium_ocellatum_UWA50 | 2 | UW 049710 | UW A50/ UW 049710A | 221 | *Tetrabrachium ocellatum* | No | |
| Tetrabrachium_ocellatum_C | 1 | UW 049710 | UW 049710C | 14 | no CO1 sequence | No | Specimen ID corroborated by tree topology |
| Caulophryne_pelagica | 2 | | NMNZ P.043296/TS3 | 75 | *Caulophryne polynema, Caulophryne pelagica, Caulophryne jordani* | No | |
| **Caulophryne_pelagica_L3_S208** | **1** | | **NSMT-P 93887(1)** | **11** | ***Melanocetus murrayi*** | **No** | **Specimen likely to be misidentified as *Caulophryne* based on BOLD results and tree topology** |
| Cryptopsaras_couesii | 2 | | YPM ICH 25702 | 37 | *Cryptopsaras couesii* | No | |
| Diceratias_pileatus | 2 | not reported | UW D7/04-091 (gift from M. Miya) | 89 | *Diceratias pileatus* | Yes | |
| Diceratias_sp | 2 | not reported | UW D27 | 96 | no CO1 sequence | No | Specimen ID corroborated by tree topology |
| Gigantactis_vanhoeffeni | 2 | | YPM ICH 027791 | 93 | *Gigantactis vanhoeffeni, Gigantactis paxtoni* | No | |

| | | | | | | | |
|---|---|---|---|---|---|---|---|
| Rhynchactis_macrothrix_04_016_S219 | 1 | UW 048054 | | 12 | ambiguous | Yes | Very rare species. Position in tree is consistent with taxonomic hypotheses (within Gigantactinidae) |
| Haplophryne_mollis | 2 | | NMNZ P.041209/TS3 | 82 | *Haplophryne mollis* | Yes | |
| **Melanocetus_johnsonii** | **2** | **UW 115879** | | **109** | ***Melanocetus murrayi*** | **No** | **Specimen likely to be misidentified as *M. johnsonii,* based on BOLD results and tree topology** |
| **Melanocetus_murrayi** | **2** | | **YPM ICH 027583** | **94** | ***Melanocetus johnsonii*** | **No** | **Specimen likely to be misidentified as *M. murrayi* based on BOLD results and tree topology** |
| Neoceratias_spinifer | 2 | not reported | UW D17 (gift from M. Miya) | 69 | *Neoceratias spinifer* | Yes | |
| Bertella_idiomorpha_L5_S209 | 1 | | NSMT-P 99996(1) | 9 | *Bertella idiomorpha* | No | |
| Chaenophryne_longiceps | 2 | | YPM ICH 25647 | 51 | *Chaenophryne longiceps* | No | |
| Dolopichthys_karsteni | 2 | | KU:IT 5926, MCZ 165969 | 357 | *Dolopichthys sp., Dolopichthys karsteni* | No | |
| Lasiognathus_intermedius | 2 | | YPM ICH 25927 | 45 | ambiguous | Yes | Rare species. Specimen ID corroborated by tree topology |
| Thaumatichthys_sp_G228_S222 | 1 | | G228 (gift from Chris Kenaley) | 16 | no CO1 sequence | No | Specimen ID corroborated by tree topology |
| Chaunax_pictus_04-115-Chpi | 1 | | Chpi_04_115 (gift from M. Miya) | 43 | *Chaunax pictus* | No | |

| | | | | | | | |
|---|---|---|---|---|---|---|---|
| Chaunax_stigmaeus | 2 | MCZ 166061 | KU:IT 8225 | 12 | Several *Chaunax* sequences | No | *Chaunax* have short branch lengths, hard to distinguish species using mitochondrial DNA |
| Chaunax_suttkusi | 2 | MCZ 166070 | KU:IT 8159 | 82 | *Chaunax pictus* | No | *Chaunax* have short branch lengths, hard to distinguish species using mitochondrial DNA |
| Lophiodes_caulinaris_04-078-Loca | 1 | | Loca_04_078 (gift from M. Miya) | 24 | no CO1 sequence | No | Specimen ID corroborated by tree topology |
| Lophius_gastrophysus | 2 | | Pers. Coll.: J.Moore, JAM99-109 | 156 | *Lophius gastrophysus* | No | |

**Table S3.** List of legacy marker sequences retrieved from GenBank, and results from quality control steps. Studies: (1) Miya et al. [10]; (2) Lundsten et al. [74]; (3) Near et al. [42]; (4) Chen et al. [75]; (5) Matschiner et al. [76].

| **Species** | **Voucher number** | **Study** | **Used in PCM trees?** | **BOLD top hit** | **GenBank ID** | | | | | | |
|---|---|---|---|---|---|---|---|---|---|---|---|
| | | | | | **ND1** | **CO1** | **glyt** | **myh6** | **rag1** | **sreb2** | **tbr** |
| *Acentrophryne dolichonema* | HUMZ 189134 | 1 | Yes | ambiguous | AB282855.1 | AB282855 | | | | | |
| *Caulophryne jordani* | Not reported | 1 | Yes | *Caulophryne jordani* | AP004417.1 | AP004417 | | | | | |
| *Ceratias uranoscopus* | NSMT-P 99996 | 1 | Yes | *Ceratias uranoscopus* | AB282851.1 | AB282851 | | | | | |
| *Chaunacops coloratus* | CAS 216055 | 2 | Yes | *Chaunacops coloratus* | | JN235967.1 | | | | | JN235981.1 |
| *Chaunacops coloratus* | CAS 232088 | 2 | Yes | *Chaunacops coloratus* | | JN235966.1 | | | | | JN235980.1 |
| *Chaunax tosaensis* | Not reported | 1 | Yes | *Chaunax penicillatus* | AP004416.1 | AP004416 | | | | | |
| *Lophius americanus* | YFTC 13690 | 3 | No | No CO1 sequence | | | JX188679 | JX189629 | JX189781 | JX189940 | JX189166 |
| *Lophius budegassa* | Not reported | 4 | No | No CO1 sequence | | | | | EF095637.1 | | |
| *Lophius vaillanti* | Not reported | 5 | No | No CO1 sequence | | | | HM050058.1 | | | HM050240.1 |
| *Malthopsis jordani* | BSKU 94678 | 1 | Yes | *Malthopsis sp.* | AP005978.1 | AP005978 | | | | | |
| *Sladenia gardineri* | Not reported | 1 | No | *Sladenia sp.* | AB282827.1 | AB282827 | | | | | |

**Table S4.** Habitat states used in analyses. Binary codes refer to presence (1) or absence (0).

| Species | Family | Suborder | Benthic shelf | Benthic slope or deeper | Bathypelagic | Notes |
|---|---|---|---|---|---|---|
| *Antennarius commerson* | Antennariidae | Antennarioidei | 1 | 0 | 0 | |
| *Antennarius maculatus* | Antennariidae | Antennarioidei | 1 | 0 | 0 | |
| *Antennarius multiocellatus* | Antennariidae | Antennarioidei | 1 | 0 | 0 | |
| *Antennarius pardalis* | Antennariidae | Antennarioidei | 1 | 0 | 0 | |
| *Antennarius pictus* | Antennariidae | Antennarioidei | 1 | 0 | 0 | |
| *Antennarius hispidus* | Antennariidae | Antennarioidei | 1 | 0 | 0 | |
| *Antennarius striatus* | Antennariidae | Antennarioidei | 1 | 0 | 0 | |
| *Antennarius indicus* | Antennariidae | Antennarioidei | 1 | 0 | 0 | |
| *Antennatus rosaceus* | Antennariidae | Antennarioidei | 1 | 0 | 0 | |
| *Antennarius pauciradiatus* | Antennariidae | Antennarioidei | 1 | 0 | 0 | |
| *Antennatus sanguineus* | Antennariidae | Antennarioidei | 1 | 0 | 0 | |
| *Antennatus coccineus* | Antennariidae | Antennarioidei | 1 | 0 | 0 | |
| *Antennatus tuberosus* | Antennariidae | Antennarioidei | 1 | 0 | 0 | |
| *Antennatus strigatus* | Antennariidae | Antennarioidei | 1 | 0 | 0 | |
| *Antennatus nummifer* | Antennariidae | Antennarioidei | 1 | 0 | 0 | |
| *Antennatus dorehensis* | Antennariidae | Antennarioidei | 1 | 0 | 0 | |
| *Histrio histrio* | Antennariidae | Antennarioidei | 1 | 0 | 0 | While sometimes considered pelagic, we consider it benthic following [77] (it rests on floating sargassum weeds) |
| *Nudiantennarius subteres* | Antennariidae | Antennarioidei | 1 | 0 | 0 | |
| *Fowlerichthys radiosus* | Antennariidae | Antennarioidei | 1 | 0 | 0 | |
| *Fowlerichthys avalonis* | Antennariidae | Antennarioidei | 1 | 0 | 0 | |
| *Fowlerichthys ocellatus* | Antennariidae | Antennarioidei | 1 | 1 | 0 | |
| *Fowlerichthys scriptissimus* | Antennariidae | Antennarioidei | 1 | 0 | 0 | |
| *Histiophryne cryptacanthus* | Histiophrynidae | Antennarioidei | 1 | 0 | 0 | |
| *Histiophryne pogonia* | Histiophrynidae | Antennarioidei | 1 | 0 | 0 | |
| *Histiophryne psychedelica* | Histiophrynidae | Antennarioidei | 1 | 0 | 0 | |
| *Histiophryne bougainvilli* | Histiophrynidae | Antennarioidei | 1 | 0 | 0 | |
| *Histiophryne maggiewalker* | Histiophrynidae | Antennarioidei | 1 | 0 | 0 | |
| *Lophiocharon lithinostomus* | Histiophrynidae | Antennarioidei | 1 | 0 | 0 | |
| *Lophiocharon trisignatus* | Histiophrynidae | Antennarioidei | 1 | 0 | 0 | |
| *Tathicarpus butleri* | Tathicarpidae | Antennarioidei | 1 | 0 | 0 | |

| | | | | | |
|---|---|---|---|---|---|
| *Kuiterichthys furcipilis* | Rhycheridae | Antennarioidei | 1 | 0 | 0 |
| *Porophryne erythrodactylus* | Rhycheridae | Antennarioidei | 1 | 0 | 0 |
| *Phyllophryne scortea* | Rhycheridae | Antennarioidei | 1 | 0 | 0 |
| *Echinophryne crassisspina* | Rhycheridae | Antennarioidei | 1 | 0 | 0 |
| *Rhycherus filamentosus* | Rhycheridae | Antennarioidei | 1 | 0 | 0 |
| *Brachionichthys hirsutus* | Brachionichthyidae | Antennarioidei | 1 | 0 | 0 |
| *Brachionichthys australis* | Brachionichthyidae | Antennarioidei | 1 | 0 | 0 |
| *Brachiopsilus dianthus* | Brachionichthyidae | Antennarioidei | 1 | 0 | 0 |
| *Thymichthys verrucosus* | Brachionichthyidae | Antennarioidei | 1 | 0 | 0 |
| *Tetrabrachium ocellatum* | Tetrabrachiidae | Antennarioidei | 1 | 0 | 0 |
| *Malthopsis jordani* | Ogcocephalidae | Ogcocephaloidei | 0 | 1 | 0 |
| *Malthopsis gigas* | Ogcocephalidae | Ogcocephaloidei | 0 | 1 | 0 |
| *Dibranchus atlanticus* | Ogcocephalidae | Ogcocephaloidei | 0 | 1 | 0 |
| *Dibranchus spinosus* | Ogcocephalidae | Ogcocephaloidei | 0 | 1 | 0 |
| *Dibranchus tremendus* | Ogcocephalidae | Ogcocephaloidei | 0 | 1 | 0 |
| *Coelophrys micropa* | Ogcocephalidae | Ogcocephaloidei | 0 | 1 | 0 |
| *Halieutopsis sp* | Ogcocephalidae | Ogcocephaloidei | 1 | 1 | 0 |
| *Halieutichthys aculeatus* | Ogcocephalidae | Ogcocephaloidei | 1 | 1 | 0 |
| *Ogcocephalus corniger* | Ogcocephalidae | Ogcocephaloidei | 1 | 1 | 0 |
| *Ogcocephalus parvus* | Ogcocephalidae | Ogcocephaloidei | 1 | 0 | 0 |
| *Ogcocephalus cubifrons* | Ogcocephalidae | Ogcocephaloidei | 1 | 0 | 0 |
| *Ogcocephalus radiatus* | Ogcocephalidae | Ogcocephaloidei | 1 | 0 | 0 |
| *Ogcocephalus pantostictus* | Ogcocephalidae | Ogcocephaloidei | 1 | 0 | 0 |
| *Ogcocephalus declivirostris* | Ogcocephalidae | Ogcocephaloidei | 1 | 1 | 0 |
| *Ogcocephalus nasutus* | Ogcocephalidae | Ogcocephaloidei | 1 | 1 | 0 |
| *Ogcocephalus darwini* | Ogcocephalidae | Ogcocephaloidei | 1 | 0 | 0 |
| *Zalieutes elater* | Ogcocephalidae | Ogcocephaloidei | 1 | 0 | 0 |
| *Zalieutes mcgintyi* | Ogcocephalidae | Ogcocephaloidei | 1 | 1 | 0 |
| *Halieutaea brevicauda* | Ogcocephalidae | Ogcocephaloidei | 1 | 1 | 0 |
| *Halieutaea fumosa* | Ogcocephalidae | Ogcocephaloidei | 0 | 1 | 0 |
| *Halieutaea stellata* | Ogcocephalidae | Ogcocephaloidei | 1 | 1 | 0 |
| *Chaunacops coloratus* | Chaunacidae | Chaunacoidei | 0 | 1 | 0 |
| *Chaunax abei* | Chaunacidae | Chaunacoidei | 1 | 1 | 0 |
| *Chaunax nebulosus* | Chaunacidae | Chaunacoidei | 0 | 1 | 0 |
| *Chaunax pictus* | Chaunacidae | Chaunacoidei | 1 | 1 | 0 |
| *Chaunax stigmaeus* | Chaunacidae | Chaunacoidei | 0 | 1 | 0 |
| *Chaunax tosaensis* | Chaunacidae | Chaunacoidei | 0 | 1 | 0 |
| *Chaunax endeavouri* | Chaunacidae | Chaunacoidei | 1 | 1 | 0 |

| | | | | | | |
|---|---|---|---|---|---|---|
| *Chaunax suttkusi* | Chaunacidae | Chaunacoidei | 0 | 1 | 0 | |
| *Caulophryne pelagica* | Caulophrynidae | Ceratioidei | 0 | 0 | 1 | |
| *Caulophryne jordani* | Caulophrynidae | Ceratioidei | 0 | 0 | 1 | |
| *Centrophryne spinulosa* | Centrophrynidae | Ceratioidei | 0 | 0 | 1 | |
| *Ceratias holboelli* | Ceratiidae | Ceratioidei | 0 | 0 | 1 | |
| *Ceratias uranoscopus* | Ceratiidae | Ceratioidei | 0 | 0 | 1 | |
| *Ceratias tentaculatus* | Ceratiidae | Ceratioidei | 0 | 0 | 1 | |
| *Cryptopsaras couesii* | Ceratiidae | Ceratioidei | 0 | 0 | 1 | |
| *Gigantactis elsmani* | Gigantactinidae | Ceratioidei | 0 | 0 | 1 | potentially benthopelagic[78] |
| *Gigantactis gargantua* | Gigantactinidae | Ceratioidei | 0 | 0 | 1 | potentially benthopelagic[78] |
| *Gigantactis microdontis* | Gigantactinidae | Ceratioidei | 0 | 0 | 1 | potentially benthopelagic[78] |
| *Gigantactis ios* | Gigantactinidae | Ceratioidei | 0 | 0 | 1 | potentially benthopelagic[78] |
| *Gigantactis paxtoni* | Gigantactinidae | Ceratioidei | 0 | 0 | 1 | potentially benthopelagic[78] |
| *Gigantactis vanhoeffeni* | Gigantactinidae | Ceratioidei | 0 | 0 | 1 | potentially benthopelagic[78] |
| *Rhynchactis macrothrix* | Gigantactinidae | Ceratioidei | 0 | 0 | 1 | |
| *Diceratias pileatus* | Diceratiidae | Ceratioidei | 0 | 0 | 1 | potentially benthopelagic[78] |
| *Bufoceratias thele* | Diceratiidae | Ceratioidei | 0 | 0 | 1 | potentially benthopelagic[78] |
| *Himantolophus albinares* | Himantolophidae | Ceratioidei | 0 | 0 | 1 | |
| *Himantolophus appelii* | Himantolophidae | Ceratioidei | 0 | 0 | 1 | |
| *Himantolophus brevirostris* | Himantolophidae | Ceratioidei | 0 | 0 | 1 | |
| *Himantolophus sagamius* | Himantolophidae | Ceratioidei | 0 | 0 | 1 | |
| *Himantolophus groenlandicus* | Himantolophidae | Ceratioidei | 0 | 0 | 1 | |
| *Himantolophus stewarti* | Himantolophidae | Ceratioidei | 0 | 0 | 1 | |
| *Melanocetus johnsonii* | Melanocetidae | Ceratioidei | 0 | 0 | 1 | |
| *Lasiognathus intermedius* | Oneirodidae | Ceratioidei | 0 | 0 | 1 | potentially benthopelagic[78] |
| *Lasiognathus beebei* | Oneirodidae | Ceratioidei | 0 | 0 | 1 | potentially benthopelagic[78] |
| *Puck pinnata* | Oneirodidae | Ceratioidei | 0 | 0 | 1 | |
| *Bertella idiomorpha* | Oneirodidae | Ceratioidei | 0 | 0 | 1 | |
| *Dolopichthys longicornis* | Oneirodidae | Ceratioidei | 0 | 0 | 1 | |
| *Dolopichthys danae* | Oneirodidae | Ceratioidei | 0 | 0 | 1 | |
| *Dolopichthys karsteni* | Oneirodidae | Ceratioidei | 0 | 0 | 1 | |
| *Chaenophryne draco* | Oneirodidae | Ceratioidei | 0 | 0 | 1 | |
| *Chaenophryne melanorhabdus* | Oneirodidae | Ceratioidei | 0 | 0 | 1 | |
| *Chaenophryne longiceps* | Oneirodidae | Ceratioidei | 0 | 0 | 1 | |
| *Microlophichthys microlophus* | Oneirodidae | Ceratioidei | 0 | 0 | 1 | |
| *Phyllorhinichthys balushkini* | Oneirodidae | Ceratioidei | 0 | 0 | 1 | |
| *Lophodolos acanthognathus* | Oneirodidae | Ceratioidei | 0 | 0 | 1 | |

| | | | | | | |
|---|---|---|---|---|---|---|
| *Oneirodes acanthias* | Oneirodidae | Ceratioidei | 0 | 0 | 1 | |
| *Oneirodes kreffti* | Oneirodidae | Ceratioidei | 0 | 0 | 1 | |
| *Oneirodes eschrichtii* | Oneirodidae | Ceratioidei | 0 | 0 | 1 | |
| *Oneirodes bulbosus* | Oneirodidae | Ceratioidei | 0 | 0 | 1 | |
| *Oneirodes thompsoni* | Oneirodidae | Ceratioidei | 0 | 0 | 1 | |
| *Oneirodes epithales* | Oneirodidae | Ceratioidei | 0 | 0 | 1 | |
| *Oneirodes macrosteus* | Oneirodidae | Ceratioidei | 0 | 0 | 1 | |
| *Oneirodes theodoritissieri* | Oneirodidae | Ceratioidei | 0 | 0 | 1 | |
| *Spiniphryne gladisfenae* | Oneirodidae | Ceratioidei | 0 | 0 | 1 | |
| *Acentrophryne dolichonema* | Linophrynidae | Ceratioidei | 0 | 0 | 1 | |
| *Linophryne bicornis* | Linophrynidae | Ceratioidei | 0 | 0 | 1 | |
| *Linophryne macrodon* | Linophrynidae | Ceratioidei | 0 | 0 | 1 | |
| *Linophryne brevibarbata* | Linophrynidae | Ceratioidei | 0 | 0 | 1 | |
| *Haplophryne mollis* | Linophrynidae | Ceratioidei | 0 | 0 | 1 | |
| *Linophryne arborifera* | Linophrynidae | Ceratioidei | 0 | 0 | 1 | |
| *Photocorynus spiniceps* | Linophrynidae | Ceratioidei | 0 | 0 | 1 | |
| *Thaumatichthys pagidostomus* | Thaumatichthyidae | Ceratioidei | 0 | 0 | 1 | potentially benthopelagic[78] (T. Pietsch pers comm) |
| *Neoceratias spinifer* | Neoceratiidae | Ceratioidei | 0 | 0 | 1 | |
| *Lophiodes caulinaris* | Lophiidae | Lophioidei | 1 | 1 | 0 | |
| *Lophiodes spilurus* | Lophiidae | Lophioidei | 1 | 1 | 0 | |
| *Lophiodes reticulatus* | Lophiidae | Lophioidei | 1 | 1 | 0 | |
| *Lophiodes mutilus* | Lophiidae | Lophioidei | 1 | 1 | 0 | |
| *Lophiomus setigerus* | Lophiidae | Lophioidei | 1 | 1 | 0 | |
| *Lophius americanus* | Lophiidae | Lophioidei | 1 | 1 | 0 | |
| *Lophius gastrophysus* | Lophiidae | Lophioidei | 1 | 1 | 0 | |
| *Lophius litulon* | Lophiidae | Lophioidei | 1 | 1 | 0 | |
| *Sladenia sp* | Lophiidae | Lophioidei | 0 | 1 | 0 | |

**Table S5.** BioGeoBEARS model fitting results.

| **Master tree** | **Model** | **LnL** | **# param** | ***d*** | ***e*** | ***j*** | **AICw** |
|---|---|---|---|---|---|---|---|
| MCMCtree | DEC | -97.03 | 2 | 0.0051 | 1.00E-12 | NA | <0.01 |
| | DEC+J | -97.03 | 3 | 0.0051 | 1.00E-12 | 1.00E-05 | <0.01 |
| | DIVALIKE | -105.5 | 2 | 0.0055 | 1.00E-12 | NA | <0.01 |
| | DIVALIKE+J | -105.5 | 3 | 0.0055 | 1.00E-12 | 1.00E-05 | <0.01 |
| | BAYAREALIKE | -80.18 | 2 | 0.0018 | 0.0034 | NA | 0.24 |
| | BAYAREALIKE+J | -77.99 | 3 | 0.0015 | 0.003 | 0.0021 | **0.76** |
| RelTime | DEC | -93.64 | 2 | 0.0063 | 1.00E-12 | NA | <0.01 |
| | DEC+J | -93.65 | 3 | 0.0063 | 1.00E-12 | 1.00E-05 | <0.01 |
| | DIVALIKE | -101.6 | 2 | 0.0069 | 1.00E-12 | NA | <0.01 |
| | DIVALIKE+J | -101.6 | 3 | 0.0069 | 1.00E-12 | 1.00E-05 | <0.01 |
| | BAYAREALIKE | -80.05 | 2 | 0.0023 | 0.0041 | NA | 0.32 |
| | BAYAREALIKE+J | -78.25 | 3 | 0.0019 | 0.0036 | 0.0021 | **0.68** |

**Table S6.** MiSSE model fitting results. Blank cells indicate the model fit had a relative Akaike weight <0.05.

| **Tree estimation** | **With Plecto** | **Calibration method** | **Pruned** | **Akaike weight by number of rate classes** | | | | | | | | | |
|---|---|---|---|---|---|---|---|---|---|---|---|---|---|
| | | | | **1** | **2** | **3** | **4** | **5** | **6** | **7** | **8** | **9** | **10** |
| IQ-TREE | no | MCMCTree | no | 0.409 | 0.337 | 0.159 | 0.063 | - | - | - | - | - | - |
| IQ-TREE | no | MCMCTree | yes | 0.692 | 0.196 | 0.072 | - | - | - | - | - | - | - |
| IQ-TREE | yes | MCMCTree | no | 0.405 | 0.336 | 0.159 | 0.063 | - | - | - | - | - | - |
| IQ-TREE | yes | MCMCTree | yes | 0.691 | 0.197 | 0.072 | - | - | - | - | - | - | - |
| ASTRAL | no | MCMCTree | no | 0.077 | 0.349 | 0.289 | 0.159 | 0.074 | - | - | - | - | - |
| ASTRAL | no | MCMCTree | yes | 0.409 | 0.298 | 0.165 | 0.075 | - | - | - | - | - | - |
| ASTRAL | yes | MCMCTree | no | 0.077 | 0.348 | 0.289 | 0.159 | 0.074 | - | - | - | - | - |
| ASTRAL | yes | MCMCTree | yes | 0.409 | 0.298 | 0.165 | 0.075 | - | - | - | - | - | - |
| IQ-TREE | no | RelTime | no | - | 0.170 | 0.348 | 0.265 | 0.133 | 0.054 | - | - | - | - |
| IQ-TREE | no | RelTime | yes | - | 0.349 | 0.331 | 0.181 | 0.079 | - | - | - | - | - |
| IQ-TREE | yes | RelTime | no | - | 0.279 | 0.355 | 0.212 | 0.095 | - | - | - | - | - |
| IQ-TREE | yes | RelTime | yes | 0.052 | 0.423 | 0.296 | 0.140 | 0.056 | - | - | - | - | - |
| ASTRAL | no | RelTime | no | - | - | 0.159 | 0.280 | 0.250 | 0.156 | 0.080 | - | - | - |
| ASTRAL | no | RelTime | yes | - | 0.102 | 0.258 | 0.269 | 0.186 | 0.103 | - | - | - | - |
| ASTRAL | yes | RelTime | no | - | - | 0.208 | 0.297 | 0.229 | 0.129 | 0.061 | - | - | - |
| ASTRAL | yes | RelTime | yes | - | 0.137 | 0.285 | 0.262 | 0.166 | 0.086 | - | - | - | - |

**Table S7.** Specimen vouchers used for CT scanning, with MorphoSource archival numbers or Virtual Natural History Museum stack numbers ("VNHM"). * = published by Heiple et al. [25]. NS = new scan (published by this study, to be archived upon acceptance).

| Species | Family | Suborder | Specimen voucher | Voxel size (µm) | Reference number | Notes |
|---|---|---|---|---|---|---|
| *Antennarius commerson* | Antennariidae | Antennarioidei | UW 020983 | 34.1 | 000550132/ VNHM #10803 | |
| *Antennarius hispidus* | Antennariidae | Antennarioidei | UW 020872 | 35.5 | 000097935/ VNHM #112762 | |
| *Antennarius indicus* | Antennariidae | Antennarioidei | UW 118817 | 35.5 | 000097953/ VNHM #112763 | |
| *Antennarius maculatus* | Antennariidae | Antennarioidei | UW 20828 | 29.8 | 000504368* | |
| *Antennarius multiocellatus* | Antennariidae | Antennarioidei | ANSP 121882 | 30 | 000550138/ VNHM #6186 | |
| *Antennarius pictus* | Antennariidae | Antennarioidei | UW 020852 | 35 | 000068482/ VNHM #105733 | |
| *Antennarius striatus* | Antennariidae | Antennarioidei | UW 21490 | 35.5 | 000097941/ VNHM #112760 | |
| *Antennatus coccineus* | Antennariidae | Antennarioidei | UW 20863 | 29.8 | 000504364* | |
| *Antennatus dorehensis* | Antennariidae | Antennarioidei | UW 118989 | 18.4 | 000097955/ VNHM #112766 | |
| *Antennatus nummifer* | Antennariidae | Antennarioidei | UW 20991 | 23 | 000097940/ VNHM #112770 | |
| *Antennatus pauciradiatus* | Antennariidae | Antennarioidei | MCZ 36235 | 9.9 | **NS** | |
| *Antennatus rosaceus* | Antennariidae | Antennarioidei | UW 20880 | 13.1 | 000097937/ VNHM #112758 | |
| *Antennatus sanguineus* | Antennariidae | Antennarioidei | UW 22624 | 24.8 | 000097943/ VNHM #112774 | |
| *Antennatus strigatus* | Antennariidae | Antennarioidei | SIO 11-86 | 35.5 | 000504358* | |
| *Antennatus tuberosis* | Antennariidae | Antennarioidei | UW 48622 | 23 | 000097947/ VNHM 112769 | |
| *Fowlerichthys avalonis* | Antennariidae | Antennarioidei | UW 20813 | 35.5 | 000504383* | |
| *Fowlerichthys ocellatus* | Antennariidae | Antennarioidei | UW 48073 | 29.8 | 000097946/ VNHM #112752 | |
| *Fowlerichthys radiosus* | Antennariidae | Antennarioidei | UW 151706 | 24.8 | 000097958/ VNHM #112772 | |
| *Fowlerichthys scriptissimus* | Antennariidae | Antennarioidei | UW 20838 | 34 | 000097934/ VNHM #112754 | |
| *Histrio histrio* | Antennariidae | Antennarioidei | UW 020959 | 26.5 | 000068486 | |

| | | | | | | |
|---|---|---|---|---|---|---|
| *Nudiantennarius subteres* | Antennariidae | Antennarioidei | UW 119524 | 23 | 000097957/ VNHM #112771 | |
| *Brachionichthys australis* | Brachionichthyidae | Antennarioidei | UW 116843 | 24.8 | 000097950/ VNHM #112773 | |
| *Brachionichthys hirsutus* | Brachionichthyidae | Antennarioidei | UW 21018 | 34 | **NS** | |
| *Brachiopsilus dianthus* | Brachionichthyidae | Antennarioidei | CSIRO H 4995-01 | 35.1 | **NS** | |
| *Thymichthys verrucosus* | Brachionichthyidae | Antennarioidei | CSIRO H 4453-04 | 31.9 | **NS** | |
| *Histiophryne cryptacanthus* | Histiophrynidae | Antennarioidei | UW 118816 | 29.8 | **NS** | |
| *Histiophryne pogonius* | Histiophrynidae | Antennarioidei | UW 119522 | 35.5 | 000097956/ VNHM #112776 | |
| *Histiophryne psychedelica* | Histiophrynidae | Antennarioidei | UW 22454 | 35.5 | 000097942/ VNHM #112778 | |
| *Lophiocharon lithinostomus* | Histiophrynidae | Antennarioidei | UW 115749 | 29.8 | 000097949/ VNHM #112751 | |
| *Lophiocharon trisignatus* | Histiophrynidae | Antennarioidei | UW 43037 | 35.5 | **NS** | |
| *Echinophryne crassispina* | Rhycheridae | Antennarioidei | UW 20982 | 18.1 | **NS** | |
| *Kuiterichthys furcipilis* | Rhycheridae | Antennarioidei | UW 20985 | 34 | **NS** | |
| *Phyllophryne scortea* | Rhycheridae | Antennarioidei | AMS I.17614-028 | 31.9 | 000504352* | |
| *Porophryne erythrodactylus* | Rhycheridae | Antennarioidei | UW 118988 | 23.4 | 000550144/ VNHM #112775 | |
| *Rhycherus filamentosus* | Rhycheridae | Antennarioidei | AMS I.41560-001 | 28.4 | **NS** | |
| *Tathicarpus butleri* | Tathicarpidae | Antennarioidei | UW 21038 | 18.1 | 000504379* | |
| *Tetrabrachium ocellatum* | Tetrabrachiidae | Antennarioidei | UW 021021 | 15.6 | 000550150/ VNHM #108272 | |
| *Caulophryne jordani* | Caulophrynidae | Ceratioidei | UW 113763 | 28 | 000495725* | |
| *Centrophryne spinulosa* | Centrophrynidae | Ceratioidei | UW 117074 | 28 | 000499464* | |
| *Ceratias holboelli* | Ceratiidae | Ceratioidei | UW 157500 | 35.1 | 000504316* | |
| *Cryptopsaras couesi* | Ceratiidae | Ceratioidei | UW 157503 | 25.9 | **NS** | |
| *Bufoceratias thele* | Diceratiidae | Ceratioidei | UW 156168 | 25.9 | 000499516* | |
| *Diceratias bispinosus* | Diceratiidae | Ceratioidei | UW 156169 | 26.9 | 000495095* | Stand-in for *D. pileatus* in the tree |
| *Gigantactis paxtoni* | Gigantactinidae | Ceratioidei | AMS I.47816-007 | 28.4 | 000495731* | |
| *Gigantactis vanhoeffeni* | Gigantactinidae | Ceratioidei | UW 157508 | 29.8 | 000503541* | |
| *Himantolophus albinares* | Himantolophidae | Ceratioidei | MCZ 161522 | 25.9 | 000503535* | |
| *Himantolophus appelii* | Himantolophidae | Ceratioidei | UW 25871 | 29.8 | 000499552* | |
| *Himantolophus groenlandicus* | Himantolophidae | Ceratioidei | MCZ 49842 | 25.9 | 000501788* | |
| *Himantolophus sagamius* | Himantolophidae | Ceratioidei | LACM 42698-1 | 17 | 000494769* | |

| | | | | | | |
|---|---|---|---|---|---|---|
| *Himantolophus stewarti* | Himantolophidae | Ceratioidei | CSIRO H 2985-01 | 51.1 | **NS** | The only new specimen not scanned at FHL (due to size); scanned instead at Evans Lab at Rice University |
| *Haplophryne mollis* | Linophrynidae | Ceratioidei | UW 157518 | 24.5 | 000070425/ VNHM #106359 | |
| *Linophryne arborifera* | Linophrynidae | Ceratioidei | MCZ 49822 | 35.5 | 000499492* | |
| *Melanocetus johnsonii* | Melanocetiidae | Ceratioidei | UW 157527 | 25.9 | **NS** | |
| *Bertella idiomorpha* | Oneirodidae | Ceratioidei | UW 48712 | 25.9 | 000503529* | |
| *Chaenophryne melanorhabdus* | Oneirodidae | Ceratioidei | UW 46491 | 35.5 | 000503696* | |
| *Dolopichthys karsteni* | Oneirodidae | Ceratioidei | MCZ 149624 | 17.4 | 000499703* | |
| *Dolopichthys longicornis* | Oneirodidae | Ceratioidei | UW 46115 | 8.8 | 000504262* | |
| *Lasiognathus beebei* | Oneirodidae | Ceratioidei | MCZ 167891 | 26.9 | 000503709* | |
| *Lophodolos acanthognathus* | Oneirodidae | Ceratioidei | UW 157541 | 35.5 | **NS** | |
| *Microlophichthys microlophus* | Oneirodidae | Ceratioidei | UW 157545 | 28 | 000504280* | |
| *Oneirodes bulbosus* | Oneirodidae | Ceratioidei | UW 48101 | 35.5 | **NS** | |
| *Oneirodes eschrichtii* | Oneirodidae | Ceratioidei | CSIRO H 7320-03 | 35.5 | 000494939* | |
| *Oneirodes kreffti* | Oneirodidae | Ceratioidei | AMS I.22810-029 | 35.5 | **NS** | |
| *Oneirodes thompsoni* | Oneirodidae | Ceratioidei | UW 117115 | 28 | 000504274* | |
| *Phyllorhinichthys balushkini* | Oneirodidae | Ceratioidei | UW 20826 | 28 | 000504297* | |
| *Puck pinnata* | Oneirodidae | Ceratioidei | SIO 04-35 | 25.9 | **NS** | |
| *Spiniphryne gladisfenae* | Oneirodidae | Ceratioidei | UW 157553 | 26.9 | 000501782* | |
| *Thaumatichthys binghami* | Thaumatichthyidae | Ceratioidei | UW 47537 | 17 | 000499470* | Stand-in for *T. pagidostomus* in the tree |
| *Chaunacops melanostomus* | Chaunacidae | Chaunacoidei | AMS I.31151-004 | 28 | **NS** | Stand-in for *C. coloratus* in the tree |
| *Chaunax abei* | Chaunacidae | Chaunacoidei | SIO 19-9 | 28 | **NS** | |
| *Chaunax endeavouri* | Chaunacidae | Chaunacoidei | AMS I.29738-004 | 35.5 | **NS** | |
| *Chaunax pictus* | Chaunacidae | Chaunacoidei | UW 158220 | 25.9 | 000504340* | |
| *Chaunax stigmaeus* | Chaunacidae | Chaunacoidei | UW 21468 | 28 | **NS** | |
| *Chaunax suttkusi* | Chaunacidae | Chaunacoidei | UW 46521 | 28 | **NS** | |
| *Chaunax tosaensis* | Chaunacidae | Chaunacoidei | UW 113953 | 35.5 | **NS** | |
| *Lophiodes caulinaris* | Lophiidae | Lophioidei | UW 43047 | 25.9 | 000504346* | |
| *Lophiodes mutilus* | Lophiidae | Lophioidei | UF 159767 | 8.3 | 000051387 | |

| | | | | | | |
|---|---|---|---|---|---|---|
| *Lophiodes reticulatus* | Lophiidae | Lophioidei | FSBC 23940 | 35.5 | **NS** | |
| *Lophiodes spilurus* | Lophiidae | Lophioidei | CSULB 3583 | 35.7 | VNHM #107068 | |
| *Lophiomus setigerus* | Lophiidae | Lophioidei | UW 16758 | 35.1 | 000504334* | |
| *Lophius americanus* | Lophiidae | Lophioidei | VIMS 35563 | 25.9 | 000097821/ VNHM #9720 | |
| *Sladenia remiger* | Lophiidae | Lophioidei | AMS I.29742-007 | 25.9 | 000504328* | |
| *Coelophrys arca* | Ogcocephalidae | Ogcocephaloidei | SIO 70-340 | 16.6 | **NS** | Stand-in for *C. micropa* in the tree |
| *Dibranchus atlanticus* | Ogcocephalidae | Ogcocephaloidei | UW 21712 | 18.1 | **NS** | |
| *Dibranchus tremendus* | Ogcocephalidae | Ogcocephaloidei | UW 46510 | 35.5 | **NS** | |
| *Halieutaea brevicauda* | Ogcocephalidae | Ogcocephaloidei | CSIRO H 7266-08 | 35.5 | **NS** | |
| *Halieutaea stellata* | Ogcocephalidae | Ogcocephaloidei | SIO 19-9 | 29.1 | **NS** | Voucher specimen ID was changed from *H. fitzsimonsi* to *H. stellata* based on molecular data |
| *Halieutichthys aculeatus* | Ogcocephalidae | Ogcocephaloidei | UW 25505 | 18.1 | **NS** | |
| *Halieutopsis stellifera* | Ogcocephalidae | Ogcocephaloidei | SIO 69-19 | 16.6 | **NS** | |
| *Malthopsis gigas* | Ogcocephalidae | Ogcocephaloidei | CSIRO H 7148-01 | 35.5 | **NS** | |
| *Malthopsis jordani* | Ogcocephalidae | Ogcocephaloidei | MCZ 51677 | 9.9 | **NS** | |
| *Ogcocephalus corniger* | Ogcocephalidae | Ogcocephaloidei | UF 182051 | 35.5 | **NS** | |
| *Ogcocephalus cubifrons* | Ogcocephalidae | Ogcocephaloidei | UF 176166 | 35.5 | **NS** | |
| *Ogcocephalus darwini* | Ogcocephalidae | Ogcocephaloidei | SIO H51-51-65A | 71.4 | 000550156/ VNHM #5684 | |
| *Ogcocephalus declivirostris* | Ogcocephalidae | Ogcocephaloidei | FSBC 24830 | 28 | **NS** | |
| *Ogcocephalus nasutus* | Ogcocephalidae | Ogcocephaloidei | CSULB 1945 | 45 | VNHM #107014 | |
| *Ogcocephalus pantostictus* | Ogcocephalidae | Ogcocephaloidei | MCZ 11749 | 33 | **NS** | |
| *Ogcocephalus parvus* | Ogcocephalidae | Ogcocephaloidei | FSBC 24308 | 28 | **NS** | |
| *Zalieutes elater* | Ogcocephalidae | Ogcocephaloidei | UW 043027 | 35.5 | 000076080/ VNHM #108191 | |
| *Zalieutes mcgintyi* | Ogcocephalidae | Ogcocephaloidei | FSBC 24829 | 35.5 | **NS** | |

**Table S8.** Disparity analyses by suborder.

Table S8a. Procrustes variances and proportion of total lophiiform disparity for the three phenotypic datasets

| **Phenotype** | **Suborder** | **Procrustes variance** | **Proportion of total disparity** |
|---|---|---|---|
| body shape | Antennarioidei | 0.1825 | 0.129 |
| body shape | Ceratioidei | 0.5587 | 0.396 |
| body shape | Chaunacoidei | 0.0654 | 0.046 |
| body shape | Lophioidei | 0.1771 | 0.125 |
| body shape | Ogcocephaloidei | 0.4289 | 0.304 |
| skull shape | Antennarioidei | 0.000420 | 0.248 |
| skull shape | Ceratioidei | 0.000633 | 0.374 |
| skull shape | Chaunacoidei | 0.000102 | 0.060 |
| skull shape | Lophioidei | 0.000148 | 0.088 |
| skull shape | Ogcocephaloidei | 0.000390 | 0.230 |
| jaw shape | Antennarioidei | 0.000802 | 0.191 |
| jaw shape | Ceratioidei | 0.001735 | 0.414 |
| jaw shape | Chaunacoidei | 0.000253 | 0.060 |
| jaw shape | Lophioidei | 0.000408 | 0.097 |
| jaw shape | Ogcocephaloidei | 0.000990 | 0.236 |

Table S8b. Pairwise absolute differences by suborder, using body shape.

| | Antennarioidei | Ceratioidei | Chaunacoidei | Lophioidei | Ogcocephaloidei |
|---|---|---|---|---|---|
| Antennarioidei | NA | | | | |
| Ceratioidei | 0.376 | NA | | | |
| Chaunacoidei | 0.117 | 0.493 | NA | | |
| Lophioidei | 0.005 | 0.382 | 0.112 | NA | |
| Ogcocephaloidei | 0.246 | 0.130 | 0.364 | 0.252 | NA |

Table S8c. Pairwise absolute differences by suborder, using skull shape.

| | Antennarioidei | Ceratioidei | Chaunacoidei | Lophioidei | Ogcocephaloidei |
|---|---|---|---|---|---|
| Antennarioidei | NA | | | | |
| Ceratioidei | 0.000213 | NA | | | |
| Chaunacoidei | 0.000318 | 0.000530 | NA | | |
| Lophioidei | 0.000272 | 0.000484 | 0.000046 | NA | |
| Ogcocephaloidei | 0.000030 | 0.000243 | 0.000287 | 0.000241 | NA |

Table S8d. Pairwise absolute differences by suborder, using jaw shape.

| | Antennarioidei | Ceratioidei | Chaunacoidei | Lophioidei | Ogcocephaloidei |
|---|---|---|---|---|---|
| Antennarioidei | NA | | | | |
| Ceratioidei | 0.000933 | NA | | | |
| Chaunacoidei | 0.000549 | 0.001482 | NA | | |
| Lophioidei | 0.000394 | 0.001327 | 0.000155 | NA | |
| Ogcocephaloidei | 0.000188 | 0.000745 | 0.000737 | 0.000582 | NA |

Table S8e. Pairwise p-value by suborder, using body shape. Significant p-values in bold.

| | Antennarioidei | Ceratioidei | Chaunacoidei | Lophioidei | Ogcocephaloidei |
|---|---|---|---|---|---|
| Antennarioidei | NA | | | | |
| Ceratioidei | **0.001** | NA | | | |
| Chaunacoidei | 1.000 | 0.102 | NA | | |
| Lophioidei | 1.000 | 0.826 | **0.001** | NA | |
| Ogcocephaloidei | 0.295 | 0.982 | **0.001** | **0.023** | NA |

Table S8f. Pairwise p-value by suborder, using skull shape. Significant p-values in bold.

| | Antennarioidei | Ceratioidei | Chaunacoidei | Lophioidei | Ogcocephaloidei |
|---|---|---|---|---|---|
| Antennarioidei | NA | | | | |
| Ceratioidei | **0.046** | NA | | | |
| Chaunacoidei | 1.000 | 0.070 | NA | | |
| Lophioidei | 1.000 | **0.047** | 0.136 | NA | |
| Ogcocephaloidei | 1.000 | 0.347 | **0.007** | 0.088 | NA |

Table S8g. Pairwise p-value by suborder, using jaw shape. Significant p-values in bold.

| | Antennarioidei | Ceratioidei | Chaunacoidei | Lophioidei | Ogcocephaloidei |
|---|---|---|---|---|---|
| Antennarioidei | NA | | | | |
| Ceratioidei | **0.002** | NA | | | |
| Chaunacoidei | 1.000 | **0.003** | NA | | |
| Lophioidei | 1.000 | **0.021** | 0.141 | NA | |
| Ogcocephaloidei | 0.999 | 0.150 | **0.020** | 0.176 | NA |

**Table S9.** Disparity analyses by habitat.

Table S9a. Procrustes variances and proportion of total disparity for the three phenotypic datasets

| **Phenotype** | **Habitat** | **Procrustes variance** | **Proportion of total disparity** |
|---|---|---|---|
| body shape | shelf only | 0.3111 | 0.220 |
| body shape | slope only | 0.1638 | 0.116 |
| body shape | shelf and slope | 0.3790 | 0.268 |
| body shape | bathypelagic | 0.5587 | 0.396 |
| skull shape | shelf only | 0.000532 | 0.314 |
| skull shape | slope only | 0.000155 | 0.091 |
| skull shape | shelf and slope | 0.000374 | 0.221 |
| skull shape | bathypelagic | 0.000633 | 0.374 |
| jaw shape | shelf only | 0.001104 | 0.263 |
| jaw shape | slope only | 0.000251 | 0.060 |
| jaw shape | shelf and slope | 0.001099 | 0.262 |
| jaw shape | bathypelagic | 0.001735 | 0.414 |

Table S9b. Pairwise absolute differences by habitat, using body shape.

| | bathypelagic | shelf only | shelf and slope | slope only |
|---|---|---|---|---|
| bathypelagic | NA | | | |
| shelf only | 0.248 | NA | | |
| shelf and slope | 0.180 | 0.068 | NA | |
| slope only | 0.395 | 0.147 | 0.215 | NA |

Table S9c. Pairwise absolute differences by habitat, using skull shape.

| | bathypelagic | shelf only | shelf and slope | slope only |
|---|---|---|---|---|

| | | | | |
|---|---|---|---|---|
| bathypelagic | NA | | | |
| shelf only | 0.00010 | NA | | |
| shelf and slope | 0.00026 | 0.00016 | NA | |
| slope only | 0.00048 | 0.00038 | 0.00022 | NA |

Table S9d. Pairwise absolute differences by habitat, using jaw shape.

| | bathypelagic | shelf only | shelf and slope | slope only |
|---|---|---|---|---|
| bathypelagic | NA | | | |
| shelf only | 0.000632 | NA | | |
| shelf and slope | 0.000636 | 0.000005 | NA | |
| slope only | 0.001484 | 0.000853 | 0.000848 | NA |

Table S9e. Pairwise p-value by habitat, using body shape. Significant p-values in bold.

| | bathypelagic | shelf only | shelf and slope | slope only |
|---|---|---|---|---|
| bathypelagic | NA | | | |
| shelf only | **0.001** | NA | | |
| shelf and slope | 1 | 1 | NA | |
| slope only | 1 | 1 | **0.004** | NA |

Table S9f. Pairwise p-value by habitat, using skull shape. Significant p-values in bold.

| | bathypelagic | shelf only | shelf and slope | slope only |
|---|---|---|---|---|
| bathypelagic | NA | | | |
| shelf only | 0.829 | NA | | |
| shelf and slope | 0.239 | 1 | NA | |
| slope only | **0.008** | 0.997 | **0.026** | NA |

Table S9g. Pairwise p-value by habitat, using jaw shape. Significant p-values in bold.

| | bathypelagic | shelf only | shelf and slope | slope only |
|---|---|---|---|---|
| bathypelagic | NA | | | |
| shelf only | 0.173 | NA | | |
| shelf and slope | 0.309 | 1 | NA | |
| slope only | **0.001** | 0.996 | **0.001** | NA |

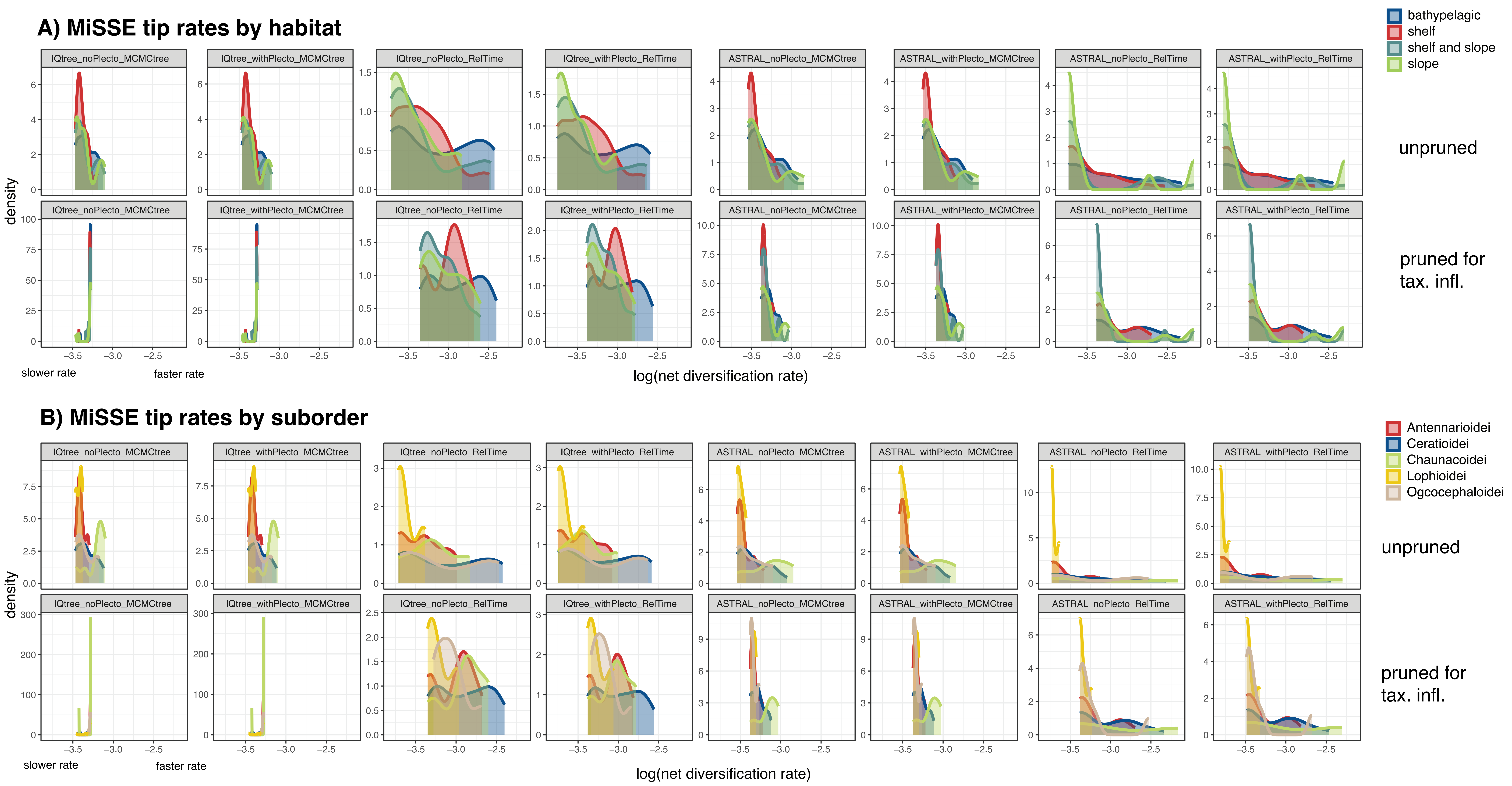

A) MiSSE tip rates by habitat
bathypelagic
shelf
shelf and slope
slope
IQtree_noPlecto_MCMCtree
IQtree_withPlecto_MCMCtree
IQtree_noPlecto_RelTime
IQtree_withPlecto_RelTime
ASTRAL_noPlecto_MCMCtree
ASTRAL_withPlecto_MCMCtree
ASTRAL_noPlecto_RelTime
ASTRAL_withPlecto_RelTime
unpruned
pruned for tax. infl.
density
slower rate
faster rate
log(net diversification rate)
B) MiSSE tip rates by suborder
Antennarioidei
Ceratioidei
Chaunacoidei
Lophioidei
Ogcocephaloidei
IQtree_noPlecto_MCMCtree
IQtree_withPlecto_MCMCtree
IQtree_noPlecto_RelTime
IQtree_withPlecto_RelTime
ASTRAL_noPlecto_MCMCtree
ASTRAL_withPlecto_MCMCtree
ASTRAL_noPlecto_RelTime
ASTRAL_withPlecto_RelTime
unpruned
pruned for tax. infl.
density
slower rate
faster rate
log(net diversification rate)

**Figure S1:** Model-averaged tip rates estimated by MiSSE by A) habitat and B) suborder. For model fits see Table S6. See justification in Appendix A2 for pruning to adjust for suspected taxonomic inflation.

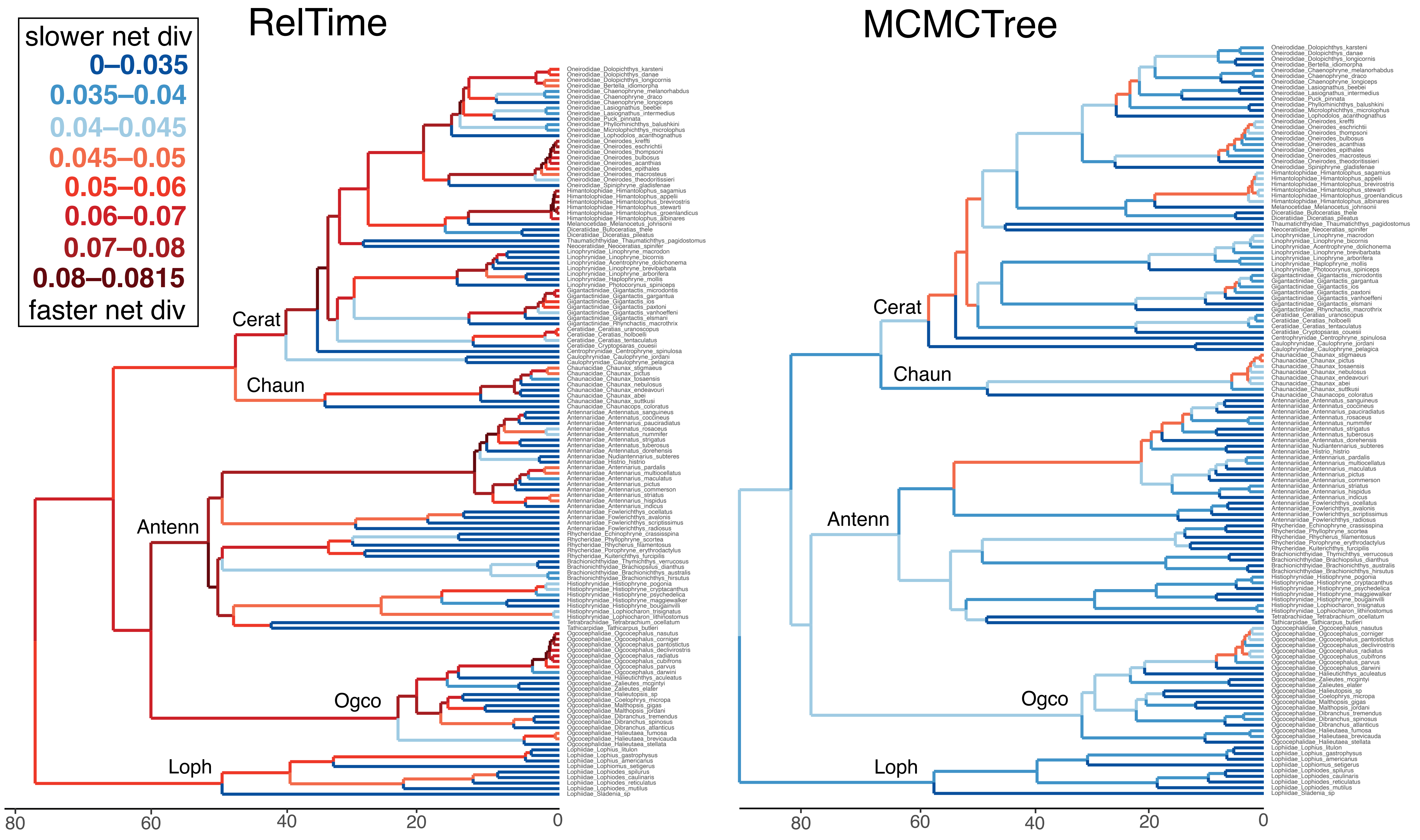

RelTime
MCMCTree
slower net div
0–0.035
0.035–0.04
0.04–0.045
0.045–0.05
0.05–0.06
0.06–0.07
0.07–0.08
0.08–0.0815
faster net div
Cerat
Chaun
Antenn
Ogco
Loph
80
60
40
20
0

**Figure S2:** Comparison of branch-associated diversification rates inferred by MiSSE on the master RelTime tree versus the master MCMCTree. Branches are colored according to a standard scale to compare the magnitude of rates between trees.

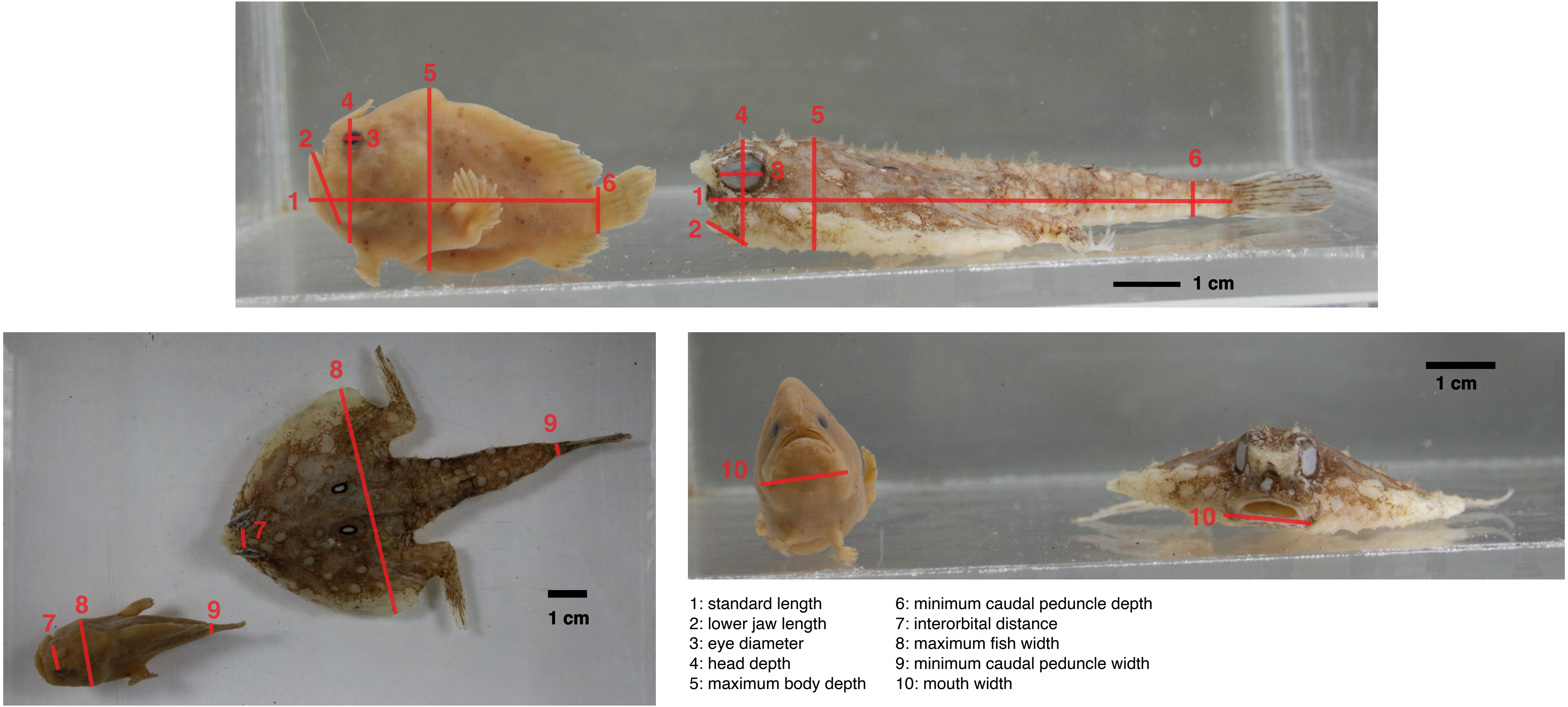

1: standard length
2: lower jaw length
3: eye diameter
4: head depth
5: maximum body depth
6: minimum caudal peduncle depth
7: interorbital distance
8: maximum fish width
9: minimum caudal peduncle width
10: mouth width

**Figure S3:** Illustration of ten body shape linear measurements on a laterally compressed frogfish and dorsoventrally compressed batfish.

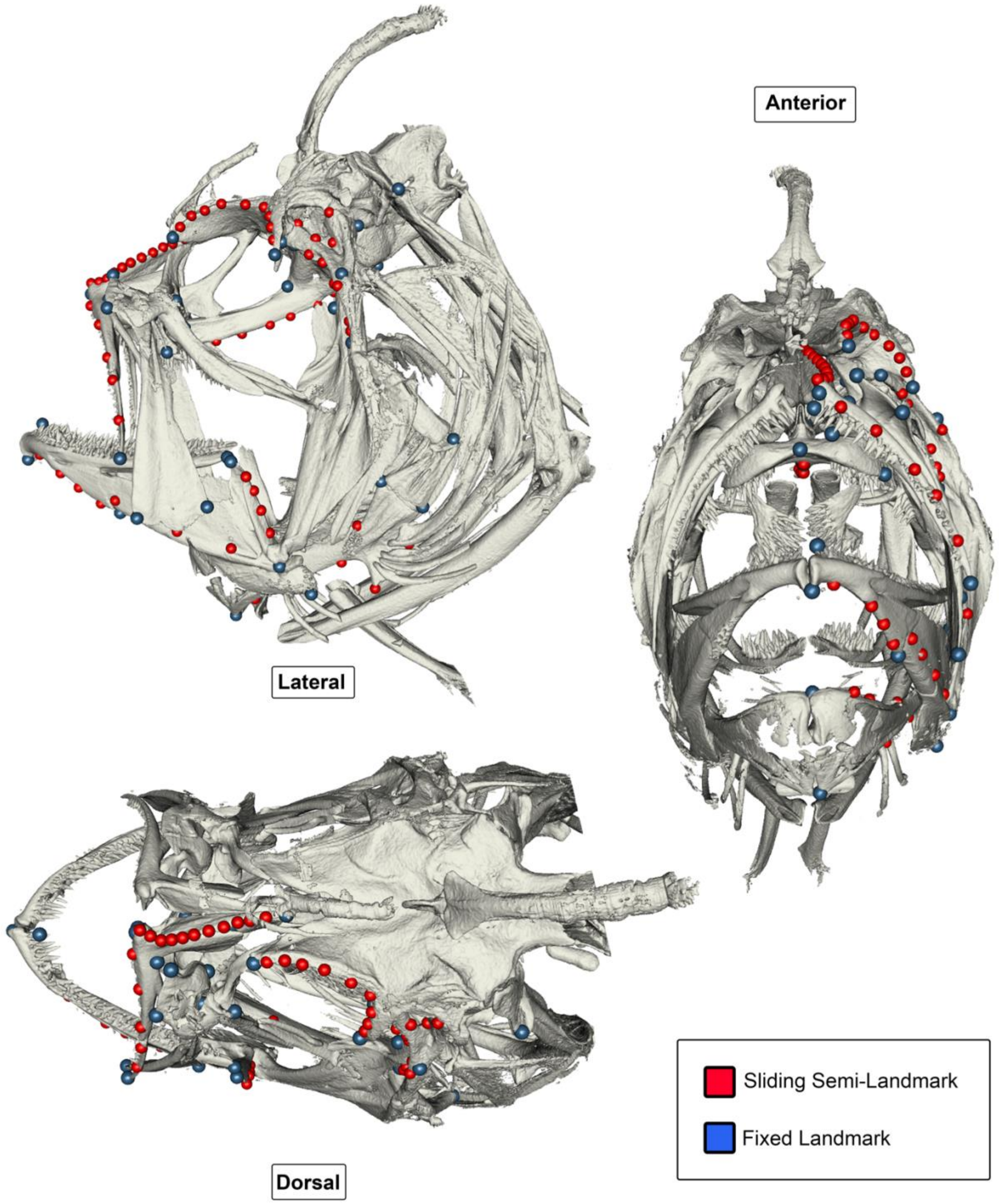

**Figure S4:** Illustration of landmarks placed on CT scans.

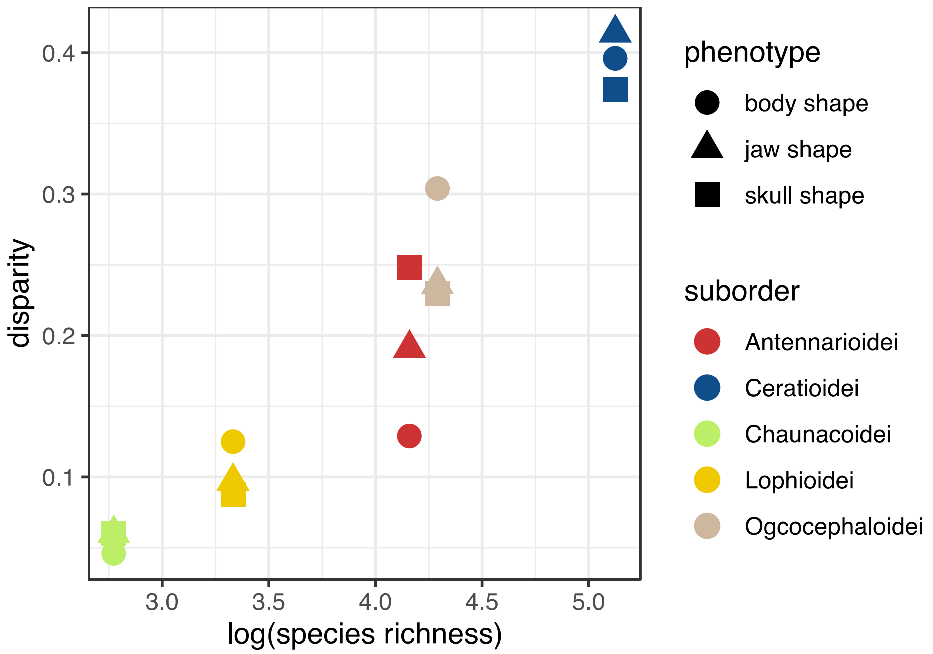

**Figure S5:** Illustration of disparity in body, skull and jaw shapes by species richness of the five suborders. See Tables S8 and S9 for full disparity analysis results.

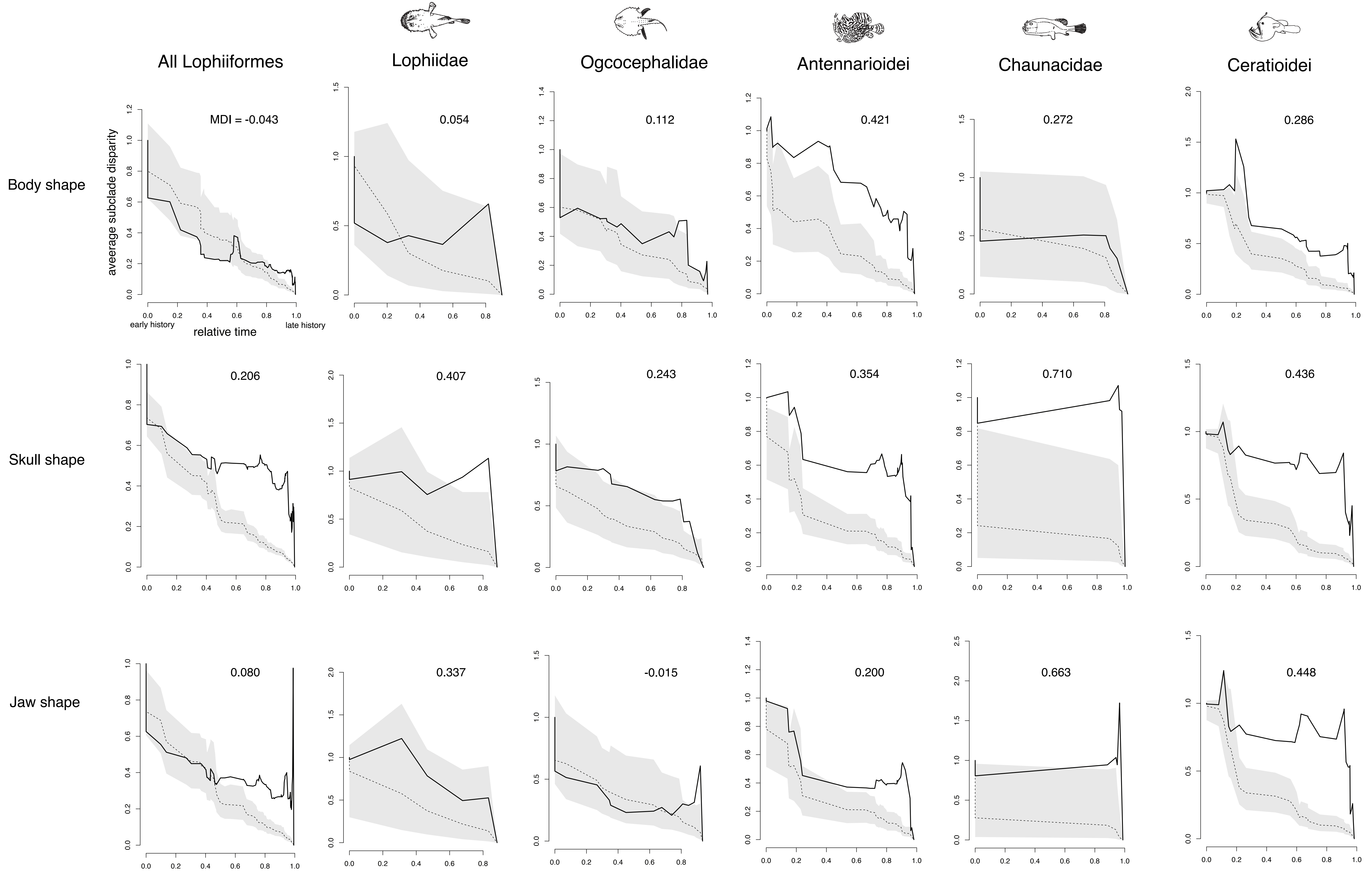
All Lophiiformes
Lophiidae
Ogcocephalidae
Antennarioidei
Chaunacidae
Ceratioidei
Body shape
Skull shape
Jaw shape
aveerage subclade disparity
relative time
early history
late history
MDI = -0.043
0.054
0.112
0.421
0.272
0.286
0.206
0.407
0.243
0.354
0.710
0.436
0.080
0.337
-0.015
0.200
0.663
0.448

**Figure S6:** Disparity of body, skull and jaw shapes through time for Lophiiformes and the five suborders individually. Negative values of the morphological disparity index indicate disparity is partitioned among subclades; positive values indicate disparity is partitioned within subclades. Solid black line shows the observed disparity; grey shading and dashed line shows expectations from a simulated Brownian motion null model inferred from the master MCMCTree.

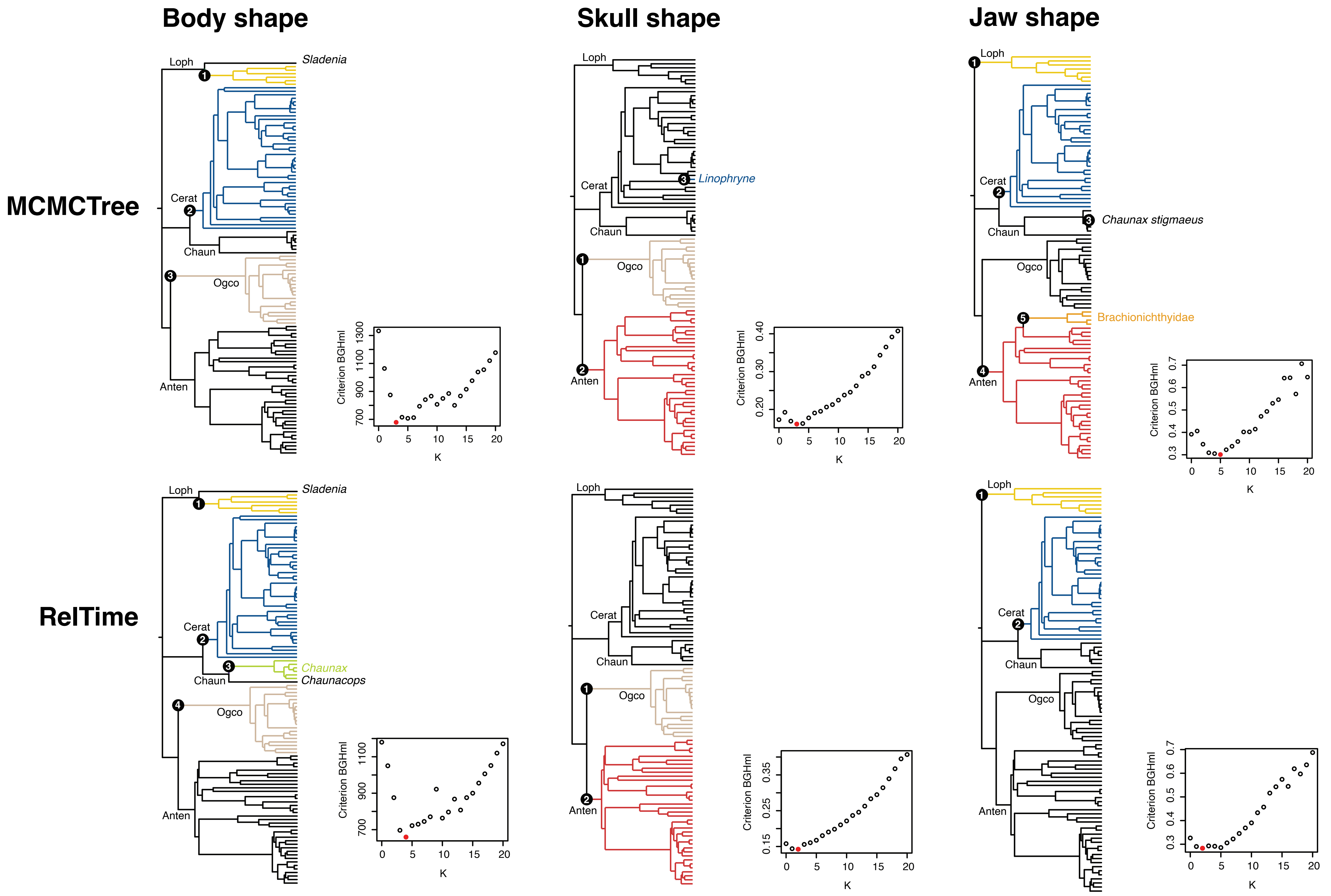
Body shape
Skull shape
Jaw shape
MCMCTree
RelTime
Loph
Cerat
Chaun
Ogco
Anten
Sladenia
Linophryne
Chaunax stigmaeus
Brachionichthyidae
Chaunax
Chaunacops
Criterion BGHml
K

**Figure S7:** Results of phyloEM model fitting using the two master trees. Inset panels show support for models with 0–20 shifts in adaptive regime, with the best-fitting model in red.

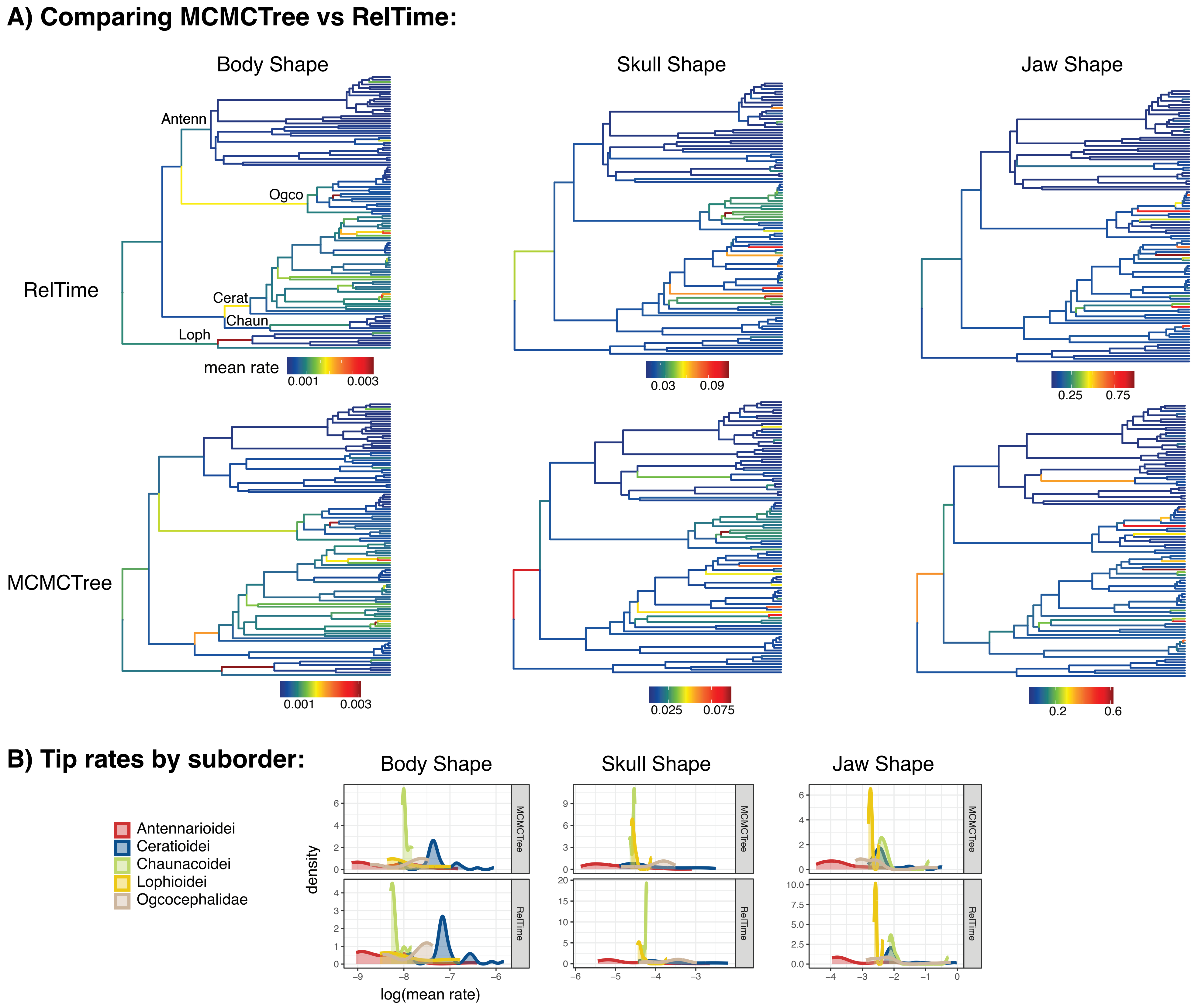

A) Comparing MCMCTree vs RelTime:
Body Shape
Skull Shape
Jaw Shape
RelTime
Antenn
Ogco
Cerat
Chaun
Loph
mean rate
0.001
0.003
0.03
0.09
0.25
0.75
MCMCTree
0.001
0.003
0.025
0.075
0.2
0.6
B) Tip rates by suborder:
Body Shape
Skull Shape
Jaw Shape
Antennarioidei
Ceratioidei
Chaunacoidei
Lophioidei
Ogcocephalidae
density
MCMCTree
RelTime
log(mean rate)

**Figure S8:** Comparison of branch-associated rates of body, skull and jaw shape evolution inferred by BayesTraits between the two master trees (A) and tip-associated evolutionary rates by suborder (B).

### SUPPLEMENTARY INFORMATION REFERENCES

9. Hart, P.B., Arnold, R.J., Alda, F., Kenaley, C.P., Pietsch, T.W., Hutchinson, D., and Chakrabarty, P. (2022). Evolutionary relationships of anglerfishes (Lophiiformes) reconstructed using ultraconserved elements. Mol. Phylogenet. Evol. *171*, 107459. 10.1016/j.ympev.2022.107459.

10. Miya, M., Pietsch, T.W., Orr, J.W., Arnold, R.J., Satoh, T.P., Shedlock, A.M., Ho, H.-C., Shimazaki, M., Yabe, M., and Nishida, M. (2010). Evolutionary history of anglerfishes (Teleostei: Lophiiformes): a mitogenomic perspective. BMC Evol. Biol. *10*, 58. 10.1186/1471-2148-10-58.

11. Arnold, R. (2014). Evolutionary Relationships of the Enigmatic Anglerfishes (Teleostei: Lophiiformes): Can Nuclear DNA Provide Resolution for Conflicting Morphological and Mitochondrial Phylogenies?

12. Rabosky, D.L., Chang, J., Title, P.O., Cowman, P.F., Sallan, L., Friedman, M., Kaschner, K., Garilao, C., Near, T.J., Coll, M., et al. (2018). An inverse latitudinal gradient in speciation rate for marine fishes. Nature *559*, 392–395. 10.1038/s41586-018-0273-1.

13. Pietsch, T.W., and Orr, J.W. (2007). Phylogenetic Relationships of Deep-sea Anglerfishes of the Suborder Ceratioidei (Teleostei: Lophiiformes) Based on Morphology. Copeia *2007*, 1–34. 10.1643/0045-8511(2007)7[1:PRODAO]2.0.CO;2.

14. Bertelsen, E. (1984). Ceratioidei: Development and relationships. In Ontogeny and Systematics of Fishes Special Publication Number 1., H. G. Moser, W. J. Richards, D. M. Cohen, M. P. Fahay, A. W. Kendall, and S. L. Richardson, eds. (American Society of Ichthyogists and Herpetologists), pp. 320–325.

15. Endo, H., and Shinohara, G. (1999). A new batfish,Coelophrys bradburyae (Lophiiformes: Ogcocephalidae) from Japan, with comments on the evolutionary relationships of the genus. Ichthyol. Res. *46*, 359–365. 10.1007/BF02673978.

16. Hunt, E. (2018). Inferring the phylogenetic relationships within two fish genera: insights into the biogeography of the northern Gulf of Mexico.

17. Arnold, R.J., and Pietsch, T.W. (2012). Evolutionary history of frogfishes (Teleostei: Lophiiformes: Antennariidae): A molecular approach. Mol. Phylogenet. Evol. *62*, 117–129. 10.1016/j.ympev.2011.09.012.

18. Pietsch, T.W., and Arnold, R.J. (2020). Frogfishes: Biodiversity, Zoogeography, and Behavioral Ecology (John Hopkins University Press).

19. Carnevale, G., and Pietsch, T.W. (2010). Eocene handfishes from Monte Bolca, with description of a new genus and species, and a phylogeny of the family Brachionichthyidae (Teleostei: Lophiiformes). Zool. J. Linn. Soc. *160*, 621–647. 10.1111/j.1096-3642.2009.00623.x.

20. Renema, W., Bellwood, D.R., Braga, J.C., Bromfield, K., Hall, R., Johnson, K.G., Lunt, P., Meyer, C.P., McMonagle, L.B., Morley, R.J., et al. (2008). Hopping Hotspots: Global Shifts in Marine Biodiversity. Science *321*, 654–657. 10.1126/science.1155674.

21. Stuart-Smith, J., Edgar, G.J., Last, P., Linardich, C., Lynch, T., Barrett, N., Bessell, T., Wong, L., and Stuart-Smith, R.D. (2020). Conservation challenges for the most threatened family of marine bony fishes (handfishes: Brachionichthyidae). Biol. Conserv. *252*, 108831. 10.1016/j.biocon.2020.108831.

22. Swann, J.B., Holland, S.J., Petersen, M., Pietsch, T.W., and Boehm, T. (2020). The immunogenetics of sexual parasitism. Science *369*, 1608–1615.

23. Bertelsen, E. (1951). The ceratioid fishes. Ontogeny, taxonomy, distribution and biology.

24. Pietsch, T.W., and Sutton, T.T. (2015). A New Species of the Ceratioid Anglerfish Genus *Lasiognathus* Regan (Lophiiformes: Oneirodidae) from the Northern Gulf of Mexico. Copeia *103*, 429–432. 10.1643/CI-14-181.

25. Heiple, Z., Huie, J.M., Medeiros, A.P.M., Hart, P.B., Goatley, C.H.R., Arcila, D., and Miller, E.C. (2023). Many ways to build an angler: diversity of feeding morphologies in a deep-sea evolutionary radiation. Biol. Lett. *19*, 20230049. 10.1098/rsbl.2023.0049.

26. Bertelsen, E., and Struhsaker, P.J. (1977). The ceratioid fishes of the genus Thaumatichthys: Osteology, relationships, distribution, and biology.

34. Friedman, M., Feilich, K.L., Beckett, H.T., Alfaro, M.E., Faircloth, B.C., Černý, D., Miya, M., Near, T.J., and Harrington, R.C. (2019). A phylogenomic framework for pelagiarian fishes (Acanthomorpha: Percomorpha) highlights mosaic radiation in the open ocean. Proc. R. Soc. B Biol. Sci. *286*, 20191502. 10.1098/rspb.2019.1502.

44. Orr, J.W. (1995). Phylogenetic relationships of the gasterosteiform fishes (Teleostei: Acanthomorpha).

45. Santaquiteria, A., Siqueira, A.C., Duarte-Ribeiro, E., Carnevale, G., White, W.T., Pogonoski, J.J., Baldwin, C.C., Ortí, G., Arcila, D., and Ricardo, B.-R. (2021).

Phylogenomics and Historical Biogeography of Seahorses, Dragonets, Goatfishes, and Allies (Teleostei: Syngnatharia): Assessing Factors Driving Uncertainty in Biogeographic Inferences. Syst. Biol. *70*, 1145–1162. 10.1093/sysbio/syab028.

46. Bannikov, A.F., and Tyler, J.C. (1995). Phylogenetic Revision of the Fish Families Luvaridae and †Kushlukiidae (Acanthuroidei), with a New Genus and Two New Species of Eocene Luvarids. Smithson. Contrib. Paleobiology *81*.

47. Gavrilov, Y., Kodina, L.A., Lubchenko, I.Y., and Muzylev, N.G. (1997). The Late Paleocene Anoxic Event in Epicontinental Seas of Peri-Tethys and Formation of the Sapropelite Unit: Sedimentology and Geochemistry. Lithol. Miner. Resour. *32*, 427–450.

48. Gavrilov, Y.O., Shcherbinina, E.A., and Oberhänsli, H. (2003). Paleocene-Eocene boundary events in the Northeastern Peri-Tethys. Geol. Soc. Am. Spec. Pap. *369*.

49. Kaya, M.Y., Dupont-Nivet, G., Frieling, J., Fioroni, C., Rohrmann, A., Altıner, S.Ö., Vardar, E., Tanyaş, H., Mamtimin, M., and Zhaojie, G. (2022). The Eurasian epicontinental sea was an important carbon sink during the Palaeocene-Eocene thermal maximum. Commun. Earth Environ. *3*, 124. 10.1038/s43247-022-00451-4.

50. Carnevale, G., Johnson, G.D., Marramà, G., and Bannikov, A.F. (2017). A reappraisal of the Eocene priacanthid fish *Pristigenys substriata* (Blainville, 1818) from Monte Bolca, Italy. J. Paleontol. *91*, 554–565. 10.1017/jpa.2017.19.

51. Friedman, M., and Carnevale, G. (2018). The Bolca Lagerstätten: shallow marine life in the Eocene. J. Geol. Soc. *175*, 569–579. 10.1144/jgs2017-164.

52. Arcila, D., and Tyler, J.C. (2017). Mass extinction in tetraodontiform fishes linked to the Palaeocene–Eocene thermal maximum. Proc. R. Soc. B Biol. Sci. *284*, 20171771. 10.1098/rspb.2017.1771.

53. Troyer, E.M., Betancur-R, R., Hughes, L.C., Westneat, M., Carnevale, G., White, W.T., Pogonoski, J.J., Tyler, J.C., Baldwin, C.C., Ortí, G., et al. (2022). The impact of paleoclimatic changes on body size evolution in marine fishes. Proc. Natl. Acad. Sci. *119*, e2122486119. 10.1073/pnas.2122486119.

54. Santini, F., and Tyler, J.C. (2003). A phylogeny of the families of fossil and extant tetraodontiform fishes (Acanthomorpha, Tetraodontiformes), Upper Cretaceous to Recent. Zool. J. Linn. Soc. *139*, 565–617. 10.1111/j.1096-3642.2003.00088.x.

55. Arcila, D., Alexander Pyron, R., Tyler, J.C., Ortí, G., and Betancur-R., R. (2015). An evaluation of fossil tip-dating versus node-age calibrations in tetraodontiform fishes (Teleostei: Percomorphaceae). Mol. Phylogenet. Evol. *82*, 131–145. 10.1016/j.ympev.2014.10.011.

56. Bannikov, A.F., Tyler, J.C., Arcila, D., and Carnevale, G. (2017). A new family of gymnodont fish (Tetraodontiformes) from the earliest Eocene of the Peri-Tethys

(Kabardino-Balkaria, northern Caucasus, Russia). J. Syst. Palaeontol. *15*, 129–146. 10.1080/14772019.2016.1149115.

57. Bannikov, A.F., and Tyler, J.C. (2008). A new genus and species of triggerfish from the Middle Eocene of the Northern Caucasus, the earliest member of the Balistidae (Tetraodontiformes). Paleontol. J. *42*, 615–620. 10.1134/S0031030108060075.

58. van der Boon, A., van der Ploeg, R., Cramwinckel, M.J., Kuiper, K.F., Popov, S.V., Tabachnikova, I.P., Palcu, D.V., and Krijgsman, W. (2019). Integrated stratigraphy of the Eocene-Oligocene deposits of the northern Caucasus (Belaya River, Russia): Intermittent oxygen-depleted episodes in the Peri-Tethys and Paratethys. Palaeogeogr. Palaeoclimatol. Palaeoecol. *536*, 109395. 10.1016/j.palaeo.2019.109395.

59. Pietsch, T.W., and Carnevale, G. (2011). A New Genus and Species of Anglerfish (Teleostei: Lophiiformes: Lophiidae) from the Eocene of Monte Bolca, Italy. Copeia *2011*, 64–71. 10.1643/CI-10-080.

60. Bannikov, A.F. (2004). The First Discovery of an Anglerfish (Teleostei, Lophiidae) in the Eocene of the Northern Caucasus. Paleontol. J. *38*, 420–425.

61. Carnevale, G., and Pietsch, T.W. (2011). Batfishes from the Eocene of Monte Bolca. Geol. Mag. *148*, 461–472. 10.1017/S0016756810000907.

62. Carnevale, G., and Pietsch, T.W. (2009). An Eocene Frogfish from Monte Bolca, Italy: The Earliest Known Skeletal Record for the Family. Palaeontology *52*, 745–752. 10.1111/j.1475-4983.2009.00874.x.

63. Carnevale, G., Pietsch, T.W., Bonde, N., Leal, M.E.C., and Marramà, G. (2020). †Neilpeartia ceratoi, gen. et sp. nov., a new frogfish from the Eocene of Bolca, Italy. J. Vertebr. Paleontol. *40*, e1778711. 10.1080/02724634.2020.1778711.

64. Pietsch, T.W. (1981). The osteology and relationships of the anglerfish genus Tetrabrachium, with comments on lophiiform classification. US Fish. Bull. *79*, 387–419.

65. Huddleston, R.W., and Takeuchi, G.T. (2006). A New Late Miocene Species of Sciaenid Fish, Based Primarily on an *in situ* Otolith from California. Bull. South. Calif. Acad. Sci. *105*, 30–42. 10.3160/0038-3872(2006)105[30:ANLMSO]2.0.CO;2.

66. Nazarkin, M.V., and Carnevale, G. (2018). A Miocene pearleye, Benthalbella praecessor, sp. nov. (Teleostei, Aulopiformes), from Sakhalin Island, Russia: the first known skeletal record for the family Scopelarchidae. J. Vertebr. Paleontol. *38*, e1511992. 10.1080/02724634.2018.1511992.

67. Nazarkin, M.V. (2021). The Structure of the Miocene Northwestern Pacific Ichthyofauna as Revealed By Two Fossil Fish Assemblages From Sakhalin Island, Russia. Paleontol. Res. *25*. 10.2517/2021PR005.

68. Rincon-Sandoval, M., Duarte-Ribeiro, E., Davis, A.M., Santaquiteria, A., Hughes, L.C., Baldwin, C.C., Soto-Torres, L., Acero P., A., Walker, H.J., Carpenter, K.E., et al. (2020). Evolutionary determinism and convergence associated with water-column transitions in marine fishes. Proc. Natl. Acad. Sci. *117*, 33396–33403. 10.1073/pnas.2006511117.

69. Ribeiro, E., Davis, A.M., Rivero-Vega, R.A., Ortí, G., and Betancur-R, R. (2018). Post-Cretaceous bursts of evolution along the benthic-pelagic axis in marine fishes. Proc. R. Soc. B Biol. Sci. *285*, 20182010. 10.1098/rspb.2018.2010.

70. Sibert, E., Friedman, M., Hull, P., Hunt, G., and Norris, R. (2018). Two pulses of morphological diversification in Pacific pelagic fishes following the Cretaceous–Palaeogene mass extinction. Proc. R. Soc. B Biol. Sci. *285*, 20181194. 10.1098/rspb.2018.1194.

76. Matschiner, M., Hanel, R., and Salzburger, W. (2011). On the Origin and Trigger of the Notothenioid Adaptive Radiation. PLoS ONE *6*, e18911. 10.1371/journal.pone.0018911.

77. Friedman, S.T., Price, S.A., Corn, K.A., Larouche, O., Martinez, C.M., and Wainwright, P.C. (2020). Body shape diversification along the benthic–pelagic axis in marine fishes. Proc. R. Soc. B Biol. Sci. *287*, 20201053. 10.1098/rspb.2020.1053.

78. Pietsch, T.W. (2009). Oceanic Anglerfishes: Extraordinary Diversity in the Deep Sea (University of California Press).
